# Proximity labeling reveals new insights into the relationships between meiotic recombination proteins in *S. cerevisiae*

**DOI:** 10.1101/2023.09.17.558147

**Authors:** Karen Voelkel-Meiman, Alex J. Poppel, Jennifer C. Liddle, Jeremy L. Balsbaugh, Amy J. MacQueen

## Abstract

Several protein ensembles facilitate MutSγ crossover recombination and the associated process of synaptonemal complex (SC) assembly during meiosis, but the physical and functional relationships between the components involved remain obscure. We have employed proximity labeling as a phenotypic tool to discern functional relationships between meiotic recombination and SC proteins in *S. cerevisiae*, and to gain deeper insight into molecular deficits of crossover-defective mutants. We find that recombination initiation (Spo11) and the Mer3 helicase are dispensable for proximity labeling of the Zip3 E3 ligase by components of the ZZS ensemble (Zip2, Zip4 and Spo16) but are required for proximity labeling of Zip3 by Msh4, consistent with the possibility that MutSγ joins Zip3 only after a specific recombination intermediate has been generated. Proximity labeling analysis of crossover-defective *zip1* mutants suggests a key shared defect is a failure to assemble an early recombination ensemble where ZZS can properly engage Zip3. We furthermore discovered that Zip3’s abundance within the meiotic cell is uniquely dependent on the presence of Zip1, and that the post-translational modification of Zip3 is promoted by most MutSγ pathway proteins but countered by Zip1. Based on this and additional data, we propose a model whereby Zip1 stabilizes a functional, unmodified form of Zip3 until intermediate steps in recombination are complete. We also find that SC structural protein Ecm11 is proximity labeled by ZZS complex proteins in a Zip4-dependent manner, but by Zip3 and Msh4, at least in part, via a distinct pathway. Finally, streptavidin pulldowns followed by mass spectrometry on eleven different proximity labeling strains uncovers shared proximity targets of MutSγ-associated proteins, some with known meiotic functions and others not yet implicated in a meiotic activity, highlighting the potential power of proximity labeling as a discovery tool.

## Introduction

During meiosis, several protein ensembles engage one another and DNA to ensure that parental genomes successfully partition into gametes. Proper orientation and segregation of homologous chromosomes (homologs) at meiosis I relies on the prior establishment of crossover recombination-based associations between them (Page and Hawley 2003); such interhomolog crossovers form through coordinately acting groups of proteins that promote homologous recombination-based repair of programmed DNA double strand breaks (DSBs) (Borner *et al*. 2023). Crosstalk between DSB repair machinery and proteins localized to the meiotic chromosome axis somehow ensures that meiotic crossovers preferentially involve non-sister chromatids of homologous chromosomes and facilitates crossover patterning such that every chromosome pair, no matter how small, receives at least one attachment. Functionally linked to meiotic recombination is the generation of a tripartite, multi-protein structure called the synaptonemal complex (SC), which assembles along the lengths of aligned homolog axes. Several of the factors that promote the coordinated processes of crossover recombination and SC assembly (synapsis) during meiosis have been identified, but how these proteins function and functionally relate to one another at the molecular level remains obscure.

In *S. cerevisiae*, many meiotic DSBs are repaired through the formation and resolution of DNA joint molecule structures (predominantly double Holliday junctions) with the help of a set of proteins collectively referred to as “ZMMs” (Lynn *et al*. 2007). ZMMs include the MutSγ (Msh4-Msh5) heterodimer, which has homology to the bacterial MutS protein family and the capacity to bind branched DNA structures (Ross-macdonald and Roeder 1994; Snowden *et al*. 2008), the Mer3 DNA helicase (Nakagawa and Ogawa 1999; Duroc *et al*. 2017), the ZZS complex consisting of Zip2, Zip4 and Spo16 (which also binds branched DNA structures *in vitro*) (De Muyt *et al*. 2018), and the meiosis-specific E3 SUMO ligase, Zip3 (Agarwal and Roeder 2000; Cheng *et al*. 2006; Serrentino *et al*.2013). The loss of any of these meiotic factors results in a dramatic reduction in Holliday junction intermediates and crossovers. When detectable, these proteins/protein complexes have been found to colocalize with one another at discrete foci on mid-meiotic prophase chromosomes, but how they spatially and functionally relate to one another during recombination processing remains obscure.

MutSγ crossovers in *S. cerevisiae* also rely on a uniquely specialized role of the SC structural component, Zip1. The central region of synaptonemal complex has two conserved structural features: Transverse filaments orient perpendicular to aligned chromosome axes and connect them along their entire lengths at a conserved ∼100 nm distance, while a distinct central element substructure exists at the midline of the SC, equidistant from each axis (Page and Hawley 2004). In *S. cerevisiae*, Zip1 is predicted to form rod-like units owing to an extensive central coiled-coil within the 875-residue protein (Dong and Roeder 2000); Zip1 units assemble in head-to-head fashion to create the transverse filament “rungs” of the SC. A distinct set of SC structural proteins, comprised of the Ecm11-Gmc2 heterodimer,comprise the “central element” substructure (Humphryes *et al*. 2013; Voelkel-meiman *et al*. 2013). MutSγ crossovers rely completely on Zip1, but removing Ecm11 or Gmc2 causes excess MutSγ crossovers (Voelkel-meiman *et al*. 2015; Voelkel-meiman *et al*. 2016). Thus, apart from its activity as an SC building block, Zip1 plays a specialized role in recombination.

Interestingly, SC assembly is triggered at recombination sites and is normally coupled to intermediate steps in the recombination process: In *zip2, zip3, zip4, spo16*, and *mer3* mutants, SC fails to assemble from recombination sites, possibly due to a failure to recruit a key SC structural building block (Zip4, for example, which binds directly to Ecm11 (Pyatnitskaya *et al*. 2022)), or a failure to achieve a particular recombination intermediate capable of triggering SC elaboration. Certain *zip1* mutants (including the null) exhibit a combined deficit in crossovers and SC similar to most *zmm* mutants, but the existence of two classes of *zip1* separation-of-function alleles – those that allow SC assembly but not MutSγ crossovers and those that allow MutSγ crossovers but not SC assembly – indicate that Zip1 promotes the two processes of recombination and SC assembly independently, and that Zip1 is likely centrally involved in the mechanism that couples the two processes (VOELKEL-MEIMAN 2018). Precisely what Zip1’s pro-crossover function entails, and how individual ZMM proteins engage with Zip1 in the coordinated processes of recombination and synapsis remain poorly understood.

In this study we use a proximity labeling approach to explore potential spatial and/or functional relationships between ZMMs and associated proteins. We created strains with transgenes encoding several different fusions between a ZMM or SC protein and TurboID, a biotin ligase engineered by directed evolution to function efficiently in yeast cells (Branon *et al*. 2018; Larochelle *et al*. 2019). We find TurboID can promote biotinylation of other proteins in meiotic cells without exogenous addition of biotin, and that it can function at terminal and internal positions within a given “bait” protein. Using two biotinylated targets that happen to be detectable on streptavidin blots, we demonstrate how proximity labeling can be used as a phenotypic tool to gain insight into hierarchical relationships between ZMMs, associated meiotic DSB repair factors, and SC proteins. This study inadvertently led us to discover that Zip3’s abundance within the meiotic cell is uniquely controlled by Zip1, and that Zip1 counters ZMM-mediated, post-translational modification of Zip3. Finally, streptavidin pull-down followed by mass spectrometry identifies many overlapping targets of eleven distinct TurboID fusion strains, some of which are known meiotic proteins and some not yet implicated in meiosis, underscoring the potential power of proximity labeling for discovering new components of meiotic chromosomal processes.

## Results

### Several meiotic recombination proteins remain functional when fused to the TurboID biotinylase

We used an engineered version of the *E coli* BirA biotinylase, TurboID (Branon *et al*. 2018), to explore proximity interactions between proteins that promote recombination in *S. cerevisiae* meiotic prophase cells. TurboID is active in yeast cells grown at 30°C (Larochelle *et al*. 2019). For most experiments we used alleles encoding C-terminal fusions between TurboID-3xMYC and various meiotic recombination proteins, including Zip2, Zip3, Zip4, Spo16, Mer3, Msh4, Msh5, Mlh3, and the synaptonemal complex (SC) proteins Ecm11 and Gmc2. For some alleles, such as *ZIP4iTurboID* and *ZIP3iTurboID*, the TurboID (but not 3xMYC) is positioned internal to the Zip4 or Zip3 polypeptide (see Methods). Apart from those used for spore viability analysis, strains evaluated in this study are homozygous for the *ndt80* null allele, to ensure that a 24 hour sporulation culture is enriched for meiocytes with abundant double Holliday junction recombination intermediates (dHJs) and SC structures (Xu *et al*. 1995; Voelkel-meiman *et al*. 2012). However, it is important to consider that a proximity labeling event may have occurred on a given target protein at any point during meiotic prophase, up to the time of harvesting the cells.

As for all epitope-tagged fusion proteins, the addition of TurboID likely compromises at least some known or unknown functions of a given bait protein. However, we observed that many *TurboID* fusion strains created for this study are capable of assembling SC, as indicated by multiple linear stretches of Zip1 coincident with Gmc2 on surface-spread meiotic chromosomes (Figure S1). Notable exceptions are the *MER3-TurboID*, *ECM11-TurboID* and *TurboID-GMC2* homozygotes, which fail to assemble SC. *MER3-TurboID* meiotic nuclei frequently show an aggregate of SC proteins referred to as a polycomplex. Polycomplexes were not observed in the other *TurboID* fusion strains created for this study. The absence of polycomplex in the SC-deficient *TurboID-GMC2* and *ECM11-TurboID* strains is consistent with the fact that not only SC but also polycomplex structures depend on the Ecm11 and Gmc2 core building blocks (Humphryes *et al*. 2013).

We furthermore observed that, while *MER3-TurboID* homozygotes display a severe sporulation defect, most of the other *TurboID* fusion strains show high spore viability. *zip2*, *zip4*, and *spo16* null mutants in our (BR) genetic background normally make very few spores (Tsubouchi *et al*. 2006; Voelkel-meiman *et al*. 2015), but *ZIP2-TurboID*, *ZIP4iTurboID*, *ZIP4-TurboID*, and *SPO16-TurboID* homozygotes each generate an abundance of spores (indistinguishable from control strains) and spores dissected from these strains are ≥88% viable (Figure S1). The *msh5* null mutant in our strain background generates spores, but with only 53% viability (n=773 tetrads), while the *MSH5-TurboID* homozygote shows 82% viability. By contrast with *msh5*, the *msh4*, *zip3*, *mlh3*, *ecm11* or *gmc2* null mutants display relatively high spore viability in the BR genetic background (71%, 81**%**, 84%, 92%, and 92%, respectively; n>700 tetrads). Although the spore viabilities calculated for *MSH4-TurboID*, *ZIP3iTurboID*, and *MLH3-TurboID* homozygotes are higher than those of the corresponding null strains (Figure S1), it is less clear for these genes that the corresponding TurboID fusions are fully functional. The fact that the viabilities of these *TurboID* strains (Figure S1) are no lower than the null mutants indicates that the fusions do not cause a strong gain-of-function meiotic defect.

Labeling with anti-MYC antibodies to detect C terminal TurboID fusion proteins revealed, in most cases, dozens of discrete foci on mid-meiotic prophase chromosomes reminiscent of the distribution profile of ZMMs (Figure S1). Taken together with the observed viability of spores and presence of SC, this result suggests that most ZMM-TurboID fusion proteins engage functionally with recombination ensembles. Ecm11-TurboID also shows punctate localization along meiotic chromosomes, likely reflecting its Zip4-mediated recruitment to recombination sites (Pyatnitskaya *et al*. 2022). Mer3- TurboID, on the other hand, is not detected on chromosomes and only detectable faintly at the polycomplex structure (Figure S1).

For this exploratory study, we evaluated proximity labeling in all of the *TurboID* strains we created, regardless of whether a particular TurboID fusion appears fully functional.

### MutSγ-associated proteins proximity label Zip3 in Zip1-dependent but Ecm11-independent manner

To identify proteins proximity labeled by recombination or SC proteins during meiotic prophase, we used streptavidin blotting. Total protein harvested from cells after 24 hours of sporulation was separated on either an 8% or a 12% polyacrylamide gel and transferred to membrane, which was then probed with streptavidin:HRP to label biotinylated species (Figure 1). As previously observed for yeast mitotic cells (Kim *et al*. 2004; Larochelle *et al*. 2019), several naturally biotinylated proteins are robustly detected in *S. cerevisiae* meiotic cells using this method (such naturally biotinylated proteins are observed in extracts from all strains including the no TurboID control; Figure 1A, yellow dots); three particularly prominent naturally biotinylated proteins migrate at ∼27 kDa, ∼42 kDa, and ∼130 kDa.

**Figure 1.**
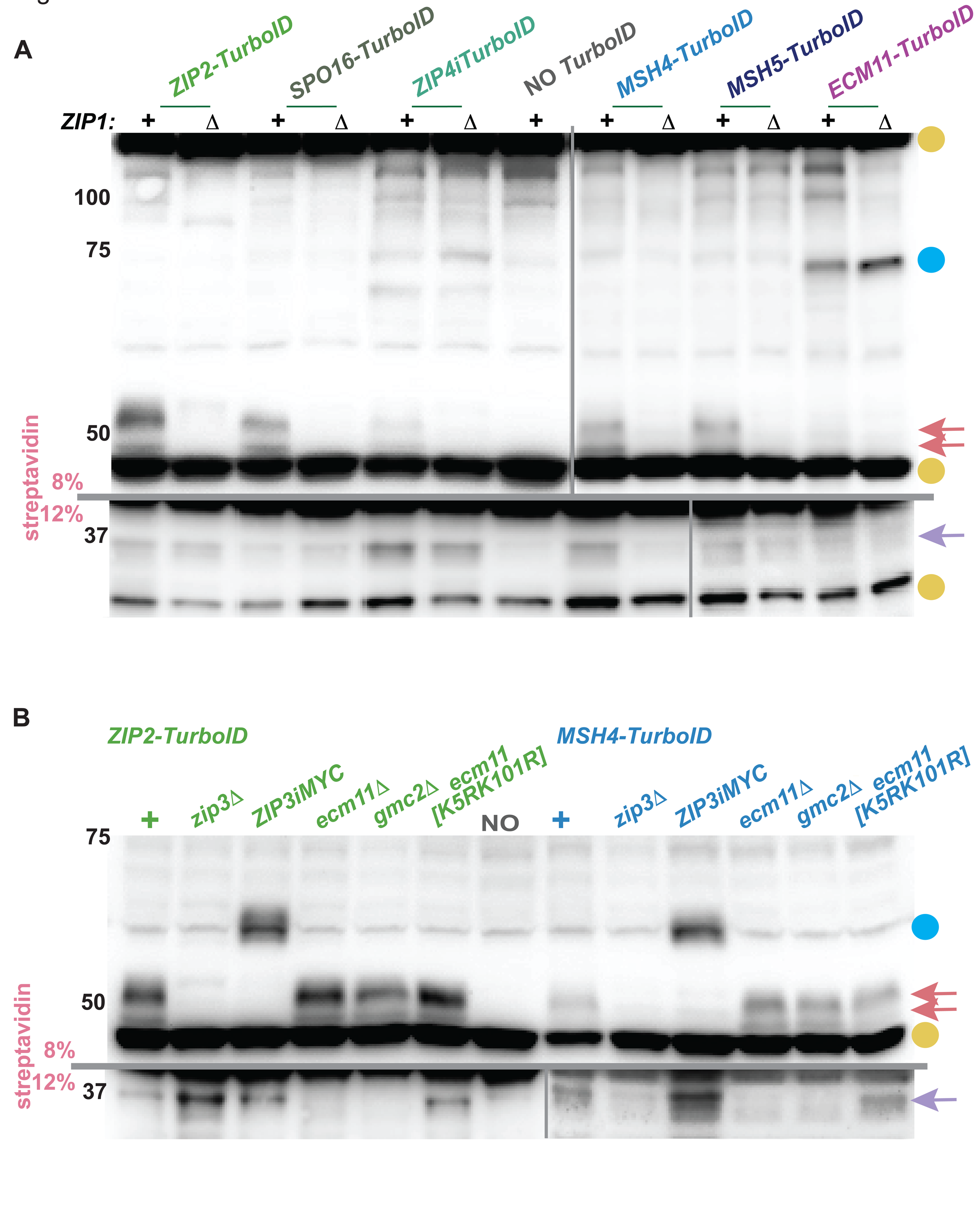
Streptavidin blots detect two proximity labeling targets of ZZS and MutSγ proteins. Proteins extracted from cells arrested at mid-meiotic prophase (strains are homozygous for *ndt80* and collected after 24 hours in sporulation medium) were separated on an 8% (above grey line) or 12% (below grey line) polyacrylamide gel and transferred to nitrocellulose, then probed with streptavidin:HRP. Blots in (A) show proteins from one of six distinct *TurboID* fusion strains, either *ZIP1+* or *zip1Δ*. The “NO *TurboID*” lane corresponds to a strain devoid of any TurboID transgene. Blots in (B) show proteins from strains homozygous for *ZIP2-TurboID* (green) or *MSH4-TurboID* (blue) and carrying various alleles of *ZIP3*, *ECM11* or *GMC2*. Gold circles at the right of blots in (A, B) indicate prominent naturally biotinylated proteins found in meiotic cells independent of TurboID. Blue circle in (A) corresponds to a new species detected in *ECM11-TurboID* strains, which is likely the Ecm11-TurboID protein itself. Arrows in (A, B) indicate the position of biotinylated proteins detected only in strains carrying a particular *TurboID* fusion gene. Pink arrows correspond to a population of biotinylated Zip3 proteins, which shift to a position of greater mass in *ZIP3iMYC* strains (blue circle at right of blot in (B)); purple arrow corresponds to a ∼37 kDa protein that is not detected in strains with tagged or null versions of *ECM11* or *GMC2*. Shown are representative data (grey vertical and horizontal lines demarcate data from independent membranes); two or more biological replicates were examined for all strains.

Only a few biotinylated factors were observed in a *TurboID* fusion strain but not in the “no TurboID” control using the streptavidin blot. One set of proximity labeled targets, ranging in size between ∼45-55 kDa (Pink arrows, Figure 1A), is specifically detected in *ZIP2-TurboID, SPO16-TurboID, MSH4- TurboID,* and *MSH5-TurboID,* but is barely detectable in *ZIP4iTurboID* and undetectable in *ECM11- TurboID*. Interestingly, proximity labeling of this group of proteins by Zip2-TurboID, Spo16-TurboID, Msh4-TurboID and Msh5-TurboID is dependent on the presence of Zip1 (Figure 1A).

We determined that the entire set of 45-55 kDa biotinylated target proteins corresponds to Zip3. In *ZIP2-TurboID* or *MSH4-TurboID* strains homozygous for *ZIP3iMYC* (which encodes Zip3 with an internal 64 residue MYC tag), the 45-55 kDa biotinylated proteins are no longer detected, while a larger species that migrates between the 50 and 75 kDa markers is observed (Figure 1B).

The Zip1-dependency raises the possibility that SC structure is required for Zip3 to be proximity labeled by Zip2-TurboID, Spo16-TurboID, Msh4-TurboID and Msh5-TurboID. However, this is not the case, as neither the Ecm11 nor Gmc2 proteins are required for Zip3 proximity labeling by *ZIP2-TurboID* or *MSH4-TurboID* (Figure 1B). Together, these data indicate that Zip2, Spo16, Msh4, and Msh5 are arranged within the meiotic prophase cell in a manner that allows the TurboID fusion version of each protein to proximity label Zip3, and that is independent of mature SC but dependent on Zip1.

### A 37 kDa protein proximity labeled by ZZS proteins and MutSγ likely corresponds to Ecm11

A second proximity labeling target of several meiotic TurboID fusions migrates near the 37 kDa marker on a 12% polyacrylamide gel (Figures 1, S2). This ∼37 kDa protein has a less robust signal on the streptavidin blot relative to Zip3, but is routinely detected in *ZIP2-TurboID, ZIP3iTurboID, ZIP4iTurboID,* and *MSH4-TurboID* strains (purple arrows, Figures 1A, 3A), and is faint but consistently detectable in *SPO16-TurboID* and *MSH5-TurboID* strains (Figure 1). The ∼37 kDa biotinylated protein is not detected in *ECM11-TurboID* strains (Figure 1A, 3A), nor in any *TurboID* strains examined that lack the Ecm11 or Gmc2 proteins (Figures 1B, S2A), but is detected in strains homozygous or heterozygous for *TurboID-GMC2* (Figure S2B). Analysis of strains homozygous for *ecm11[K5R, K101R]*, demonstrate that Ecm11 SUMOylation is dispensable for Zip2-TurboID or Msh4-TurboID to proximity label the ∼37 kDa target (Figure 1B).

The Ecm11 protein interacts directly with Zip4 (Pyatnitskaya *et al*. 2022) and its molecular weight is 34 kD, raising the possibility that the ∼37 kDa proximity target of Gmc2-, Zip2-, Spo16-, Zip4-, Zip3- and Msh4-TurboID fusions is Ecm11. This possibility is supported by the dependence of the ∼37 kDa target on Ecm11 and Gmc2, which form a heterocomplex (Humphryes *et al*. 2013). Furthermore, we observe that heterozygotes carrying one copy of *ECM11-TurboID* exhibit two specific biotinylated proteins on a streptavidin blot - one corresponding to the size expected of the Ecm11-TurboID and another robust target that migrates at the ∼37 kDa target position (Figure S2B, blue circle and purple arrow respectively). The 37 kDa target is no longer detectable in *ZIP4iTurboID* strains homozygous for *ECM11-3XFLAG* (Figure 3A) consistent with the possibility that this target corresponds to Ecm11, however the expected shifted biotinylated protein (corresponding to epitope-tagged Ecm11) is not observed. Taken together, our data support either of two distinct possibilities: i) The TurboID fusions examined here biotinylate a ∼37 kDa protein that is not Ecm11 but depends upon Ecm11 function, or ii) our TurboID fusions proximity label Ecm11, but not Ecm11-3XFLAG. We favor the latter possibility, in part because an amino acid substitution in the Zip4 polypeptide that specifically compromises a yeast two-hybrid interaction with Ecm11 (encoded by *zip4[N919Q]*; (Pyatnitskaya *et al*. 2022)) also abolishes the proximity labeling interactions between Zip2-, Zip3-, and Spo16-TurboID fusions and the∼37 kDa target protein; furthermore, the Zip4[N919Q]iTurboID protein fails to proximity label the 37 kDa target (Figure S2B). Hereafter, we refer to the ∼37 kDa proximity labeling target of our TurboID fusion strains as Ecm11.

### Proximity labeling of Zip3 by Zip2- or Spo16-TurboID fusions requires an intact ZZS ensemble and Zip1 but not recombination initiation, SC, or polycomplex structure

Cytological, ChIP, and functional data indicate that ZMM proteins and Zip1 co-localize to the same recombination sites (Agarwal and Roeder 2000; Tsubouchi *et al*. 2006; Serrentino *et al*. 2013; De muyt *et al*. 2018; Pyatnitskaya *et al*. 2022). However, ZMMs and Zip1 also localize to sites predicted to be relatively devoid of recombination intermediates, such as centromeres and the nucleolus, where polycomplexes assemble when SCs are delayed or unable to form (Tsubouchi and Roeder 2005; Tsubouchi *et al*. 2008; Macqueen and Roeder 2009). To explore whether the proximity labeling of Zip3 that we detect on streptavidin blots occurs at recombination sites, we evaluated proximity labeling outcomes of several TurboID fusions in strains missing key components of the MutSγ recombination pathway (Figures 2 and 3). A summary of all genetic dependency data reported in Figures 1-3 is illustrated in Figure 3B.

**Figure 2.**
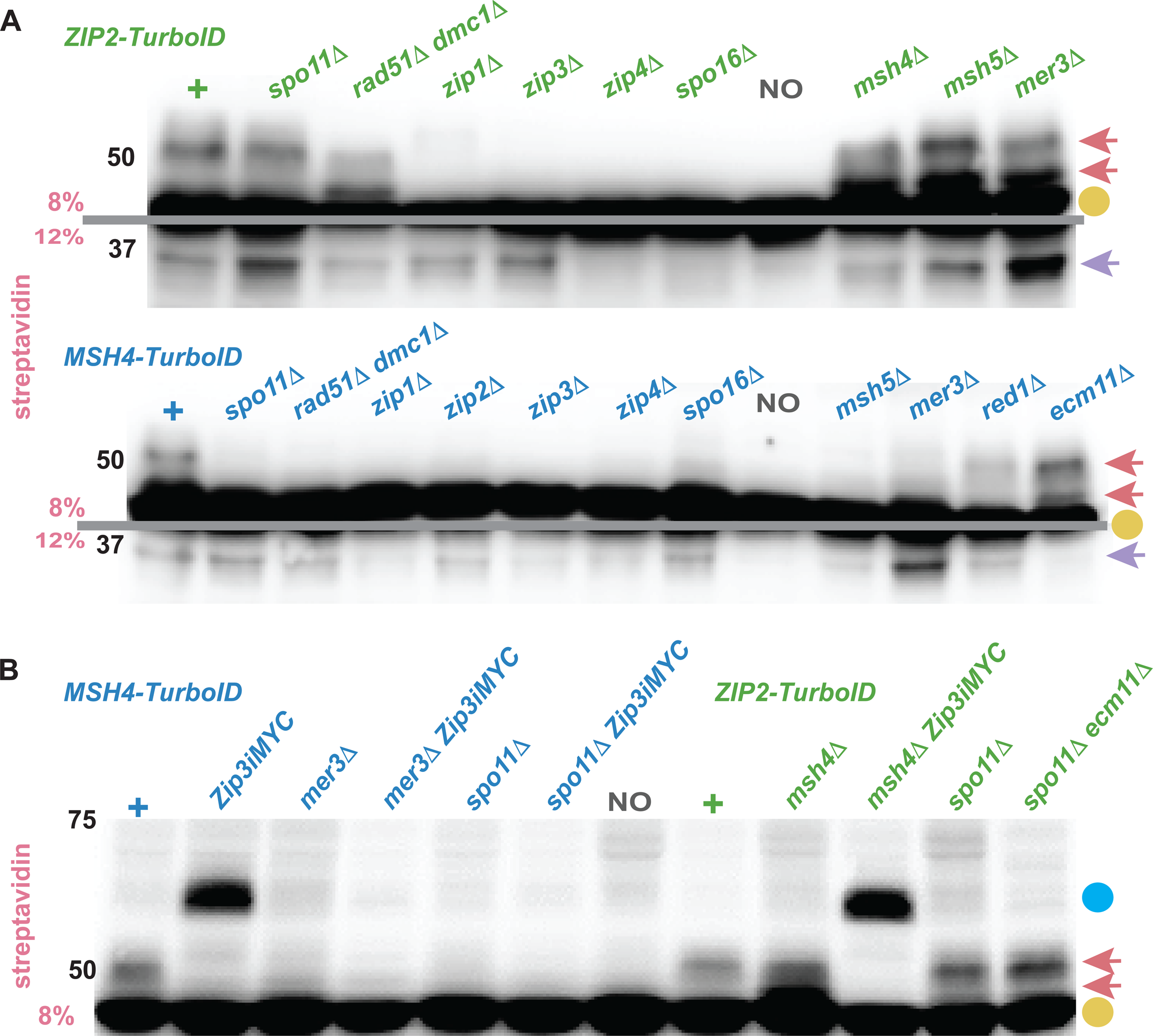
Overlapping but distinct components of the MutSγ meiotic recombination pathway are required for Zip2 and Msh4 to proximity label Zip3 and the 37 kDa protein. Blots show biotinylated proteins extracted from mid-meiotic prophase arrested (*ndt80)* cells and separated on an 8% or 12% polyacrylamide gel (above or below the grey line, respectively, in (A)). (A) Strains homozygous for *ZIP2- TurboID* (green, top blot) or *MSH4-TurboID* (blue, bottom blot) and missing the function of one of several genes required for proper MutSγ crossover recombination. Blot in (B) shows strains carrying *MSH4-TurboID* (blue) or *ZIP2-TurboID* (green) and homozygous for alleles notated across the top. Gold circles indicate naturally biotinylated proteins found in meiotic cells independent of TurboID, pink arrows correspond to biotinylated Zip3 proteins (two pink arrows are meant to highlight the presence of multiple Zip3 species that migrate at two or more positions within this area of the blot). The blue circle corresponds to biotinylated Zip3iMYC (encoding a Zip3 protein with 3xMYC inserted in-frame within the protein), while the purple arrow corresponds to the ∼37 kDa biotinylated target whose identity is likely Ecm11 (see Figures 3 and S2). Shown are representative blots; two or more biological replicates were examined for all strains. See Figure 3B for a genetic dependency chart which summarizes all genetic dependency data.

**Figure 3.**
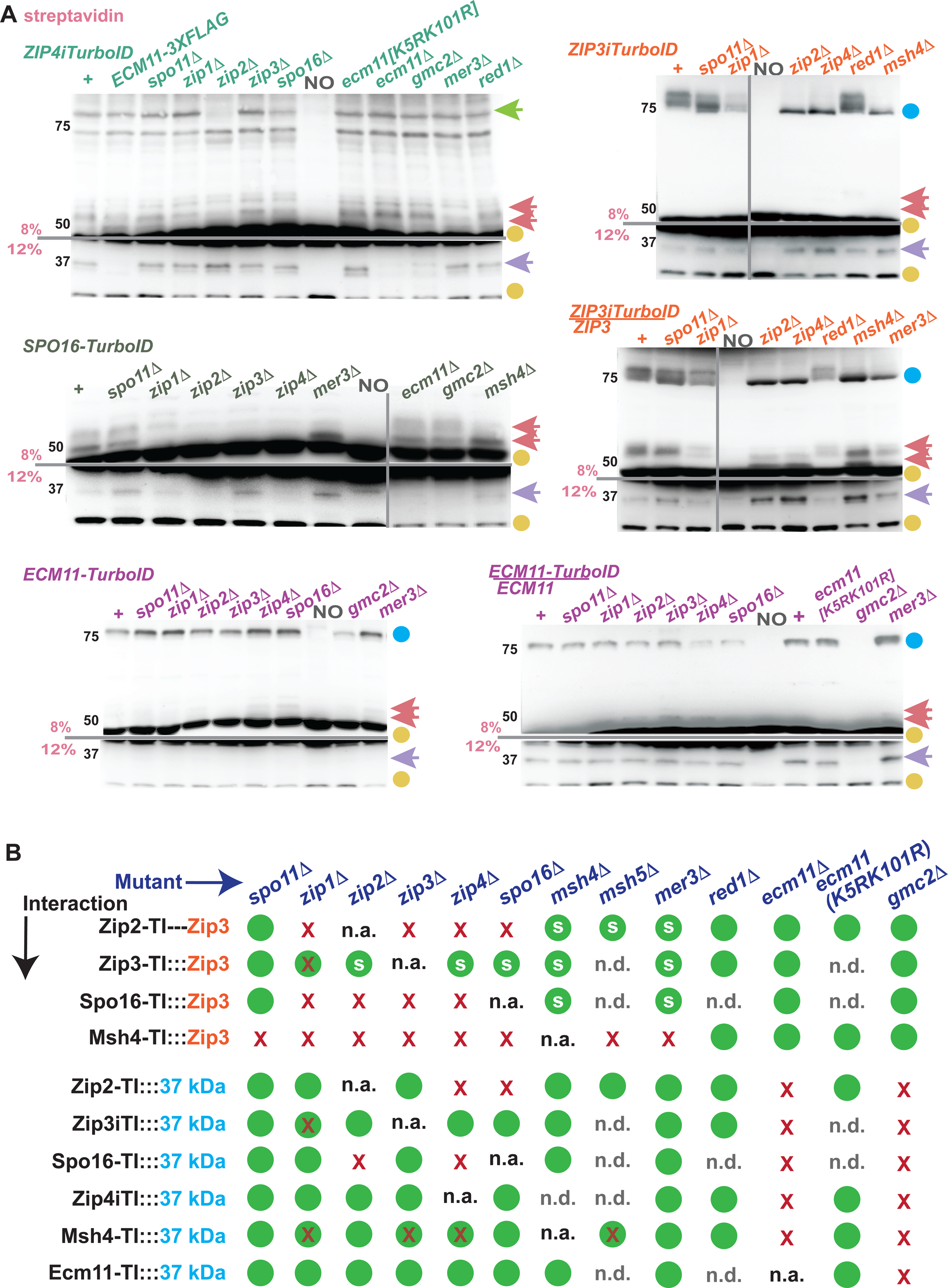
Zip3 and the 37 kDa protein are proximity labeled by several ZMM proteins. Blots in (A) display biotinylated proteins extracted from *ndt80* cells homozygous for a *TurboID* transgene, separated on an 8% (above grey lines) or 12% (below grey lines) polyacrylamide gel and visualized with streptavidin:HRP. *TurboID* fusions shown are: *ZIP4iTurboID* (green, top left; encodes an internal, in- frame fusion between Zip4 and TurboID), *SPO16-TurboID* (grey, middle left), *ECM11-TurboID* (purple, bottom left), *ZIP3iTurboID* (orange, top right; encodes an internal, in-frame, fusion between Zip3 and TurboID), *ZIP3iTurboID/ZIP3+* (orange, middle right) or *ECM11-TurboID/ECM11+* (purple, bottom right). Distinct *TurboID* strains on each blot lack the function of a gene required for proper MutSγ crossover recombination or SC assembly (mutants listed across top of blots). Gold circles indicate naturally biotinylated proteins found in meiotic cells independent of TurboID. Green arrow indicates a∼80 kDa biotinylated protein in Zip4iTurboID strains that may correspond to Zip2. Pink arrows correspond to biotinylated Zip3 proteins (two pink arrows are meant to highlight the presence of multiple Zip3 species that migrate at two or more positions within this area of the blot). Blue circles correspond to tagged protein species (either Zip3iTurboID or Ecm11-TurboID), purple arrow corresponds to the ∼37 kDa species whose identity is likely Ecm11. Note the 12% SPO16-TurboID blot displays a nonspecific biotinylated protein that is positioned (slightly above) the position of the 37 kDa target protein biotinylated by Zip4iTurboID, Spo16-TurboID, and other TurboID strains presented in this study; this background protein is occasionally detected. Shown are representative blots (grey lines demarcate independent membranes); two or more biological replicates were examined for all strains. Chart in (B) illustrates whether a proximity labeling interaction with Zip3 (top four rows) or with the 37 kDa protein (bottom six rows) is robustly detected (green circle), is less robustly detected relative to the control (red X inside a green circle), or not detected (red X) in that meiotic mutant. A green circle with an “s” indicates that the population of Zip3 species labeled appears faster migrating than the population of Zip3 biotinylated in the control. Data plotted is informed by at least two biological replicates, and multiple technical replicates; representative blots are shown in Figures 1, 2, 3 and S2). Possible Zip4iTurboID:::Zip3 and Ecm11-TurboID:::Zip3 interactions are not illustrated because the signal to noise corresponding to each of these potential interactions is too low to reliably interpret. n.a. = not applicable; n.d. = not determined.

Proximity labeling of Zip3 by ZZS complex proteins shows interdependencies that are consistent with prior biochemical data indicating Zip2, Zip4, and Spo16 form a stable subcomplex in yeast meiotic cells (De Muyt *et al*. 2018): Zip4 and Spo16 are required for Zip2-TurboID to proximity label Zip3, but Spo11, the Mer3 helicase, MutSγ (Msh4 or Msh5), the meiotic axis-associated protein Red1, and the SC structural components Ecm11 or Gmc2 are each dispensable for the Zip2-Zip3 proximity interaction (Figure 2). Similarly, Zip4 and Zip2, but not Spo11, Mer3, MutSγ, or Gmc2 are required for Spo16- TurboID to proximity label Zip3 (Figure 3). The findings are consistent with a ZZS-Zip1-Zip3 ensemble forming independent of early steps in recombination, MutSγ, and SC assembly. Unsurprisingly given the independence of the interactions on Spo11, Zip2-TurboID proximity labels Zip3 and Ecm11 even when Dmc1 and Rad51 recombinases are both absent (Figure 2).

In a recombination-defective meiotic cell, SC components aggregate to form a polycomplex, and recombination proteins decorate the polycomplex structure (e.g. Tsubouchi *et al*. 2006). We thus expected that the recombination-independent proximity labeling interactions observed between ZZS proteins and Zip3 would depend upon the existence of polycomplex structures. To our surprise, we found that Ecm11, an essential structural component of SC and polycomplex, is dispensable for the Zip2-Zip3 proximity interaction in both *SPO11+* and *spo11* null cells (Figure 2B). Thus, ZZS-Zip3 ensembles exist in meiotic cells independent of recombination initiation and independent of SC or polycomplex assembly. These data also demonstrate that Zip1’s pro-recombination activity, not its SC or polycomplex assembly function, is critical for supporting the formation of ZZS-Zip3 ensembles that allow proximity interactions to occur between Zip2, Spo16 and Zip3.

Zip3 is modified by phosphorylation (Cheng *et al*. 2006; Serrentino *et al*. 2013). We noticed that when Msh4 or Mer3 is missing, the population of Zip3 proteins proximity labeled by Zip2-TurboID or Spo16-TurboID migrate faster through the gel as if they are less modified relative to the population of Zip3 proximity labeled in control cells (see double pink arrow in Figure 2). The faster migrating species detected in these mutants correspond to Zip3 protein, as they change migration position in *ZIP2-TurboID msh4* strains homozygous for *ZIP3iMYC* (Figure 2B; see below for further discussion).

### Spo11, RecA recombinases, and Mer3 are essential for the Msh4-Zip3 proximity interaction

In contrast to the Zip2-TurboID fusion protein, Msh4-TurboID fails to proximity label Zip3 in the *spo11* mutant, or in the *dmc1 rad51* double mutant (Figure 2). Furthermore, while dispensable for proximity labeling of Zip3 by Zip2-TurboID, the Mer3 helicase is required for proximity labeling of Zip3 by Msh4- TurboID. These data together point to the possibility that the ensemble mediating proximity labeling of Zip3 by MutSγ relies on the prior formation of a recombination intermediate structure via RecA-mediated strand invasion and the Mer3 helicase.

ZZS proteins and MutSγ are normally found colocalized with Zip3 at discrete foci on mid-meiotic prophase chromosomes (Voelkel-meiman *et al*. 2019); accordingly we observe numerous discrete Zip3iMYC foci co-localized with either Zip4iHA or Msh4-MYC on surface-spread, mid-meiotic prophase chromosomes in *MER3*+ *ndt80* cells (Figure 4) We observe a reduction in the number of Zip3 foci on chromosomes in *mer3* mutants relative to the control strain, but several larger chromosome-associated Zip3 structures were observed, and Zip4 protein co-localizes with Zip3 at these structures (Figure 4, top right). By contrast, we observe hardly any Msh4 associated with meiotic chromosomes in *mer3* mutants (Figure 4, bottom right). The detection of a proximity interaction does not necessarily predict cytological detection of co-localized proteins (and *vice versa*), but our cytological observations in this case are consistent with a differential reliance on Mer3 for the formation of a Zip2-Zip3 ensemble, versus a Msh4-Zip3 ensemble, that is productive for the proximity labeling of Zip3.

**Figure 4.**
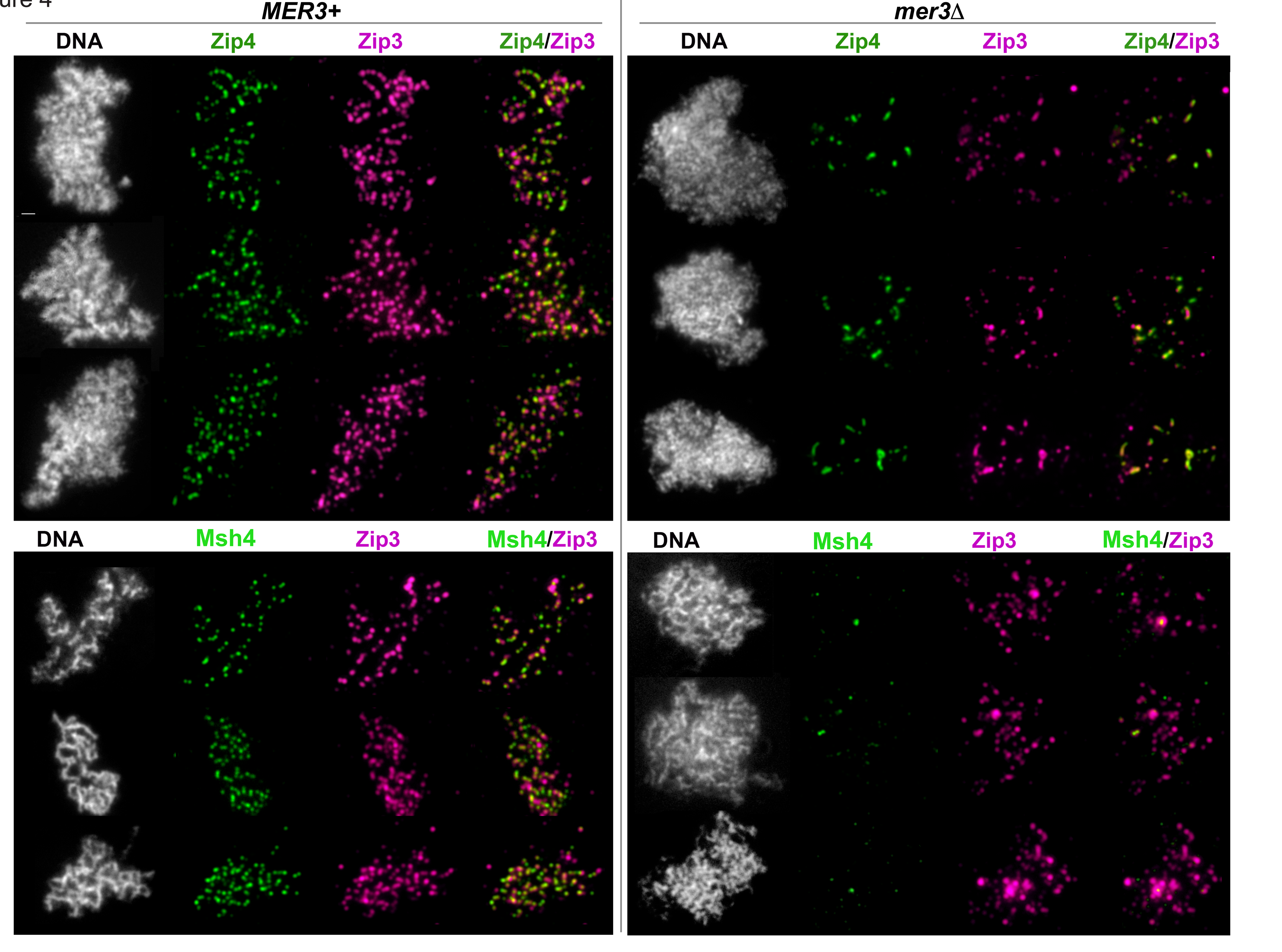
Zip3 colocalizes with ZZS protein Zip4 but not with MutSγ at chromosome-associated structures on mid-meiotic prophase chromosomes when Mer3 helicase is absent. Images show surface-spread chromosomes from strains homozygous for *ndt80* and either *MER3+* (left) or a *mer3* null allele (right). Strains prepared for top panel images carry *ZIP3iMYC* and *ZIP4iHA* alleles, while strains in bottom panel images carry *ZIP3iMYC* and *MSH4-HA* alleles. Zip3iMYC is shown in magenta, while Zip4iHA or Msh4-HA is shown in green; DAPI labels DNA (white). Three distinct nuclei for each genotype are depicted. Bar, 1 micron.

In addition to recombination initiation, strand invasion, and the Mer3 helicase, detectable proximity labeling of Zip3 by Msh4-TurboID also relies on the ZZS proteins (Figures 2 and 3). These data align with the idea that MutSγ engages a meiotic recombination intermediate after it has undergone some initial assembly.

In summary, our genetic dependency data indicate that Zip2, Zip3, Zip4, and Spo16 collaborate to form an ensemble configured such that Zip3 can be proximity labeled by Zip2 and Spo16, and that this ensemble forms independent of recombination and Ecm11 but is dependent on Zip1. On the other hand, MutSγ (Msh4-TurboID) requires recombination initiation and most all MutSγ pathway recombination proteins to proximity label Zip3.

### Zip3-TurboID proximity labels itself in *trans*

Analysis of biotinylated proteins in *ZIP3iTurboID* homozygous and heterozygous meiotic cells revealed that Zip3iTurboID can proximity label itself or a Zip3 protein in *trans* (Figure 3). (These data do not rule out nor demonstrate the existence of Zip3iTurboID *cis*-biotinylation.) We find that *trans* Zip3 labeling occurs independently of Spo11 and the core MutSγ recombination proteins (Figure 3), but the abundance of biotinylated Zip3 and Zip3iTurboID is diminished in *ZIP3iTurboID* strains missing Zip1. We also note that in *ZIP3iTurboID* homozygotes or heterozygotes lacking Zip2 or Zip4, the size of biotinylated Zip3 (both untagged and TurboID tagged) is shifted in a manner that suggests a diminishment in Zip3 post-translational modifications (double arrows in Figure 3), reminiscent of the Zip3 size shift observed in Zip2-TurboID or Spo16-TurboID strains missing a ZMM protein (reported above).

### Differential reliance on Zip1 and Zip3 distinguish the ZZS and MutSγ interactions with Ecm11

Ecm11 is proximity labeled by Zip4iTurboID even when Zip2 or Spo16 is missing (Figure 3), consistent with the existence of a direct interaction between Zip4 and Ecm11 (Pyatnitskaya *et al*. 2022).

However, Zip2 and Spo16 rely not only on Zip4 but also one another to proximity label Ecm11, suggesting Zip2 and Spo16 depend upon a fully formed ZZS complex to engage with Ecm11.

Streptavidin blots of *zip1* mutants reveal that Zip1 is dispensable for proximity labeling of Ecm11 by Zip2-TurboID, Zip4iTurboID, and Spo16-TurboID (Figure 1) indicating that a putative ZZS-Ecm11 ensemble does not require Zip1.

Interestingly, Zip1 is partly required for the proximity labeling of Ecm11 by both Msh4-TurboID and Zip3iTurboID (Figures 1, 3), suggesting the existence of a Zip1-Zip3-MutSγ-Ecm11 ensemble that is independent of an assembly carrying ZZS-Ecm11. Consistently, Zip3 is (like Zip1) dispensable for the proximity labeling of Ecm11 by Zip2-TurboID but partly required for Ecm11’s proximity labeling by Msh4-TurboID (Figure 1B).

MutSγ may engage with both the putative ZZS-Ecm11 and Zip1-Zip3-Ecm11 ensembles, as the proximity labeling of Ecm11 by Msh4-TurboID partly relies not only on Zip1 and Zip3, but also Zip4 (Figure 2, 3B). Our data point to the existence of two ensembles (or sub-ensembles) that facilitate MutSγ proximity to Ecm11: one involving Zip4 and Ecm11, and one involving Zip1, Zip3, and Ecm11.

Finally, the proximity labeling of Ecm11 by both Zip2-TurboID and Msh4-TurboID occurs even when Spo11, or the Rad51 and Dmc1 strand invasion proteins are missing (Figure 2A). Thus, not only ZZS but also MutSγ interacts with recombination and/or SC protein ensembles (in a manner that permits proximity labeling of Ecm11) independent of recombination intermediates. In the case of MutSγ, the MutSγ-Ecm11 containing ensembles that rely on Zip1 could be SC protein-based polycomplex structures that form in synapsis-defective *spo11* mutants.

### Zip3 is no longer detected as a proximity labeling target of Zip2 and Msh4 in several crossover-defective *zip1* mutants

The proximity labeling of Zip3 by TurboID-fused recombination or SC proteins is a phenotype that might be useful for understanding the deficiencies of mutants with reduced MutSγ crossovers. To explore this possibility, we examined the capacity Zip2-TurboID or Msh4-TurboID to proximity label Zip3 in an array of *zip1* or *zip3* mutant alleles.

We evaluated 10 non-null *zip1* alleles, each of which encodes a Zip1 protein with a small number of internally deleted or substituted residues within the amino terminal half of the protein (Figure 5). The regions affected by these alleles include one that carries adjacent pro-crossover and pro-synapsis domains (Zip1’s residues 1 through ∼163; (Voelkel-meiman *et al*. 2019)), and a region within the first half of Zip1’s predicted coiled-coil (residues 258 to 354).

**Figure 5.**
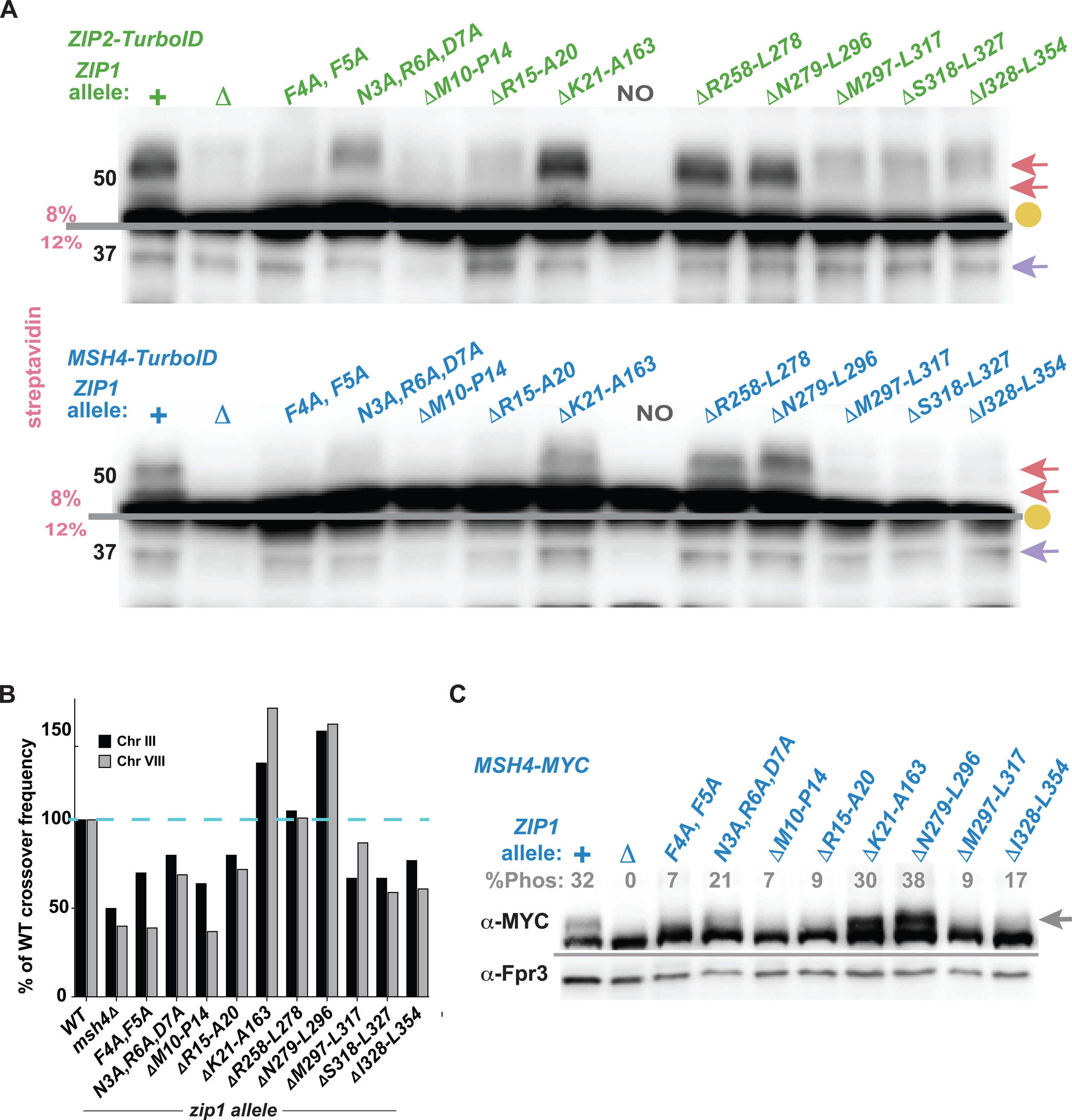
Crossover-defective zip1 mutants lack detectable proximity labeling of Zip3 by Zip2- TurboID and Msh4-TurboID. Blots display biotinylated proteins extracted from *ndt80* cells homozygous for *ZIP2-TurboID* (green) or *MSH4-TurboID* (blue) separated on an 8% (above grey lines) or 12% (below grey lines) polyacrylamide gel. *TurboID* strains are homozygous for *zip1* point mutations or in-frame deletion alleles (listed across top of blots). Gold circles indicate naturally biotinylated proteins, pink arrows correspond to biotinylated Zip3 proteins, and purple arrow indicates the biotinylated 37 kDa protein (inferred to be Ecm11). Shown are representative blots; two or more biological replicates were examined for all strains. Graph in (B) plots crossover recombination frequency on chromosomes III and VIII in each *zip1* mutant strain; four genetic intervals that span most of the length of III (black bars) and three intervals spanning over half of chromosome VIII (grey bars) are plotted as a percentage of wild type (100%, dotted blue line). Most data are calculated from more than 400 tetrads; we previously published some of the displayed crossover data (Voelkel-meiman *et al*. 2016; Voelkel-meiman *et al*. 2019; VOELKEL-MEIMAN 2021). See Table S2 for raw data. Western blot in (C) displays Msh4-MYC proteins from *ndt80* strains homozygous for *zip1* alleles; phosphorylated Msh4-MYC protein appears as a slower migrating species indicated by the grey arrow. Percentage of total Msh4-MYC that corresponds to the phosphorylated version is indicated above each lane (grey); values given are an average of two biological replicates with the following standard deviations: *+*=1.2; *zip111*=0; *zip1[F4A, F5A]*=0.6; *zip1[N3A, R6A, D7A]*=4.4; *zip1[Δ10-14]*=0.1; *zip1[Δ15-20]*=1.7; *zip1[Δ21-163]*=5.0; *zip1[Δ279-296]*=4.5; *zip1[Δ297-317]=*2.9; *zip1[Δ328-354]*=1.6. Fpr3 (shown below grey line) was utilized as a loading control.

We find that Zip3 proximity labeling by Zip2-TurboID and by Msh4-TurboID correlates with a capacity for each *zip1* mutant to generate crossovers. For example, Zip2-TurboID and Msh4-TurboID proximity label Zip3 robustly in *zip1[Δ21-163], zip1[Δ258-278],* and *zip1[Δ279-296]* strains (Figure 5A), and crossovers are at wild type levels or higher in these strains (Figure 5B). However, in strains homozygous for *zip1[F4A, F5A]*, and *zip1[Δ10-14]*, proximity labeling of Zip3 is not detected by Zip2- TurboID nor Msh4-TurboID (Figure 5A), and these *zip1* alleles have the strongest diminishment in crossovers, as measured genetically across seven intervals on chromosomes III and VIII, (Figure 5B and Table S2). Meiotic crossovers in *zip1[N3A, R6A, D7A], zip1[Δ15-20], zip1[Δ297-317]*, *zip1[Δ318-327]*, and *zip1[Δ328-354]* homozygotes are at intermediate levels between the level found in *msh4* null strains and wild-type, and in these strains Zip3 proximity labeling by Zip2-TurboID is barely detected, while the Msh4-TurboID proximity labeling of Zip3 is not detected aside from a diminished but detectable signal in *zip1[N3A, R6A, D7A]* (Figures 5A, B). Altogether these data suggest that the role of Zip1 (at least the Zip1 residues investigated here) in promoting crossovers may be to facilitate an interaction between Zip3 and the ZZS protein-containing ensemble.

### Zip3 is proximity labeled by Zip2 but not Msh4 in some *zip3* mutant strains

A similar analysis of Zip3 proximity labeling was performed for nine non-null *zip3* alleles (Figure 6). These *zip3* alleles disrupt residues across the entire length of Zip3, including the unstructured first forty residues, a conserved isoleucine within the SUMO Interaction Motif (SIM) adjacent to the RING domain (I96; (Cheng *et al*. 2006; Serrentino *et al*. 2013)), a predicted coil region just downstream of the RING domain, and parts of the unstructured C terminal half of the protein.

**Figure 6.**
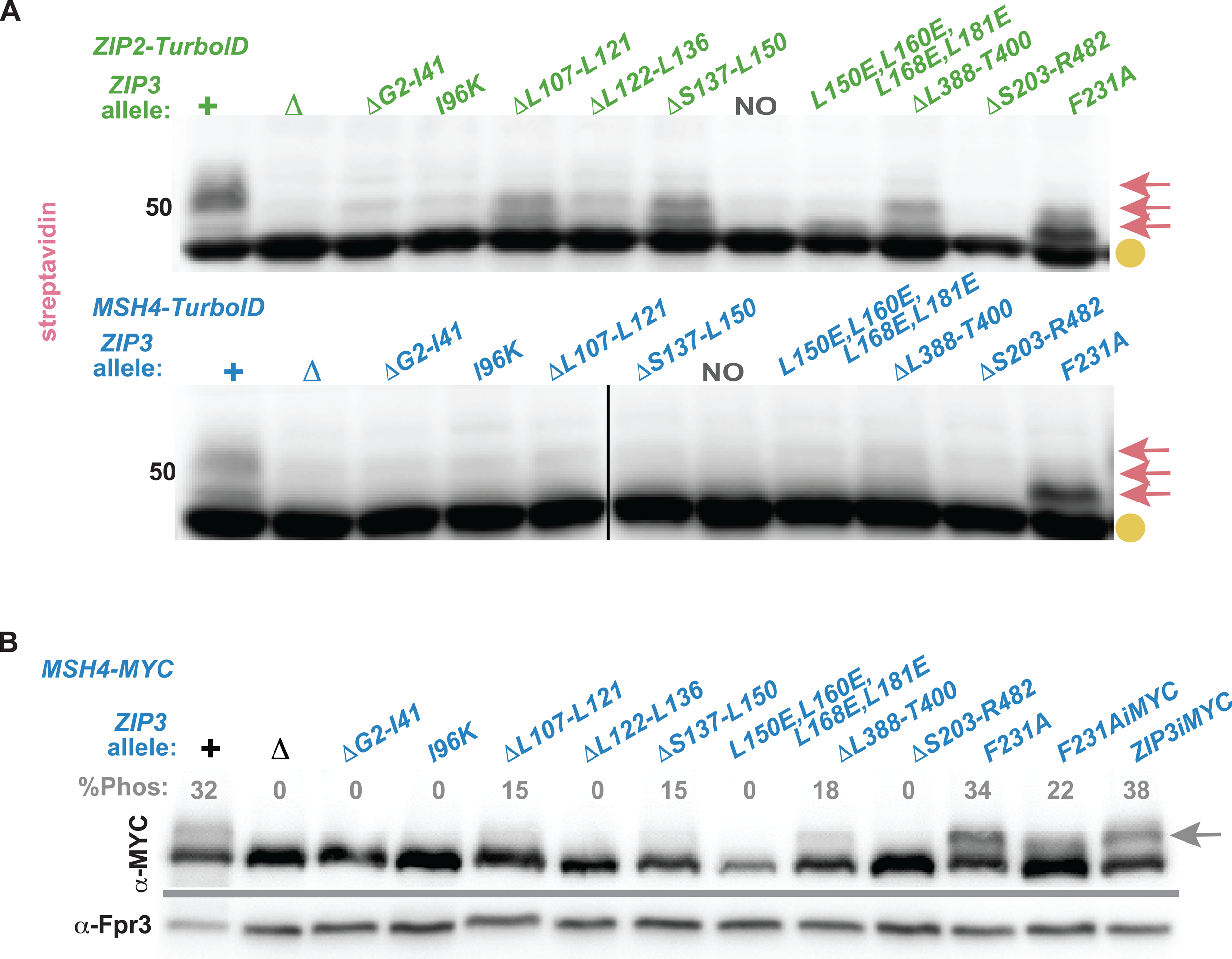
In some putative crossover-defective *zip3* mutants, Zip3 is not detectable as a proximity labeling target of Msh4 but remains a target of Zip2. Blots display biotinylated proteins extracted from *ndt80* cells homozygous for *ZIP2-TurboID* (green) or *MSH4-TurboID* (blue) arrested at mid-meiotic prophase and separated on an 8% polyacrylamide gel. *TurboID* strains are homozygous for a *zip3* point mutant or in-frame deletion allele (listed across top of blots). Gold circles indicate naturally biotinylated proteins, pink arrows correspond to biotinylated Zip3 proteins. Shown are representative blots; two or more biological replicates were examined for all strains. Western blot in (B) shows Msh4-MYC protein in *ndt80* strains homozygous for certain *zip3* alleles at mid-meiotic prophase; a reduction in phosphorylated Msh4-MYC may indicate a deficit in MutSγ crossover recombination (He *et al*. 2020 and Figure 5).Percentage of total Msh4-MYC that corresponds to the phosphorylated version is indicated above each lane (grey); values given are an average of two biological replicates with the following standard deviations: +=5.3; *zip311, zip3[Δ 2-41], zip3[I96K]=0; zip3[Δ107-121]=1.1; zip3[Δ122-136]=0; zip3[Δ137-150]=0.8; zip3[L150E,L160E,L168E,L181E]=0; zip3[Δ388-400]=1.7; zip3[Δ203-482]=0*;*zip3[F231A]*=6; *ZIP3[F231A]iMYC*=2.3; *ZIP3iMYC*=0.3. Fpr3 (shown below grey line) was utilized as a loading control.

We found that Zip2-TurboID and Msh4-TurboID both fail to proximity label Zip3 in four of the nine mutants: *zip3[Δ2-41]*, *zip3[I96K], zip3[Δ122-136],* and *zip3[Δ203-482]*. These *zip3* mutants thus may fail to assemble an early recombinosome ensemble with Zip1 and the ZZS proteins. Consistently, the*zip3[I96K]* mutant was previously found to be deficient in Zip3 function and localization to chromosomes (Cheng *et al*. 2006; Serrentino *et al*. 2013). Furthermore, our *zip1* alleles (Figure 5C) and *zmm* mutants (He *et al*. 2020; Figure S3) indicate that phosphorylated Msh4 correlates with successful MutSγ crossover recombination, and the *zip3[Δ2-41]*, *zip3[I96K], zip3[Δ122-136],* and *zip3[Δ203-482]* mutants do not accumulate normal levels of phosphorylated Msh4 during a meiotic prophase arrest (Figure 6B);.

For another group of *zip3* mutants, including the *zip3[Δ107-121], zip3[Δ137-150], zip3[L150E, L160E, L168E, L181E],* and *zip3[Δ388-400]* alleles, Zip2-TurboID proximity labels Zip3 robustly but Msh4-TurboID does not (Figure 6A). (We note that while Zip2-TurboID proximity labels an abundance of Zip3 in these mutants, much of the biotinylated Zip3 target population is faster migrating relative to the target population found in *ZIP3+* cells, reminiscent of the effect of Mer3 or Msh4 removal, described above; this is particularly true for the *zip3[L150E, L160E, L168E, L181E]* mutant.) We propose that in these *zip3* strains, a ZZS-Zip3-Zip1 ensemble may form but is deficient, at least to some degree, in maturing into a recombination complex that MutSγ can properly engage. Consistent with this idea, these *zip3* mutants accumulate sub-normal levels of Msh4 phosphorylation in meiotic prophase-arrested cells (Figure 6B), suggesting they each are defective in generating MutSγ crossovers.

Both Zip2-TurboID and Msh4-TurboID show robust proximity labeling of Zip3 in only one of the *zip3* alleles we tested, the *zip3[F231A]* mutant. *zip3[F231A]* is also unique among the nine mutants examined in supporting normal accumulation of Msh4 phosphorylation during prophase arrest (Figure 6B), suggesting that MutSγ crossovers are generated. Indeed, genetic analysis of meiotic crossovers in *zip3[F231A]* and *zip3[F231A] msh4* double mutant meiotic cells reveals that *zip3[F231A]* cells generate nearly normal levels of MutSγ crossovers (100% of the wild-type level on chromosome *III*, and 91% of wild type on chromosome *VIII*; Figure S6C, Table S2). However, in the *zip3[F231A]* strain we again observe that the biotinylated Zip3 target population is faster migrating relative to the target population found in *ZIP3+* cells.

Based on these data, our *zip3* alleles represent distinct classes of Zip3 proteins: Some fail to support the formation of an ensemble that permits proximity labeling by Zip2-TurboID, while others can assemble a ZZS-Zip3 ensemble that permits proximity labeling by Zip2-TurboID but cannot generate the ensemble that permits proximity labeling by Msh4-TurboID. The fact that no mutants were found to strong proximity labeling of Zip3 by Msh4-TurboID but not by Zip2-TurboID is consistent with the idea that formation of the ZZS protein-containing recombination ensemble is a prerequisite for the subsequent formation of a MutSγ-containing ensemble.

Finally, the *zip3[F231A]* allele supports the formation of both types of ensembles - one that enables proximity labeling by Zip2-TurboID and another that allows proximity labeling by Msh4-TurboID - however the Zip3 that is proximity labeled lacks post-translational modification. Altogether, the data indicate that successful MutSγ crossover recombination is correlated with the capacity for both ZZS and MutSγ to proximity label Zip3, but not necessarily correlated with a “normal” profile of Zip3 post-translational modification.

### Zip3’s abundance in meiotic prophase cells is uniquely controlled by Zip1

Detectable proximity labeling of Zip3 by either ZZS or MutSγ relies on Zip1. Zip1 might promote the structural integrity of each ensemble; an alternative, not mutually exclusive, possibility is that Zip3’s abundance within the cell relies on Zip1.

We used western blots to evaluate the abundance of Zip3iMYC protein in meiotic prophase (*ndt80* arrested) cells missing Spo11, Zip1, Zip2, Zip4, Spo16, Mer3, Msh4, Msh5, Red1, Ecm11, Gmc2, or homozygous for the *zip4[N919Q]* allele (which encodes a Zip4 protein that fails to interact with Ecm11; (Pyatnitskaya *et al*. 2022)) (Figure 7). Zip3iMYC displays close to wild-type abundance in most of the meiotic mutants examined. To our surprise, however, we found that Zip3iMYC accumulates to only 10- 20% of the wild-type level in *zip1 ndt80* mutants (Figure 7A, C).

**Figure 7.**
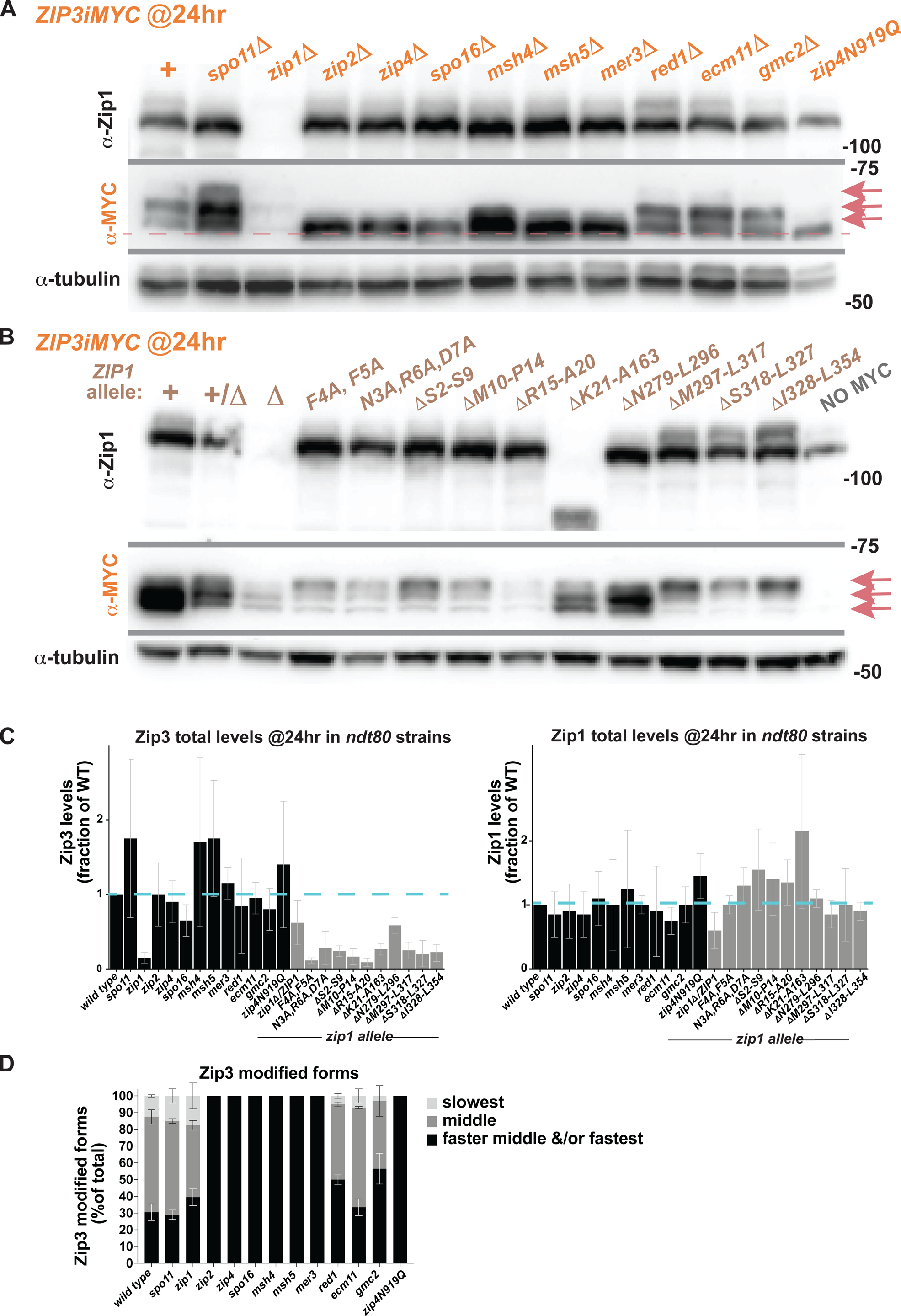
Zip3 abundance relies uniquely on the presence of Zip1, and its post-translational modification is promoted by ZMMs. Proteins extracted from meiotic prophase cells of strains homozygous for *ndt80* and *ZIP3iMYC* were separated on an 8% polyacrylamide gel and transferred to nitrocellulose, which was then sequentially probed with anti-MYC, anti-tubulin, and anti-Zip1 antibodies. Blot in (A) displays protein from various meiotic mutant strains while blot in (B) shows strains homozygous for a *zip1* non-null allele (alleles listed above the blots). Pink arrows in (A, B) indicate Zip3iMYC proteins, which consists of several species of distinct sizes, depending on the mutant background. Graphs in (C) plot the levels of total Zip3iMYC and Zip1 protein in each strain relative to the *ZIP3iMYC* control strain, utilizing tubulin as a loading control (dotted blue bar indicates the level of Zip3iMYC or Zip1 detected in *ZIP3iMYC*, which is set to one). Stacked bars in (D) indicate the relative abundance of different sized forms of Zip3 in various meiotic mutants. Dark shading refers to the fraction of total Zip3 protein migrating fastest (note that in *zmm* mutants this category may include two distinct forms of Zip3 – unmodified and minimally modified – but these forms could not be independently measured in a reliable manner), while grey shading and light grey shading indicates the fraction of total Zip3 found in the slower and slowest positions on the blot, respectively. Two biological replicates were used to evaluate protein levels, and bars indicate standard deviations for measurements in (C, D).

We asked whether Zip1 uniquely influences the abundance of other MutSγ pathway proteins (in *ndt80*, mid-meiotic prophase cells) by first examining the level of Msh4 protein in *zip1, spo11, zip2, zip3, zip4, spo16, msh5, mer3, ecm11* and *gmc2* mutants homozygous for *ndt80* and an epitope-tagged version of Msh4 (Figure S3). We found a diminishment in Msh4 levels in all the mutants examined, apart from *ecm11* and *gmc2.* However, *zip1* mutants did not show a more substantial diminishment in Msh4 relative to the other *zmm* mutants. We furthermore observed no change in the abundance of epitope-tagged Zip4 in meiotic cells missing Zip1 (Figure S3C). We conclude that Zip1 is uniquely required for maintaining the abundance of Zip3 within the *ndt80* meiotic cell. A time course analysis of Zip3iMYC abundance in suggests that Zip1 is required to maintain Zip3 abundance from the earliest stages of meiotic prophase in *ndt80* cells, not just at the 24 hour time point (Figure S4).

We note that the phosphorylated form of Msh4 is missing in *spo11* and every *zmm* mutant, but present in the MutSγ crossover-proficient *ecm11* and *gmc2* mutants (Figure S3A). This observation bolsters confidence in using Msh4 phosphorylation as a tool for preliminary analysis of whether a given mutant is proficient for MutSγ crossover recombination.

Are Zip1 and Zip3 interdependent for sustaining their levels in the meiotic cell? A *zip3* null was among the strains we evaluated for Msh4-MYC levels (Figure S3A), and we routinely probe our western blots with anti-Zip1 to ensure that the strains entered meiosis properly. In this experiment we observed that the level of Zip1 in the *zmm* mutants appeared slightly diminished relative to wild type, perhaps due to a reduction in SC structures, which may protect the Zip1 protein from degradation. However, we observe that Zip1 is as abundant in *zip3* as it is in the other *zmm* mutants (Figure S3). Thus, Zip3 relies on Zip1 to maintain its abundance in the meiotic prophase cell, but Zip1 does not rely on Zip3 for its stability in the same context.

Given the severe diminishment in Zip3 abundance when Zip1 is absent, the dependence of Zip3 proximity labeling on Zip1 may be due to an inability to detect intact proximity labeling events on a streptavidin blot simply because there are too few Zip3 proteins in the cell. While this is an important consideration, a close look at Zip3iMYC levels in *zip1* non-null mutants reveals that a similar low level of Zip3 can have positive or negative proximity labeling outcomes on a streptavidin blot (Figure 7B, C): For example, Zip3 levels are at ∼10%-30% of wild-type in several *zip1* mutants that abolish or severely diminish proximity labeling of Zip3 by Zip2-TurboID and Msh4-TurboID, such as *zip1[F4A, F5A], zip1[Δ10-14]*, *zip1[Δ297-317], zip1[Δ318-327],* and *zip1[Δ328-354]*. However, Zip3 levels are similarly low (∼12%-44%) in the *zip1[N3A, R6A, D7A]* mutant where proximity labeling of Zip3 by Zip2-TurboID is readily observed (Figure 5). Most strikingly, Zip3 is only at 20%-30% of the wild-type level in the *zip1[Δ21-163]* mutant where SC fails to assemble but MutSγ crossovers form in excess (VOELKEL-MEIMAN *et al*. 2016; VOELKEL-MEIMAN 2021), but proximity labeling of Zip3 by either Zip2-TurboID or Msh4-TurboID in this strain appears robust (Figures 5, 7).

From these data, we conclude that Zip1 not only controls the abundance of Zip3 within cells, but also facilitates Zip3 localization to and/or within ZZS and MutSγ-containing recombination ensembles. In the crossover-proficient *zip1[Δ21-163]* mutant, overall Zip3 levels are low, but residual Zip3 is properly positioned to generate MutSγ recombination events. For crossover-deficient alleles such as *zip1[Δ10-14]*, Zip3 levels are similarly low, but a second critical defect appears to be a failure of Zip3 to engage properly with ZZS- and MutSγ-containing recombination ensembles.

To determine whether severely diminished Zip3 protein levels might account for the apparent proximity labeling deficiencies of the *zip3* mutants examined, strains carrying epitope-tagged versions of many of these *zip3* alleles were examined by western blot. This analysis revealed that some of the alleles, including *zip3[Δ2-41]*, *zip3[I96K]*, and *zip3[Δ122-136]*, encode an unstable Zip3 protein (Figure S5). We found that Zip3[F231A] and Zip3[Δ388-400] proteins appear as abundant or more abundant than the control Zip3iMYC protein, but that Zip3[Δ107-121] and Zip3[L150E, L160E, L168E, L181E] are less than half as abundant as the control. We conclude that the failure of Zip2 and Msh4 to proximity label Zip3[Δ2-41], Zip3[I96K], or Zip3[Δ122-136] could be due to a severe instability of the altered Zip3 protein itself. However, we note that for other altered versions of Zip3, such as Zip3[Δ107-121], Zip3[Δ137-150], Zip3[L150E, L160E, L168E, L181E], and Zip3[Δ388-400], proximity labeling by Zip2- TurboID is easily detected, regardless of the lower Zip3 levels in the cell. Thus, residues altered in these latter mutants may be functionally important for the successful transition of an early recombination intermediate (involving Zip2 and Zip3) to a later intermediate that MutSγ can engage.

### ZMMs promote while Zip1 counters Zip3 post-translational modification

As we observed for biotinylated Zip3 in many of our proximity labeling strains, the entire Zip3iMYC population in otherwise wild type cells consists of at least three proteins migrating with distinct masses on an 8% polyacrylamide gel (Figures 7, 8); slower migrating forms of Zip3 have previously been observed and likely correspond to different phosphorylated forms (Cheng *et al*. 2006; Serrentino *et al*. 2013). *zip1* null mutants display at least three Zip3 forms as in wild type, albeit at severely diminished levels (Figures 7, 8). However, when a ZMM protein is missing, the slowest migrating forms of Zip3 found in wild type are not detected; instead, the Zip3 observed in *zip2*, *zip4*, *spo16*, *msh4*, *msh5* or *mer3* mutants most closely matches the fastest-migrating form detected in wild type (Figure 7), and an even faster migrating Zip3 is discernable when reduced amounts of input are applied to reduce signal on the blot (see lowest dotted line in Figure 8). These data explain why the population of biotinylated Zip3 proteins appears faster migrating in many TurboID fusion strains with disabled *ZMM* function, and indicate that, in *ndt80* cells, MutSγ pathway proteins promote the post-translational modification (PTM) of Zip3.

**Figure 8.**
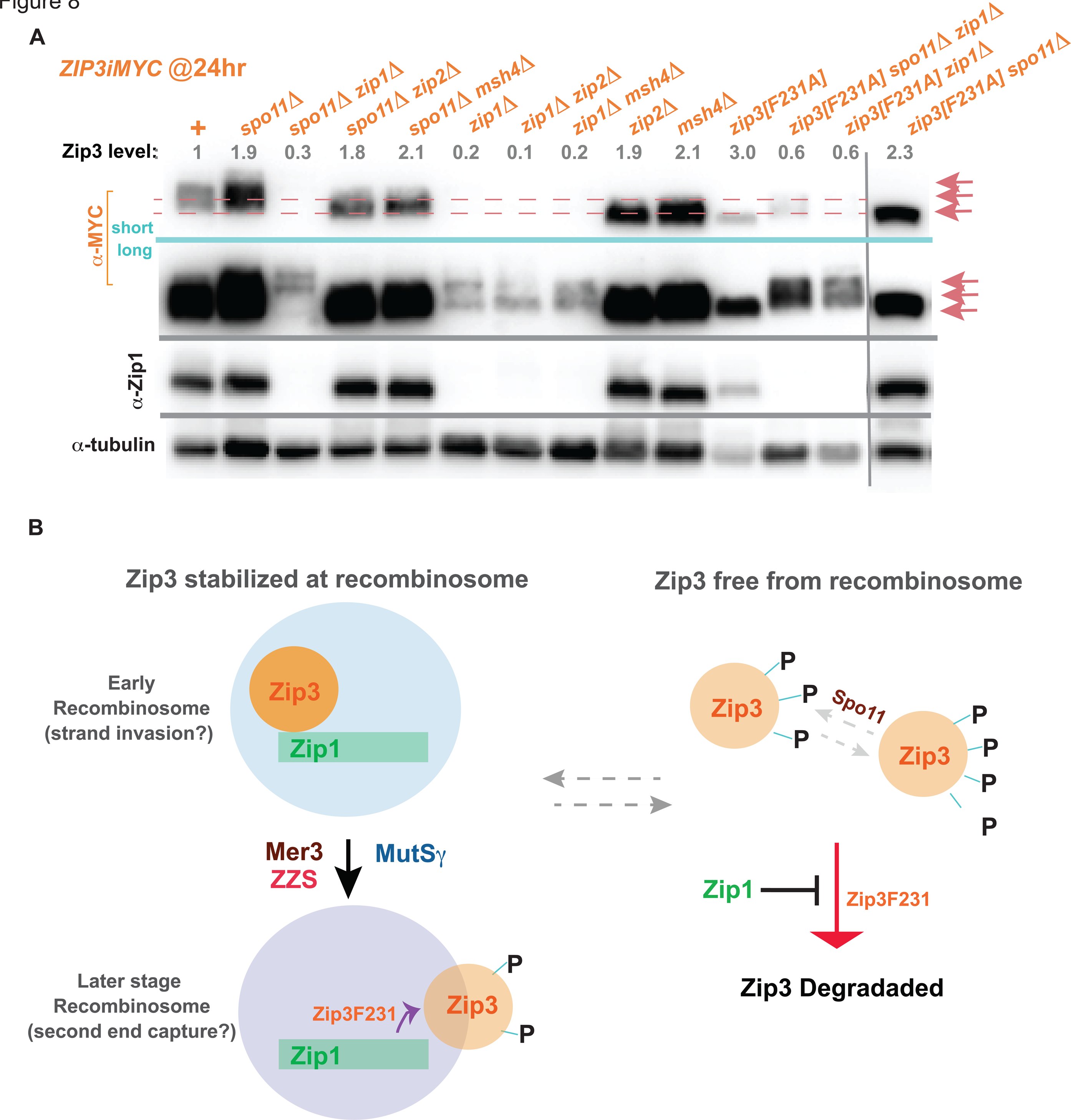
Zip3 post-translational modification is reduced when ZMM proteins are absent, so long as Zip1 is present. Blots in (A) show proteins extracted from mid-meiotic prophase cells of *ZIP3iMYC ndt80* strains homozygous for an additional meiotic mutation (listed across top of blot), separated on an 8% polyacrylamide gel. Blots were probed sequentially for anti-MYC, anti-tubulin, and anti-Zip1 antibodies. Images above and below the turquoise line show different exposure times for anti-MYC. Vertical grey line indicates independent membranes. Pink arrows indicate Zip3iMYC proteins, which comprise several species of distinct sizes depending on the presence of post-translational modifications (PTMs). Certain samples were underloaded to maximize clarity of signal; total Zip3 levels in each strain were evaluated using anti-tubulin as a loading control. Zip3 abundance (relative to the control, which is set to one) is indicated above each lane in grey; values given are an average of four experiments comprising two technical and two biological replicates, with the following standard deviations: *+*=0; *spo1111*=1.1; *spo1111 zip111*=0.3; *spo1111 zip211*=1.1; *spo1111 msh411*=1.2; *zip111*=0.1; *zip111 zip211*=0.1; *zip111 msh411*=0.04; *zip211*=1.2; *msh411*=0.8; *zip3[F231A]*=2.2; *spo1111 zip111 zip3[F231A]*=0.2; *zip111 zip3[F231A]*=0.1; *spo1111 zip3[F231A]*=1.4. Illustration in (B) suggests one interpretation of the data presented in Figures 7, 8, S4 and S5: In this model, unmodified or undermodified Zip3 is functional and stabilized by Zip1 at early ensembles of recombination proteins (possibly corresponding to the strand invasion stage). Successful completion of intermediate steps in recombination, mediated by ZMMs, leads to a release or change in configuration of Zip3 and a capacity for Zip3 to be post-translationally modified (likely phosphorylation (Serrentino *et al*. 2013)); potentially Zip3 PTMs lead to an increased likelihood of degradation at a certain meiotic stage. When Zip1 is present in the cell, Zip3’s capacity to transition from an unmodified to a modified form relies on Zip3’s phenylalanine 231, but this phenylalanine is not critical for Zip3’s pro-crossover function (Figure S6). The slowest migrating forms of Zip3 observed in *spo11* and *spo11 zip1* double mutants (A) indicate that Spo11 activity counters, to some extent, hypermodification of Zip3. Finally, the Zip3[F231A] protein appears partially resistant to degradation even when Zip1 is absent (see levels of Zip3 in (A) and Figure S5), implicating phenylalanine 231 not only in the mechanism that promotes PTMs but also in the Zip3 degradation that Zip1 counters.

In what context do MutSγ pathway proteins promote Zip3 PTMs? We found that removal of SC proteins Ecm11 or Gmc2 does not affect Zip3 PTMs in an obvious manner, nor does removal of Spo11 or the axis-associated protein Red1 (Figure 7). Our observation that Zip3 PTMs occur independent of Spo11 activity directly contrasts the conclusion of (Cheng *et al*. 2006); the different results may be due to strain background differences. We observe that *spo11 zip2* and *spo11 msh4* double mutants do not precisely phenocopy the *spo11* single mutant nor the *zip2* or *msh4* single mutants: These double mutants exhibit predominantly faster migrating Zip3 species analogous to *zip2* or *msh4* single mutants, but also show a low level of slower migrating Zip3 like the slower form detected only in wild type or *spo11* (Figure 8).

These data suggest that recombination initiation is a partial prerequisite for maintaining undermodified Zip3 when a ZMM protein is missing.

Interestingly, the Zip3[F231A] alteration causes a dramatic reduction of modified Zip3 protein in the meiotic cell. First, the population of biotinylated Zip3[F231A] protein in *ZIP2-TurboID* and *MSH4- TurboID* strains is mostly fast-migrating (Figure 6A). Second, Zip3[F231A]iMYC migrates as a single species at the position of the fastest migrating Zip3iMYC species detected in any *zmm* mutant, whether Spo11 is present or not (Figure 8). Thus, Zip3[F231A] protein may correspond to a completely unmodified form, and this result raises the possibility that Zip3’s phenylalanine 231, a conserved residue, is critical for the ZMM-mediated mechanism that promotes Zip3 PTMs.

Double mutant data also indicates that Zip1 activity counters Zip3 PTMs or stabilizes undermodified Zip3 in *zip3[F231A]* and in *zmm* mutants. In the case of *zmm zip1* double mutants such as *zip2 zip1* or *msh4 zip1*, Zip3 levels are severely diminished but the size profile of residual Zip3 resembles *zip1* single mutants where slower migrating forms are apparent, instead of the size profile of Zip3 found in the *zip2*or *msh4* single mutant (Figure 8). From these data we conclude that when a ZMM protein is absent, Zip1 blocks PTMs from being added to Zip3 (in addition to preventing Zip3 degradation) and/or selectively stabilizes unmodified Zip3.

Interestingly, the under-modification of Zip3[F231A] protein in *ndt80* cells is also dependent on Zip1: Like *zmm zip1* double mutants, the *zip1 zip3[F231A]* strain displays slower migrating forms of Zip3[F231A] (Figure 8A, B). Thus, like ZMM proteins, Zip3’s phenylalanine 231 is required for PTM addition in otherwise normal cells and is not required for Zip1 to prevent Zip3 modification. However, compared to Zip3 protein in *zmm zip1* double mutants, the Zip3[F231A] protein appears less susceptible to degradation when Zip1 is removed (Figure 8). Zip3’s phenylalanine 231 thus is also required for the mechanism that efficiently degrades Zip3 when Zip1 is missing. Taken together, our observations suggest that ZMM activity during meiotic prophase promotes Zip3 modification through a mechanism that requires Zip3’s phenylalanine 231, and that Zip1 promotes PTM removal and/or blocks PTM addition to Zip3 when a functional component of the recombination ensemble is missing, and finally that Zip1 protects Zip3 from degradation through a mechanism that also involves Zip3’s phenylalanine 231.

How might ZMM-mediated, post-translational modifications that are countered by Zip1 be related to Zip3 function? *zip3[F231A]* mutant cells are only mildly deficient in crossovers and SC assembly (Figure S6), consistent with the idea that unmodified Zip3 is functional. Also consistent with this idea is the *zip3[4AQ]* mutant, which also produces an undermodified Zip3 and shows only mild defects in crossover recombination (Serrentino *et al*. 2013). Since the Zip3[F231A] protein appears to be 2-3 fold more abundant than normal Zip3 protein in prophase arrested meiotic cells (Figures 7, 8, S5), one might suspect that unmodified Zip3 is more stable than modified Zip3 during meiotic prophase. Arguing against this possibility, however, is the fact that Zip3 is under-modified in *zmm* mutants yet does not appear dramatically more abundant (see standard deviations in Figure 8 legend).

We suggest a model (illustrated in Figure 8B) whereby unmodified Zip3 maintains a functional capacity that is diminished by phosphorylation, and this capacity is maintained (and perhaps further facilitated) by direct engagement with Zip1 at a nascent recombination ensemble. We propose that successful completion of intermediate steps in recombination (downstream of *ZMM* function) triggers a change in the interaction between Zip3 and Zip1, rendering Zip3 susceptible to PTM addition. If a component of the recombination pathway is missing, Zip3 and Zip1 fail to change configuration, and Zip3 remains under-modified. Under this model Zip1 serves a dual role when a component of the recombinosome is missing: Zip1 both protects Zip3 from degradation and maintains Zip3 in a functional state until the recombination ensemble becomes fully intact. When no recombination sites are present (i.e. a *spo11* mutant), Zip3 may exist in a mixture of modified and unmodified forms due to a relatively unstable Zip1-Zip3 interaction within pre-recombinosome ensembles off of chromosomes.

Our model is based on the observation that Zip3 is undermodified in *zmm* mutants, which fail to generate MutSγ crossovers. How can we fit into this picture the *zip3[F231A]* mutant, which does generate MutSγ crossovers? We propose that despite successful *ZMM* function and maturation of a recombination event, the Zip3[F231A] protein lacks the capacity to properly change its interaction with Zip1 in a timely fashion, which protects it from degradation but also from PTM addition (so long as Zip1 is present within the cell). Zip3[F231A]’s putative “stickiness” for Zip1 may stall crossovers at certain genomic positions, or hinder the mechanism that efficiently couples recombination to SC assembly, which could explain the mild synapsis deficiency observed in *zip3[F231A]* mutants (Figure S6). Finally, Zip3’s phenylalanine 231 appears to facilitate the degradation mechanism of Zip3 given the abundance of the Zip3[F231A] protein relative to wild type Zip3 protein in *zip1* null mutants.

### Mass spectrometry identifies shared proximity labeling targets of MutSγ pathway proteins

Thus far we have described the use of TurboID in yeast meiotic cells as a phenotypic tool to examine interactions that can be evaluated using a streptavidin blot. To explore whether *TurboID* fusions can be used as a discovery tool to potentially identify new factors involved in meiotic prophase functions associated with ZMM proteins, we conducted a pull down-mass spectrometry experiment. Protein was extracted from duplicate cultures of eleven strains, each homozygous for *ndt80* and a given *TurboID* fusion, or a “No TurboID” *ndt80* control strain. From each extract we purified biotinylated proteins using streptavidin-coated sepharose beads (see Methods). A trypsin digestion of proteins bound to the streptavidin beads was analyzed using ultra-high performance liquid chromatography coupled to tandem mass spectrometry (UPLC-MS/MS). We filtered out proteins that were identified in at least one replicate of the “no TurboID” control, and defined proximity labeling targets as those proteins identified in both replicates of a given TurboID fusion, with some exceptions: For the two distinct Zip4 TurboID fusions, we counted proteins identified in two replicates of one (Zip4iTurboID, for example) and one replicate of the other (Zip4-TurboID) as targets of either Zip4 fusion protein. Similarly, because Msh4-Msh5 and Ecm11-Gmc2 each assemble a heterocomplex, a detected protein was considered a target of a component of the complex (Msh4-TurboID or Msh5-TurboID, for example) even if only present in one biological replicate, so long as it was also detected in both biological replicates of the other strain. Using these criteria, we observed 19 targets of Zip2-TurboID, 88 targets of Zip4-TurboID, 85 targets of Zip4iTurboID, 28 targets of Spo16-TurboID, 27 targets of Zip3iTurboID, 52 targets of Mer3-TurboID, 51 targets of Msh4-TurboID, 48 targets of Msh5-TurboID, 19 targets of Mlh3-TurboID, 44 targets of Ecm11-TurboID, and 20 targets of TurboID-Gmc2 (Figure 9, Table S3).

**Figure 9.**
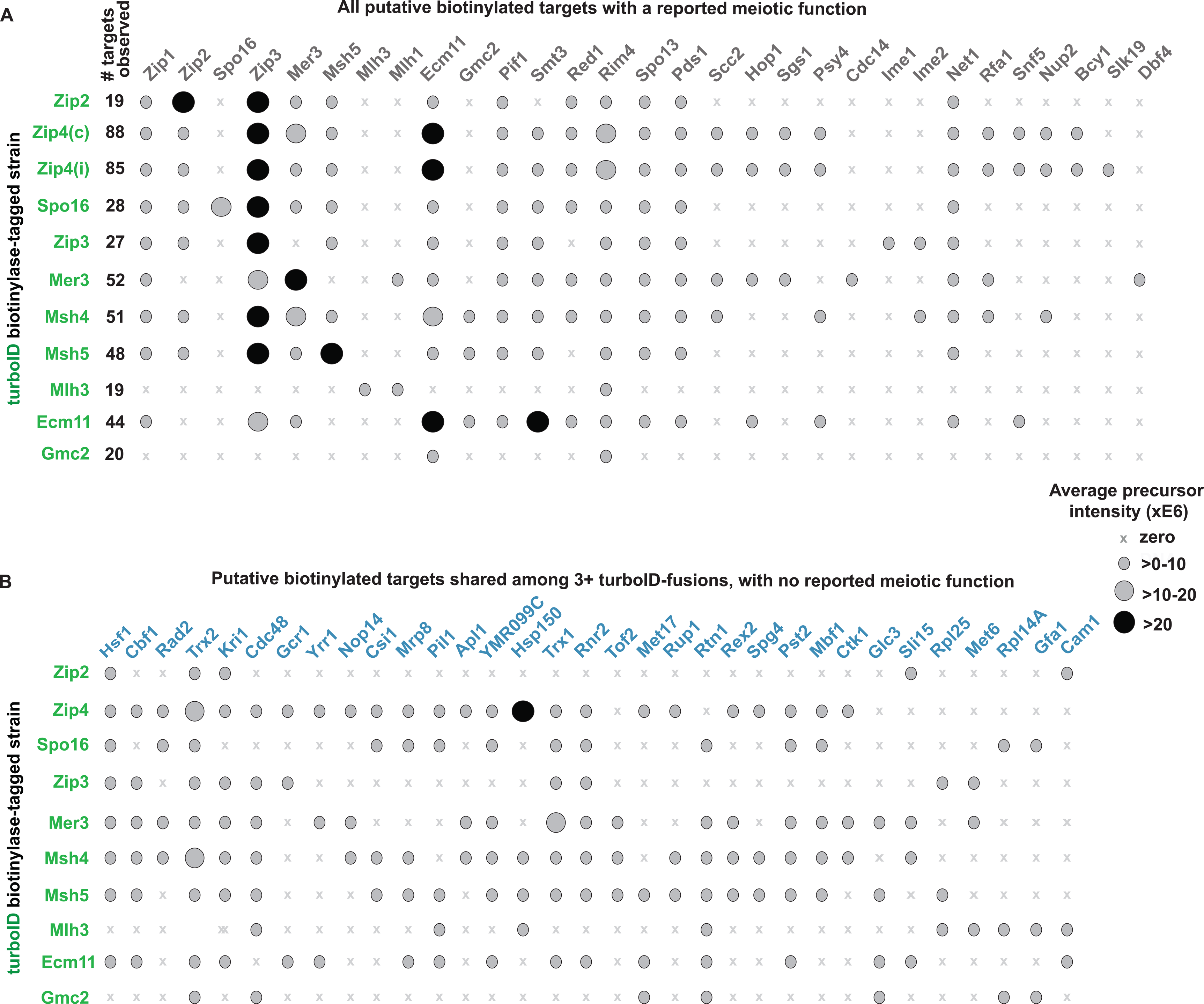
Mass spectrometry reveals additional shared proximity label targets of meiotic recombination associated proteins. Two biological replicates of twelve *ndt80* strains (eleven homozygous for a distinct *TurboID* gene fusion and one devoid of *TurboID*) were sporulated for ∼24 hours. Proteins extracted from each strain were incubated with streptavidin-coated beads, beads were washed and processed for ultra-high performance liquid chromatography coupled to tandem mass spectrometry (UPLC-MS/MS) analysis on an Orbitrap Eclipse Tribrid mass spectrometer (Thermo Scientific). Results were filtered to a 1% false discovery rate at the peptide and protein level using a target-decoy search and a reversed version of the full yeast proteome database. (A, B) Proteins carrying the TurboID fusion are listed on left *y* axis in green, and streptavidin-purified interactors for each TurboID fusion are listed across the top. Note Zip4(c) corresponds to a strain where the TurboID biotinylase is fused to the C terminus of the Zip4 protein, whereas Zip4(i) corresponds to a strain carrying *ZIP4iTurboID*, which encodes a protein containing an internal TurboID. Ovals indicate detection of a particular protein as a streptavidin-purified interactor in a given *TurboID* strain, with the smallest ovals indicating an average precursor intensity of >0-10, the larger, lightly-shaded ovals indicating an average precursor intensity of between 10 and 20, and the darkly-shaded large ovals indicating an average precursor intensity of greater than 20; an “x” indicates that the protein was not detected. The total number of targets observed for a given *TurboID* strain is listed in the “# targets observed” column in (A). Note that for most strains, a target was identified in both biological replicates of the *TurboID* strain and not in either biological replicate of the control. For *Zip4iTurboID* or *Zip4-TurboID*, a detected protein was considered a target even if present in only one biological replicate so long as it was detected in both biological replicates of the other strain (and not in either replicate of the control). Similarly, because Msh4-Msh5 and Ecm11-Gmc2 assemble heterocomplexes, a detected protein was considered a target of one component in the heterocomplex even if only present in a single biological replicate so long as it was also detected in both biological replicates of the strain carrying the TurboID fusion of the other component, (and in neither replicate of the control). (A) lists all protein targets identified with a previously reported meiotic function. (B) lists the shared targets of at least three TurboID fusion proteins that have no previously reported meiotic function. In (B), protein targets listed for Zip4 TurboID fusions were identified in both *ZIP4c-TurboID* biological replicates and at least one replicate of *ZIP4iTurboID*, or vice-versa. See Table S3 for full list of proteins identified by mass spectrometry.

As expected, proximity labeling targets of Zip4iTurboID show a large degree of overlap with targets of Zip4-TurboID (81 shared targets, 7 unique to Zip4iTurboID and 4 unique to Zip4-TurboID; Figure S7). Furthermore, most proximity labeling targets of Msh4 (43/51) were identified among the 48 targets of Msh4’s heterodimeric partner Msh5 (Figure S7). These results boost confidence in the success of our pull-down mass spectrometry approach and give a measure of validation to targets defined by a low number of biological replicates (two for each strain). Ecm11 and Gmc2 also form a heterocomplex in the meiotic cell (Humphryes *et al*. 2013); while neither the Ecm11-TurboID nor TurboID-Gmc2 fusion protein is functional in terms of SC assembly (Figure S1), nevertheless 19 of 20 proximity labeling targets identified for TurboID-Gmc2 were also identified targets of Ecm11-TurboID (Figure S7).

Zip2, Zip4, and Spo16 proteins form a stable subcomplex (ZZS) (De Muyt *et al*. 2018). Consistent with this, 14 targets are shared between Spo16-TurboID and Zip2-TurboID, and 16 of19 Zip2-TurboID targets as well as 25 of 28 Spo16-TurboID targets are also targets of a Zip4 TurboID fusion protein.

Furthermore, Zip2 is detected as a target of Zip2-TurboID, Zip4-TurboID, Zip4iTurboID, and Spo16- TurboID. Spo16 was detected as a target of Spo16-TurboID, but not of Zip2-TurboID nor either of the Zip4-TurboID fusions, however its small size makes it less likely to be identified in a mass spectrometry experiment. Curiously, Zip4 (a relatively large protein) was not detected as a proximity labeling target of any of our meiotic TurboID fusions, even though Zip4-TurboID and/or Zip4iTurboID proximity label several different meiotic factors including Zip1, Zip2, Zip3, Mer3, Msh5, and Ecm11. Lysine residues, which serve as a substrate for the biotinylation reaction, occur at a similar frequency in Zip4 relative to Zip3 (∼8% for both proteins). Perhaps Zip4 is configured with other ZMM proteins such that the solvent-accessible TurboID region of each ZMM bait is outward facing, away from a buried Zip4 protein.

Consistent with the idea that meiotic recombination and SC assembly factors form ensembles at recombination/synapsis sites, many of the same targets were identified among *TurboID* fusion strains that correspond to components not yet known to be part of stable subcomplexes. Figure 9A illustrates all targets identified with a known meiotic function, revealing many meiotic targets shared by more than one *TurboID* strain. For example, Zip3 and Zip1 are targets of every TurboID fusion except for Mlh3- TurboID and TurboID-Gmc2, and Ecm11 was identified as a target of all TurboID fusions except for Mlh3-TurboID (Figure 9A). The SUMO protein (encoded by the *SMT3* gene) was found to be a target of most TurboID fusion proteins, and peptides identified by mass spectrometry for SUMO were most abundant in the *ECM11-TurboID* strain, consistent with Ecm11 being one of the most abundant SUMOylated proteins in yeast meiotic prophase cells (Bhagwat *et al*. 2021). These data also indicate that Msh4-TurboID and Msh5-TurboID (components of MutSγ) proximity label Zip2 as well as the Mer3 helicase, possibly reflecting the engagement of MutSγ with a recombination intermediate.

Another identified target in all *TurboID* fusion strains except *MLH3-TurboID* and *TurboID-GMC2* is the Pif1 helicase. Pif1 has been found to localize to meiotic recombination sites and to be engaged with recombination intermediates in a manner is that is restrained by Mer3 (Vernekar *et al*. 2021); these proximity labeling data raise the possibility that Mer3 acts to regulate Pif1 in the physical context of the MutSγ pathway proteins Zip2, Zip4, Spo16, Zip3 and the MutSγ heterodimer (Msh4-Msh5).

Somewhat unexpectedly, Spo13 and Pds1 were identified as targets of nearly all TurboID fusions. A dramatic meiotic consequence of Pds1 loss (which binds securin and thereby protects cohesion from destruction) is premature sister chromatid separation during anaphase I, however evidence has been reported for Pds1 having a distinct role in meiotic recombination and synaptonemal complex assembly (Cooper *et al*. 2009). Spo13 binds the Cdc5 kinase and regulates signaling pathways that govern exit from meiosis after prophase, localizes to centromeres, and regulates kinetochore mono-orientation during meiosis I (Katis *et al*. 2004; Lee *et al*. 2004; Matos *et al*. 2008). The fact that Spo13 is a proximity labeling target of several MutSγ pathway pro-crossover proteins in *ndt80* cells suggests that it plays a role at MutSγ crossover sites during meiotic prophase, or perhaps that MutSγ pathway pro-crossover proteins engage with Spo13 at centromeres.

A target of every TurboID fusion is the meiosis-specific mRNA-binding protein Rim4. Rim4 forms amyloid-like aggregates in the cytoplasm of meiotic prophase cells and represses the translation of at least a subset of developmentally regulated mRNAs during meiotic prophase, but Rim4 may also bind mRNAs that are not translationally repressed (Berchowitz *et al*. 2013; Berchowitz *et al*. 2015). An economical explanation for Rim4 being a target of all TurboID fusions examined is that Rim4 associates with the mRNA encoding each of these fusion proteins, positioning it in the immediate periphery of the TurboID polypeptides as they are undergoing translation. Another possibility is that Rim4 has a yet undescribed function in ZMM- or SC-associated pathways.

Mlh1, Mlh3, and Rim4 were the only identified meiotic protein targets of Mlh3-TurboID in these experiments, which utilized *ndt80* meiotic cells. The paucity of Mlh3 targets may be explained by the fact that the Mlh1-Mlh3 heterodimer (MutLγ) – mediated resolution of crossover recombination intermediates occurs downstream of Ndt80 activation (Allers and Lichten 2001; Voelkel-meiman *et al*. 2015).

Importantly, several proteins identified as targets of more than one TurboID fusion protein during meiotic prophase do not have a reported role in meiosis (Figure 9B lists such protein targets shared by at least three different *TurboID* strains). These targets are of particular interest for future study, as at least some of them likely represent factors with meiotic functions that have not yet been investigated.

## Discussion

### Proximity labeling reinforces and refines a picture of ZMM proteins on and off the meiotic recombination intermediate

Zip2, Zip3, Zip4, Spo16, Mer3, Msh4 and Msh5 (collectively known as ZMM proteins) have long been known to function in the same meiotic crossover pathway, along with the pro-crossover form of Zip1 (Hunter 2015). As such, several of these proteins have been found to co-localize with one another at recombination sites on mid-meiotic prophase chromosomes. Recombination intermediates are also a major assembly site for the elaborate synaptonemal complex (SC) structure, thus the Ecm11-Gmc2 heterocomplex, a structural building block of SC together with the Zip1 protein, is also expected to localize to these ZMM-associated chromosomal sites. However, how all these proteins physically and functionally interact with one another prior to and during recombination has thus far only been investigated with a limited number of cytological labeling experiments. Here, our proximity labeling for several ZMM “bait” proteins i) reinforces the idea that ZMM proteins and SC proteins co-exist in ensembles within the yeast meiotic prophase nucleus and ii) refines our picture of the relationships between Zip3 and Ecm11 and the other ZMM proteins.

Figure 10 presents an illustration of the proximity labeling interactions found between ZMM proteins, Zip1 and Ecm11-Gmc2 primarily guided by our streptavidin blotting data (i.e. interactions that target Zip3 or Ecm11 protein). One take home from our experiments is that ZZS, Zip1, and Zip3 form ensembles within the meiotic nucleus independent of SC or polycomplex structures. Zip2-TurboID and Spo16-TurboID proximity label the Zip3 protein in a manner that is dependent on each other and Zip4, consistent with the fact that Zip2, Zip4, and Spo16 purify as a stable subcomplex (“ZZS”) and rely on one another for their localization to meiotic chromosomes (Tsubouchi *et al*. 2006; De muyt *et al*. 2018).

**Figure 10.**
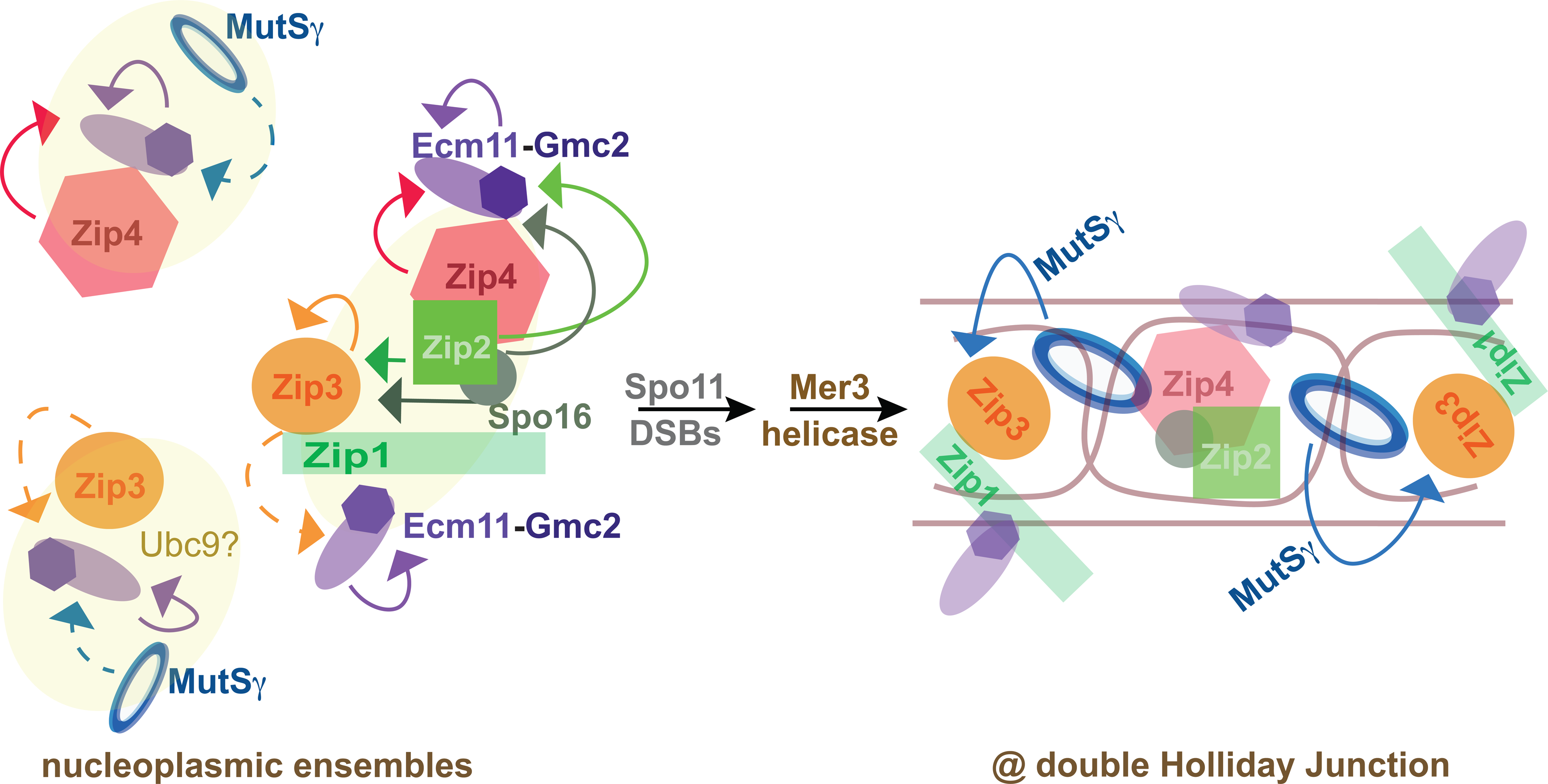
Zip3 and Ecm11 are proximity labeled in multiple ensembles on and off recombination intermediates. Cartoon ensembles in which Zip3 or Ecm11 proximity labeling occurs, inferred from streptavidin blotting experiments presented in Figures 1, 2, 3, and S2. At left is depicted three ensembles that support proximity labeling of Zip3 (orange circle) or the Ecm11-Gmc2 heterocomplex (purple) independent of recombination (Spo11 activity). Yellow ovals are meant to indicate the possibility that other (unknown) proteins might bridge the featured proteins within the ensemble. Arrows indicate *trans* biotinylation of either Ecm11 or Zip3, the color of the arrow corresponds to the bait protein doing the proximity labeling. A complex containing Zip4 and Ecm11-Gmc2 is shown, based on the direct two-hybrid interaction known between Ecm11 and Zip4 and on the fact that Ecm11 proximity labeling by Zip4iTurboID occurs independently of other meiotic proteins, including Zip2 and Spo16; The Msh4- Msh5 heterodimer (MutSγ) is depicted in blue close to Zip4-Ecm11, to account for the capacity of Msh4- TurboID to proximity label Ecm11 in a manner that is partially dependent on Zip4 (Figure 3B). Another ensemble carries Zip3 and Ecm11-Gmc2 and is meant to reflect the Zip4- and Zip1-independent proximity labeling of Ecm11 by Zip3. MutSγ is close to the Zip3-Ecm11 complex, to account for the capacity of Msh4-TurboID to proximity label Ecm11 in a manner that is partially dependent on Zip3; since Zip3 is a SUMO E3 conjugase that attenuates Ecm11 SUMOylation (Humphryes *et al*. 2013) and MutSγ is directly SUMOylated by the Ubc9 E2 (He *et al*. 2021) we speculate that Ubc9 might be a component of this complex (Figure 3B). The third ensemble carries ZZS (red Zip4, army green Spo16, and bright green Zip2) as well as Zip1-Zip3; a yellow oval surrounds these factors because the links that connect ZZS to Zip1-Zip3 are currently unknown. In this ensemble, ZZS proteins Zip2 and Spo16 can proximity label Zip3 independent of recombination initiation and SC/polycomplex assembly. Cartoon at right depicts a mature DNA joint molecule (double Holliday junction) formed downstream of Spo11- mediated DNA double strand breaks, strand invasion, and ZMM function including Mer3 helicase activity. In the context of the mature joint molecule, MutSγ proximity labels the Zip3 protein.

We furthermore found that Zip2-TurboID (ZZS) proximity labels Zip3 even in the combined absence of recombination initiation and polycomplex structure (in *spo11 ecm11* double mutants). Prior cytological data indicate that ZZS proteins localize along with Zip3 and MutSγ at polycomplex structures in *spo11* mutants (Tsubouchi *et al*. 2006; Voelkel-meiman *et al*. 2019), but our data furthermore indicate that ZZS proteins remain in proximity to Zip3 even in the absence of SC or polycomplex structure, and this interaction is dependent on the pro-crossover activity of Zip1.

Our experiments also revealed a proximity labeling interaction between Zip4 and Ecm11 that occurs independent of recombination and ZZS proteins Zip2 or Spo16, consistent with the direct interaction previously identified between Zip4 and Ecm11 (Pyatnitskaya *et al*. 2022). (By contrast, Zip2 and Spo16 rely on one another as well as Zip4 to proximity label Ecm11.) Moreover, our data indicate that Zip3 proximity labels Ecm11 in a manner that is independent of recombination and Zip4, and only partially dependent on Zip1 (Figure 10). This is interesting in light of Zip3’s role in attenuating Ecm11 SUMOylation (Humphryes *et al*. 2013); perhaps Zip1-independent proximity between Zip3 and Ecm11 occurs within a SUMOylation complex containing the Ubc9 E2 SUMO conjugase protein and the Zip3 E3 ligase. We also find that Msh4 (MutSγ) proximity labels Ecm11 in a manner that partially depends on Zip4, Zip1, and Zip3. This result is intriguing because it suggests that MutSγ independently engages with both Zip4- and Zip3-containing ensembles that carry Ecm11-Gmc2. MutSγ has recently been found to be directly targeted by the E2 SUMO conjugase Ubc9 (He *et al*. 2021), suggesting MutSγ might also be present at the hypothesized E1-E2-Zip3 ensemble (Figure 10).

Finally, streptavidin blotting revealed interesting information about the Mer3 helicase, whose function within the ZMM pathway is not fully understood (Duroc *et al*. 2017; Altmannova *et al*. 2023). We find that, unlike ZZS proteins Zip2-TurboID or Spo16-TurboID, Msh4-TurboID proximity labels Zip3 only if recombination initiation, Rad51-Dmc1-mediated strand invasion, and the Mer3 helicase are intact. This result strongly suggests that the Mer3 helicase promotes a step in the maturation of the recombination intermediate that is required for MutSγ to join the recombinosome ensemble in a manner that allows proximity labeling of Zip3 by Msh4-TurboID. One possibility is that Mer3’s helicase function influences the three-dimensional structure of the DNA joint molecule-protein intermediate in a manner that is conducive to binding by Zip1-Zip3 ensembles and/or MutSγ (Figure 10). We note that Mer3 may promote the maturation of a recombination intermediate through a non-catalytic activity, such as an interaction with a partner protein or and/or a specific DNA structure, as is suggested by the mild meiotic phenotypes of the mer3[K167A] helicase-dead mutant (Duroc *et al*. 2017). The later arrival of MutSγ at a nascent recombination intermediate is consistent with the fact that some *zip3* mutant alleles display proximity labeling of Zip3 by Zip2-TurboID but not Msh4-TurboID, but the reverse outcome (proximity labeling of Zip3 by Msh4-TurboID but not Zip2-TurboID) was not observed in any *zip1* or *zip3* mutant.

### A unique functional relationship between Zip3 and Zip1

We show that, among ZMM-associated proteins, Zip1 protein is uniquely required for maintaining abundant Zip3 within the meiotic prophase cell. The fact that Zip1 protects Zip3 from degradation might explain the Zip1-dependency of Zip3 proximity labeling by several ZMM proteins. However, some *zip1* mutants, such as the SC-deficient but crossover-proficient *zip1[Δ 21-163]* strain, exhibit abnormally low levels of Zip3 within the cell yet normal proximity labeling of Zip3 by both Zip2-TurboID and Msh4- TurboID, consistent with the MutSγ crossover proficiency of this strain (Voelkel-meiman *et al*. 2016). Thus, we suggest that Zip1 both protects Zip3 from degradation *and* promotes its proper positioning within the recombination ensemble; one possibility is that the entirety of Zip1’s pro-crossover role is to position Zip3 properly within the recombinosome. The low level of Zip3 in *zip1[Δ 21-163]* cells also raises the possibility that Zip1 may be maximally capable of protecting Zip3 from degradation when Zip1 itself is capable of SC assembly.

We furthermore discovered that Zip1 counters the post-translational modification (PTM) of Zip3 when a ZMM component is defective. This activity of Zip1 in countering Zip3 PTMs is most clear in the *zip3[F231A]* mutant, which is both less susceptible to degradation when Zip1 is missing, and defective in accumulating PTMs when Zip1 is present. We suggest that ZMM activity promotes a shift in an early stage recombinosome ensemble, reflecting a maturation step in the DNA joint molecule recombination intermediate, that triggers a change in the ZMM protein ensemble configuration such that Zip1 can no longer block Zip3 from acquiring PTMs. When a ZMM protein is missing and this maturation step fails, Zip1 prevents Zip3 from acquiring PTMs. Under our model, recombination intermediates undergo proper maturation in the *zip3[F231A]* mutant, but the Zip3[F231A] protein is less efficient at changing its configuration vis-à-vis Zip1. PTM acquisition by Zip3 may predispose the protein to degradation either immediately or after the *ndt80* arrest point at mid-late meiotic prophase, and/or prevent it from carrying out off-target effects on ongoing parallel pathways during meiotic progression.

### A promising phenotypic discovery tool for yeast meiosis, with limitations

This study uses proximity labeling in two ways: i) as a phenotypic tool to better understand the physical and functional relationships between proteins that appear to be components of the same or related pathways (i.e. MutSγ recombination and SC assembly), and ii) as a discovery tool to potentially identify new factors that function with known meiotic proteins in budding yeast. We show that many TurboID fusions with yeast meiotic recombination proteins are at least partially functional, and that the TurboID biotinylase can function not only at the N or C terminus but also when positioned internal to a protein. We also demonstrate that a relatively straightforward streptavidin blotting approach can be used to test, in parallel, genetic dependencies for specific proximity labeling events. Streptavidin pull-down followed by mass spectrometry reveals an even broader spectrum of potential interactors.

A clear limitation of proximity labeling in yeast meiosis (as we have currently performed it) is the low abundance of potential protein targets. Streptavidin blots could be a powerful phenotyping tool, as they are typically more cost-effective than pull downs followed by mass spectrometry and should be repeatable in a way that mass spectrometry analysis of very low abundance peptides may not be due to the inherent limit of detection associated with UPLC-MS/MS-based methods. However, streptavidin blots revealed only a few proximity labeling targets of the TurboID fusion proteins we evaluated. We note that in our examination of six timepoints across meiotic prophase in *ndt80* cells, Zip3iMYC and Zip1 proteins are not detectable on the blot until the third timepoint (15 hr; Figure S4). Thus, an abundance of Zip3 sufficient to detect proximity labeling within meiotic cell extracts may only occur at the late (24 hr) timepoint in *ndt80* strains when nearly all cells have reached the pachytene stage, and the proximity labeling of any lower abundance proteins is likely undetectable on a streptavidin blot. Approaches to enrich for a candidate target protein population (by fractionation or immunoprecipitation) may be required for the detection of proximity labeling events on low abundance proteins.

A second limitation of the streptavidin blotting approach for detecting proximity labeling targets is the presence of a few highly abundant, naturally-biotinylated proteins, which could obscure the signal of a *bona fide* target. We attempted to use an *arc1* knockout strain for our proximity labeling analyses (*ARC1* encodes the naturally biotinylated proteins migrating near 42 kDa (Kim *et al*. 2004)), but found that meiosis is defective in this strain. One way to circumvent the issue of naturally biotinylated species obscuring a target signal is to pre-incubate protein samples with streptavidin and then run a traditional western blot, as described in (Xiang and Koshland 2021). Using this approach with *Zip2-TurboID ZIP3iMYC*, *MSH4-TurboID ZIP3iMYC*, and *ZIP4iTurboID ZIP3iMYC* strains, we found that an anti-MYC western blot detected two forms of the Zip3iMYC protein: the unbiotinylated form and a shifted Zip3iMYC species that corresponds to biotinylated Zip3iMYC bound to streptavidin (Figure S8).

The successful covalent modification of a lysine in the target protein by the TurboID biotinylase depends on the bait-target protein conformation as well as the primary amino acid sequence of the target protein itself. Thus, like many tools for reporting a physical relationship between proteins, many “proximities” will fail to be revealed using this method. However, the determination of just two proximity interaction targets, Zip3 and Ecm11, by streptavidin blotting has led to new insights into the relationships between components of the MutSγ pathway. Furthermore, the unexpected proximity labeling targets identified by mass spectrometry supply potentially new functional components of meiotic recombination and synapsis pathways to explore.

## Methods

### Strains and crossover data

Strains created for this study are isogenic with the BR1919-8B background (Rockmill and Roeder 1998), and are listed in Table S1. Knockout alleles and C-terminal *TurboID* fusions were created by standard recombination-based gene targeting procedures. Plasmid pFB1420 (pFA6a-*TurboID-3xMYC- kanMX6*; Addgene) was used to amplify DNA for creating in-frame *TurboID* fusion alleles; note that *3xMYC* follows TurboID in C-terminal fusion alleles, while alleles encoding internal TurboID fusions carry only the four first residues of the 3xMYC tag; the *TurboID* internal fusion alleles and *zip1* and *zip3* non-null alleles were created by *CRISPR*-Cas9-mediated allele replacement as in (Voelkel-meiman *et al*. 2019). Zip3 and Zip4 epitope tags are positioned internal to the gene ORFs (after residue 91 in Zip4 and after residue 245 in Zip3), as described in (Tsubouchi *et al*. 2006) and notated as *i3xMYC, i3xHA* or *iTurboID*. *MSH4-13xMYC* and *MSH4-3xHA* were created using plasmids *pFA6a-13xMYC-kanMX6* and *pFA6a-3HA-kanMX6* respectively (Longtine *et al*. 1998).

Genetic crossover data was compiled and processed as described in (Voelkel-meiman *et al*. 2019).

### Cytological analysis and imaging

Meiotic nuclei from various *ndt80* homozygous strains were surface-spread on glass slides and imaged as described in (Voelkel-meiman *et al*. 2016). The following primary antibodies were used: affinity purified rabbit anti-Zip1 (YenZym Antibodies, LLC, as in (Sym *et al*. 1993); 1:100), mouse anti-cMYC (clone 9E10 Abcam; 1:200), mouse anti-Gmc2 (raised against purified Gmc2, ProSci Inc., 1:800), guinea pig anti-Gmc2_Ecm11 (raised against a co-purified protein complex; ProSci Inc., 1:800), Rabbit anti-HA (Abcam; 1:100), and rabbit anti-Red1 (gift from G.S. Roeder, (Smith and Roeder 1997); 1:200).

Secondary antibodies conjugated with Alexa Fluor dyes (Jackson ImmunoResearch) were used at 1:200 dilution. Microscopy and image processing were performed using a Deltavision RT imaging system (General Electric) adapted to an Olympus (IX71) microscope.

### Western and streptavidin blotting

Protein was extracted from from 5 mL of sporulating cell culture by TCA precipitation as in (Hooker and Roeder 2006); cells were vortexed with glass beads for 10 minutes at 4°C. The final protein pellet was resuspended in 2× Laemmli sample buffer supplemented with 30 mM DTT, at a concentration of ∼20 μg/μl. Protein samples were heated for 10 minutes at 65°, centrifuged at top speed and ∼100 μg was loaded onto either an 8% or a 12% polyacrylamide/SDS gel. Gels were run either at 80V (for Zip3-iMYC studies) or 100V (for TurboID, Msh4-MYC and Zip4-iHA studies). Protran 0.2μm nitrocellulose (Amersham) was used as the transfer membrane following the manufacturer’s recommendation. Transfer of proteins to nitrocellulose was performed in CAPS pH 11-10% ethanol buffer for TurboID and Zip3-iMYC studies, and Towbin Buffer-10% methanol for MSH4-MYC and Zip4-iHA studies; stir bar and ice pack were used at 100V for transfer. 12% PAGE blots were transferred for 45 minutes whereas 8% PAGE blots were transferred for 1 hour. Membranes were allowed to dry (>30 minutes) after transfer, then washed in 1x PBST buffer. Ponceau S was used to detect total protein and quality of transfer to the membrane, then the membranes were washed twice more with 1x PBST. Membranes for TurboID-biotin studies were blocked using 3% BSA in 1x PBST for 30 minutes and washed once in 1x PBST. Membranes were incubated overnight at 4°C with STAR5B Streptavidin:HRP (BioRad) at a 1:15,000 dilution in 1x PBST. Membranes were then washed three times in 1x PBST and imaged as described below. Membranes for antibody analysis were blocked in 5% non-fat dry milk powder/ 2% BSA in PBST for 30 minutes and washed once in 1x PBST. Membranes were incubated overnight at 4°C with primary antibodies in 1x PBST: Mouse anti-MYC (9E10.3, Abcam) at 1:2000 for Zip3-iMYC, 1:5000 for Msh4- MYC blots; rabbit anti-Zip1 and rat anti-tubulin YOL1/34 (Abcam) at 1:10,000, rabbit anti-Fpr3-C (gift of Dr. Jeremy Thornton) at 1:50,000, mouse anti-HA.11 (Abcam) at 1:1000. Secondary antibodies (HRP-conjugated AffiniPure Donkey anti-rabbit, goat anti-mouse (Jackson ImmunoResearch), and goat anti-rat (Santa Cruz)) were used at 1∶10,000 in 1x PBST for 1 hour at room temperature. ECL Prime Western Detection Reagent (Amersham) was used to visualize probes on the membranes; a G:Box mini (Syngene) was used to detect chemiluminescence and ImageJ (https://imagej.nih.gov/ij/index.html) was used to analyze the data.

For streptavidin shift experiments, 100 μg of protein gel sample was incubated for ten minutes with streptavidin (Invitrogen #434302), at a final concentration of 1 μg/μL in a 10 μL final volume, before loading on to the polyacrylamide gel.

### Streptavidin pull-down followed by analysis using ultra-high performance liquid chromatography coupled to tandem mass spectrometry (UPLC-MS/MS)

Protein was extracted from ∼40 mL of sporulating cell culture at the 24 hr timepoint, using TCA precipitation as described above. TCA preparations are done with 5 mL culture volumes, approximately eight TCA preparations were performed for each strain/replicate (12 strains total with two biological replicates), the excess from an originally 45 mL culture was processed for chromosome spreads to ensure strains entered meiosis successfully. The eight TCA pellets from each strain/replicate were consolidated into one tube using ∼975 μL of 2% SDS/Bead Binding Buffer (50 mM Tris-Cl pH 7.5, 150 mM NaCl, 1.5 mM MgCl2, 1 mM EGTA, 2% SDS, 1% NP-40, 2 μg/mL sodium deoxycholate, 0.5 mM DTT); protein pellets were heated for two minutes at 65 degrees, disrupted using a P1000 pipette tip, and allowed to rock at room temperature ∼30 minutes. Protein solutions were then heated at 65°C for ten minutes, microfuged for 30 seconds at top speed, and the soluble fraction (∼950 μL) added to a protein lo-bind microfuge tube (Eppendorf) carrying equilibrated streptavidin-sepharose beads (General Electic # 17-5113-01). A 30 μl volume of streptavidin beads was equilibrated for each sample, via ten washes in 1 mL of RIPA wash buffer (50 mM Tris-Cl pH 7.5, 150 mM NaCl, 1.5 mM MgCl2, 1 mM EGTA, 0.1% SDS, 1% NP-40, 2 μg/mL sodium deoxycholate, 0.5 mM DTT). Bead-protein solution was incubated for one hour at room temperature and then overnight at 4°C. Beads were washed three times in 2% SDS wash buffer (50 mM Tris-Cl pH7.5, 2% SDS), then three times on ice in cold RIPA wash buffer, and five times in 1 mL of 20 mM ammonium bicarbonate. Supernatant was removed after the final wash and beads were stored at -80°C until their transport to University of Connecticut’s Proteomics and Metabolomics Facility.

Beads were prepared for UPLC-MS/MS analysis using three washes in 0.1M ammonium bicarbonate, after which cysteine reduction and alkylation was performed using 5 mM dithiothreitol in 0.1M ammonium bicarbonate for 1.5 hour and 10mM iodoacetamide in 0.1M ammonium bicarbonate for 45 minutes in the dark, respectively. Directly afterward, sequencing-grade trypsin (Promega) was added at a 1:20(w/w) enzyme:protein ratio for 16 hour digestion at 37°C. Tryptic peptides were removed with the supernatant, quenched to a final pH of 2.5 using formic acid, then fully desalted using Pierce peptide desalting spin columns (Thermo Scientific) per manufacturers’ instructions. Desalted and dried peptides were resuspended in 0.1% formic acid, quantified, and diluted to 0.3mg/mL. A 1μL aliquot containing 300ng of peptides was loaded onto a Waters nanoEase m/z Peptide BEH C18 analytical column, separated using a 1-hour UPLC reversed-phase gradient (Solvent A: 0.1% formic acid in water, Solvent B: 0.1% formic acid in acetonitrile), and eluted directly into the Orbitrap Eclipse Tribrid mass spectrometer (Thermo Scientific) using positive mode electrospray ionization. The acquisition method incorporated a TopN data-dependent acquisition mode with a maximum cycle time of 3 seconds. Both MS and MS/MS scans were acquired at high resolution in the Orbitrap mass analyzer. Peptide and protein identifications were achieved by searching the raw data against the full Uniprot reference proteome for *Saccharomyces cerevisiae* (identifier UP000002311, accessed 07/05/2022) plus a custom FASTA database containing the turboID-tagged protein construct sequences using MaxQuant v1.6.10.43 (Cox and Mann 2008). Variable modifications included methionine oxidation, acetylation of the protein N-terminus, asparagine or glutamine deamidation, lysine biotinylation, and fixed carbamidomethyl on cysteine residues. All search results were filtered to a 1% false discovery rate at the peptide and protein levels using a target-decoy search. Scaffold 5 (www.proteomesoftware.com) was used to visualize the resulting data, and a protein identification threshold of at least two peptides per protein was used. Protein level quantitation values were calculated as “average precursor intensities”. Listed in Table S3 are the average precursor intensity values for all proteins identified in any of the replicates from a *TurboID* fusion that are not found in either replicate of the *no TurboID* control strain.

## Data Availability

Strains, plasmids and detailed protocols are freely available upon request.

## Acknowledgements

We thank Jenna Sneifer, and all undergraduates in the 2021 and 2022 MB&B 394 Advanced Lab in Molecular Genetics for creating several of the *TurboID* fusion strains used as bait in this study. We thank Samantha Hughes for creating several novel *zip1* mutant alleles, and Dr. Lina Yisehak for measuring crossover frequency in Sam Hughes’ *zip1* strains. This work was supported by National Institutes of Health grant GM116109 to A.J.M. The authors would like to acknowledge the NIH S10 High-End Instrumentation Award 1S10-OD028445-01A1, which supported this work by providing funds to acquire the Orbitrap Eclipse Tribrid mass spectrometer housed in the University of Connecticut Proteomics & Metabolomics Facility.

