## Supplemental Figures and Tables for "Proximity labeling reveals new insights into the relationships between meiotic recombination proteins in *S. cerevisiae*"

Figure S1

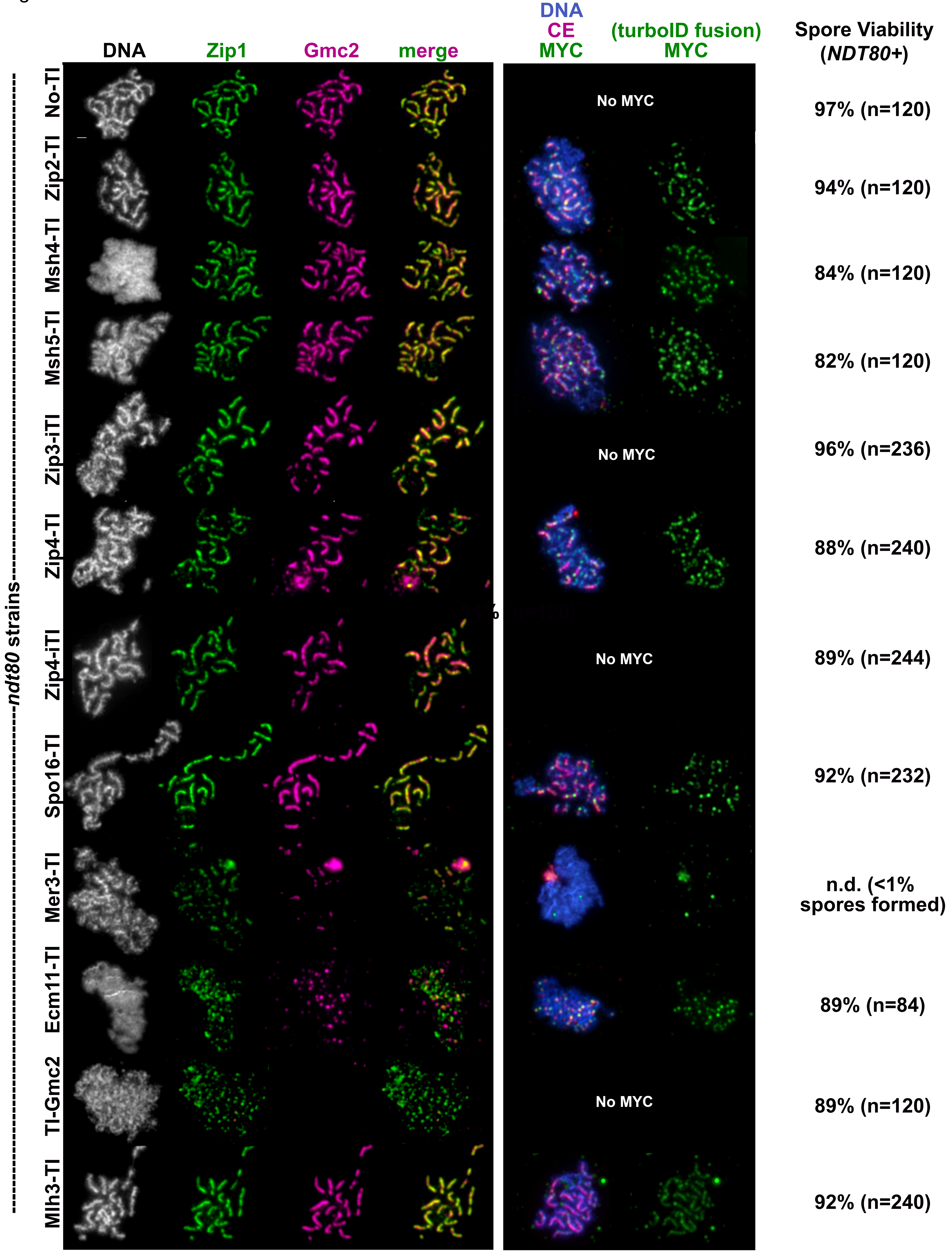
DNA
Zip1
Gmc2
merge
DNA
CE
MYC
(turboID fusion)
MYC
Spore Viability
(NDT80+)
ndt80 strains
No-TI
Zip2-TI
Msh4-TI
Msh5-TI
Zip3-iTI
Zip4-TI
Zip4-iTI
Spo16-TI
Mer3-TI
Ecm11-TI
TI-Gmc2
Mlh3-TI
No MYC
No MYC
No MYC
No MYC
97% (n=120)
94% (n=120)
84% (n=120)
82% (n=120)
96% (n=236)
88% (n=240)
89% (n=244)
92% (n=232)
n.d. (<1%
spores formed)
89% (n=84)
89% (n=120)
92% (n=240)

**Figure S1. Most TurboID fusion proteins support SC assembly, apart from Mer3-TurboID, Ecm11-TurboID, and TurboID-Gmc2.** Images show immunofluorescence on surface-spread meiotic chromosomes from *TurboID* strains homozygous for *ndt80*. Each row in both blocks of images corresponds to a different strain, with the top row carrying no *TurboID* and the lower rows homozygous for the *TurboID* fusion indicated at left. The first block of images (four columns) displays surface-spread meiotic nuclei that are each labeled with DAPI (white, first column), anti-Zip1 (green, second column), or anti-Gmc2 (magenta, third column). The fourth column shows the merge of anti-Zip1 and anti-Gmc2. C-terminal TurboID fusion proteins have a 3xMYC epitope, allowing visualization of a particular fusion protein on mid-meiotic prophase chromosomes: Rows in the second block of images display nuclei from the same strain labeled with DAPI, anti-Ecm11-Gmc2 (CE), and anti-MYC (blue, magenta, green, respectively, in the first column), and anti-MYC alone (green) in the second column. Bar, 1 micron. Spore viability for *NDT80+* versions of each *TurboID* fusion strain is listed at right (*n*=number of dissected spores evaluated).

Figure S2

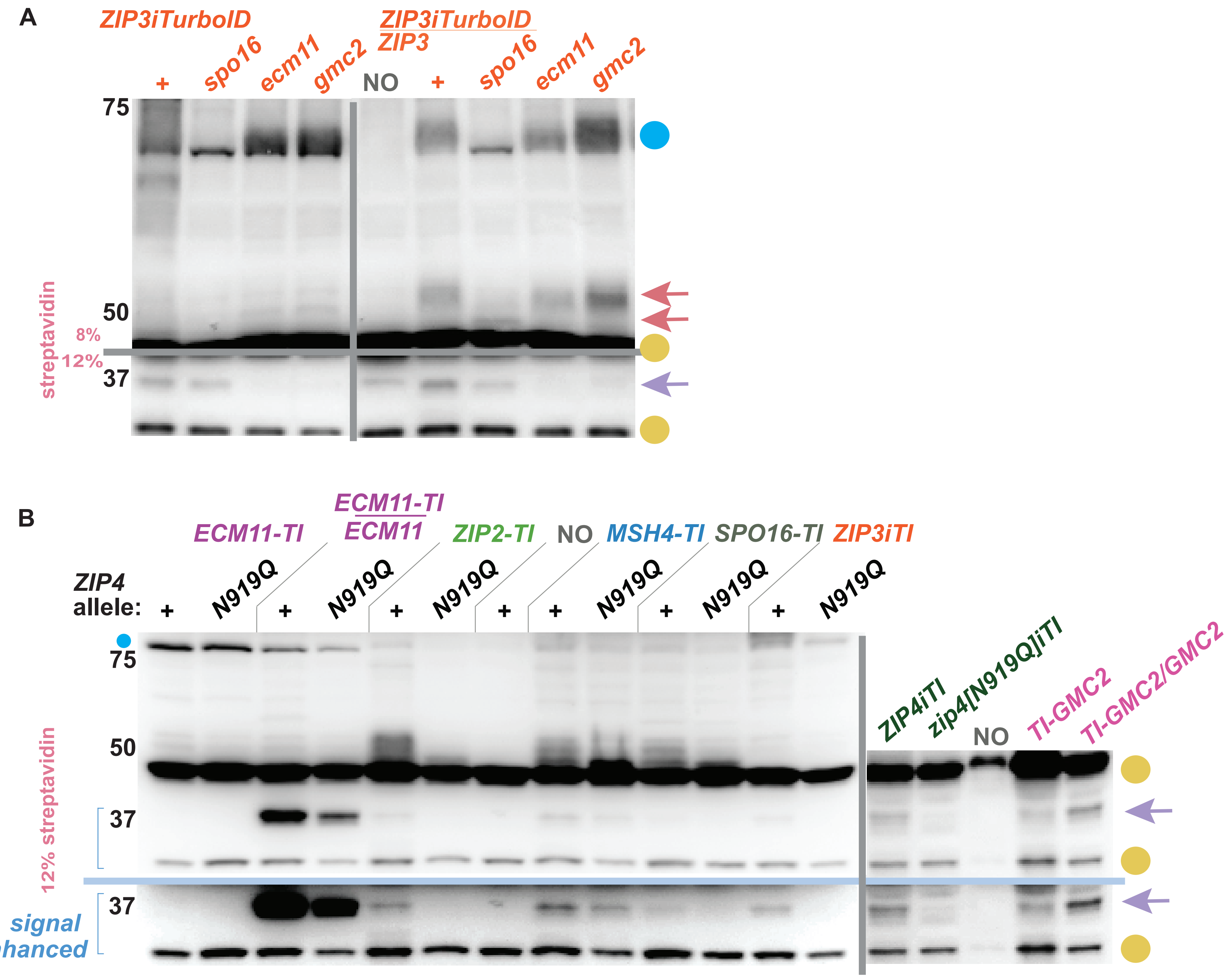
A
ZIP3iTurboID
+
spo16
ecm11
gmc2
ZIP3iTurboID
ZIP3
NO
+
spo16
ecm11
gmc2
75
50
37
8%
12%
streptavidin
B
ECM11-TI
ECM11-TI
ECM11
ZIP2-TI
NO
MSH4-TI
SPO16-TI
ZIP3iTI
ZIP4
allele:
+
N919Q
+
N919Q
+
N919Q
+
+
N919Q
+
N919Q
+
N919Q
ZIP4iTI
zip4[N919Q]iTI
NO
TI-GMC2
TI-GMC2/GMC2
75
50
37
37
12% streptavidin
signal
enhanced

**Figure S2. The 37 kDa protein proximity labeled by Zip2-TurboID and Ecm11-TurboID is likely Ecm11 itself.** Blots display proteins extracted from *ndt80* cells arrested at mid-meiotic prophase, separated on an 8% or 12% polyacrylamide gel (indicated in pink at left of blot), and probed with streptavidin to detect biotinylated proteins. Yellow circles indicate naturally biotinylated proteins, while purple arrows indicate the 37 kDa biotinylated species. Blots in (A) show biotinylated proteins from strains either homozygous or heterozygous for *ZIP3iTurboID*, and additionally homozygous for meiotic mutant alleles indicated at the top of the blot. Grey lines indicate independent membranes. The blue circle at the right of the blot indicates proteins migrating at positions consistent with that of Zip3iTurboID, while pink arrows correspond to untagged biotinylated Zip3 protein. Blot in (B) displays biotinylated protein from various *TurboID* fusion strains; some lane pairs correspond to a particular *TurboID* fusion in the *ZIP4+* or *zip4N919Q* genetic background (indicated in black across the top of the blot). In (B), independent membranes are separated by a vertical grey line, and the same blot is shown with computer-enhanced signal below the horizontal light blue line. Small blue circle shown above the 75 kDa marker indicates a band that migrates at the predicted position of Ecm11-TurboID. Representative blots are displayed; two or more biological replicates were examined for all strains. Some of the data informs the genetic dependency chart in Figure 3B.

Figure S3

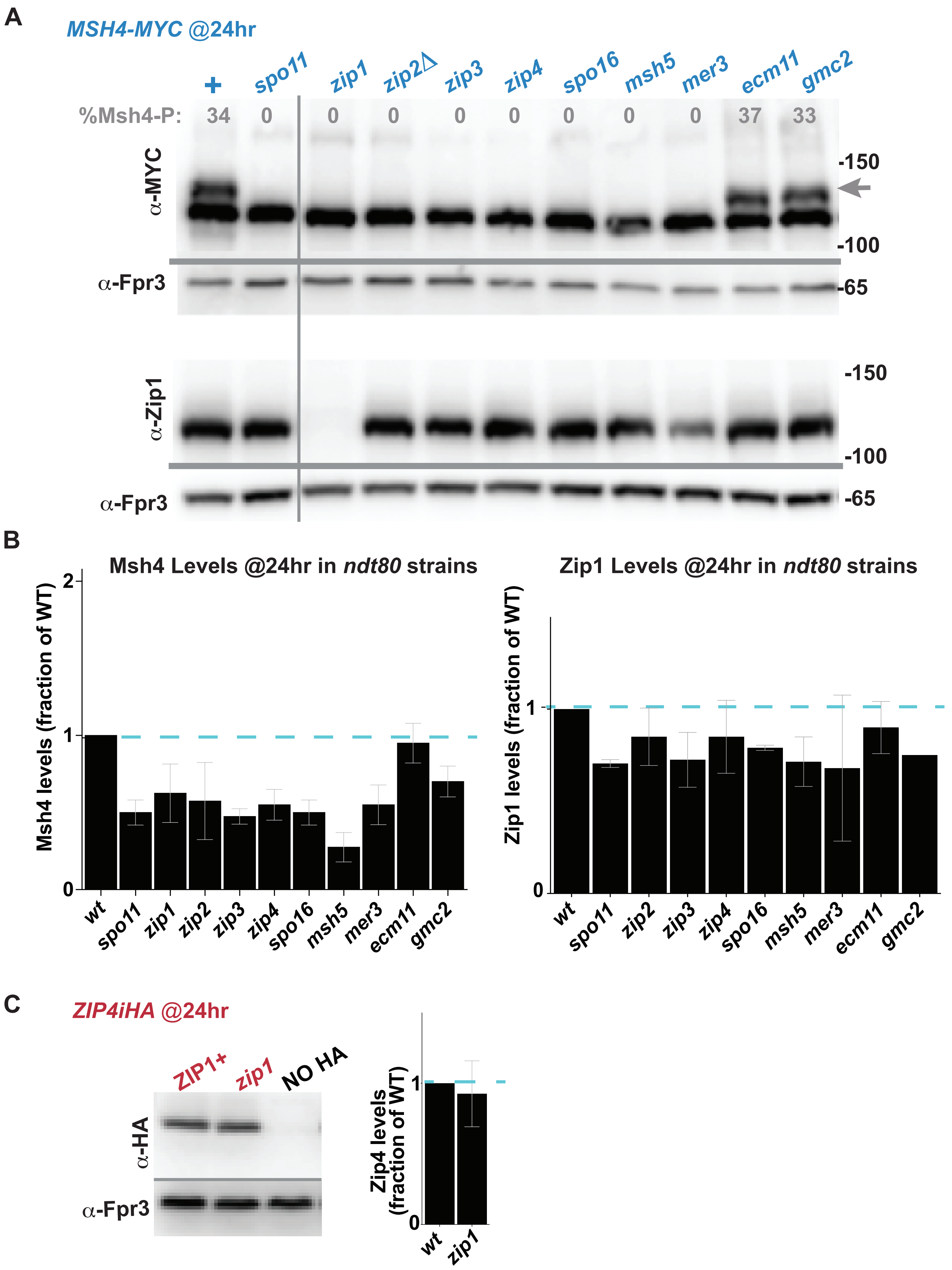
A
MSH4-MYC @24hr
+
spo11
zip1
zip2Δ
zip3
zip4
spo16
msh5
mer3
ecm11
gmc2
%Msh4-P:
34
0
0
0
0
0
0
0
0
37
33
α-MYC
-150
-100
α-Fpr3
-65
α-Zip1
-150
-100
α-Fpr3
-65
B
Msh4 Levels @24hr in ndt80 strains
Msh4 levels (fraction of WT)
2
1
0
wt
spo11
zip1
zip2
zip3
zip4
spo16
msh5
mer3
ecm11
gmc2
Zip1 Levels @24hr in ndt80 strains
Zip1 levels (fraction of WT)
1
0
wt
spo11
zip2
zip3
zip4
spo16
msh5
mer3
ecm11
gmc2
C
ZIP4iHA @24hr
ZIP1+
zip1
NO HA
α-HA
α-Fpr3
Zip4 levels (fraction of WT)
1
0
wt
zip1

**Figure S3. Msh4 and Zip1 levels in mutants missing meiotic recombination or synapsis components.** (A) Proteins extracted from *ndt80* cells arrested at mid-meiotic prophase were separated on two 8% polyacrylamide gels and transferred to nitrocellulose; membranes were then probed with either anti-MYC or anti-Zip1 antibodies. Blots show proteins from strains homozygous for *Msh4-MYC* and homozygous for one of several meiotic mutants (listed along the *x* axis above the blot). The slower migrating Msh4 species observed in the control and certain mutants corresponds to phosphorylated Msh4. Percentage of total Msh4-MYC that corresponds to the phosphorylated form is indicated above each lane (grey); values are an average of data from six blots, including multiple technical replicates of two biological replicates, with the following standard deviations: +=1.8; *spo11, zip1*, *zip2, zip3, zip4, spo16, msh5, mer3*=0; *ecm11*=2.1; *gmc2*=3.2. Fpr3 was detected on each blot (shown below grey line) and utilized as a loading control. Graphs in (B) plot the relative levels of Msh4-MYC and Zip1 protein in each strain, normalized using Fpr3; the level of Msh4-MYC and Zip1 protein in the control strain is set to one and indicated by a dotted blue line. Bars indicate standard deviations. The membrane in (C) was probed with anti-HA, anti-Fpr3 antibodies. Graph plots relative levels of Zip4iHA in each strain, normalized using Fpr3; level of Zip4iHA in the control is set to one, indicated by a dotted blue line.

Figure S4

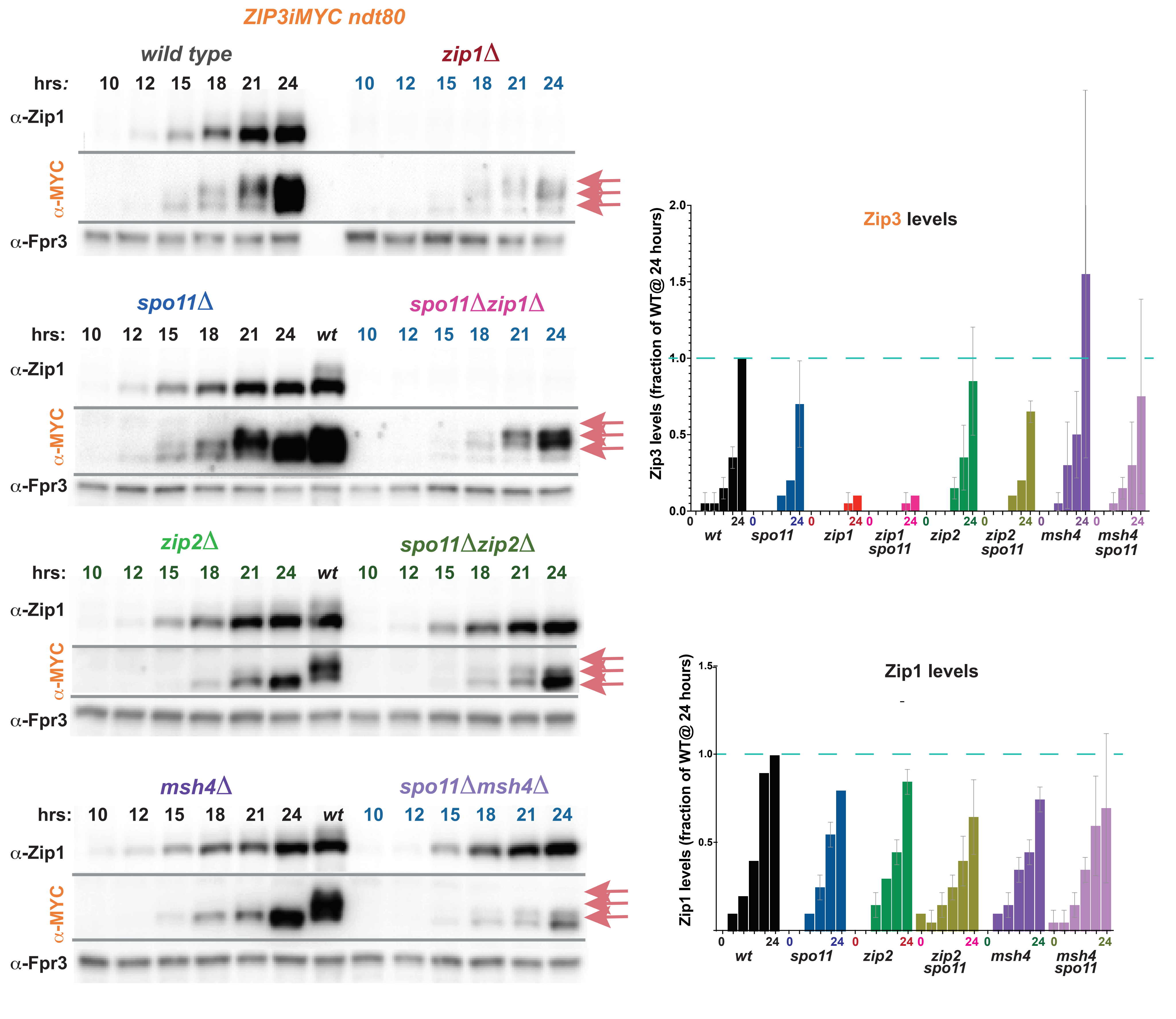
ZIP3iMYC ndt80
wild type
zip1Δ
hrs: 10 12 15 18 21 24
10 12 15 18 21 24
α-Zip1
α-MYC
α-Fpr3
spo11Δ
spo11Δzip1Δ
hrs: 10 12 15 18 21 24 wt 10 12 15 18 21 24
zip2Δ
spo11Δzip2Δ
msh4Δ
spo11Δmsh4Δ
Zip3 levels
Zip3 levels (fraction of WT@ 24 hours)
0.0
0.5
1.0
1.5
2.0
wt
spo11
zip1
zip1 spo11
zip2
zip2 spo11
msh4
msh4 spo11
Zip1 levels
Zip1 levels (fraction of WT@ 24 hours)
0.5
1.0
1.5
0
24

**Figure S4. Modified Zip3 species increase abundance in parallel with total Zip3 protein during meiotic prophase progression.** Western blots show protein extracted from *ndt80 Zip3iMYC* strains at different timepoints during meiotic prophase (10, 12, 15, 18, 21, and 24 hour timepoints are listed along the top of each blot), separated on an 8% polyacrylamide gel and transferred to nitrocellulose; the membranes were sequentially probed with anti-MYC, anti-Fpr3, and anti-Zip1 antibodies. Seven strains are homozygous for mutant alleles at additional loci: *zip1* (upper right), *spo11* (second row left), *spo11 zip1* (second row right), *zip2* (third row left), *spo11 zip2* (third row right), *msh4* (bottom row left) or *spo11 msh4* (bottom row right). Pink arrows mark the positions on the blot where Zip3 forms are detected. Note that lane seven in second, third and bottom row blots carries a wild type control, 24 hour sample for reference. Two biological replicates gave similar results. Graphs at the right plot abundance of total Zip3 (top) or Zip1 (bottom) protein at each of the six timepoints of the meiotic timecourse, relative to that found in control strains at the 24 hour timepoint (set to 1). Fpr3 is used as a loading control for normalization. Two biological replicates were used to evaluate protein levels, bars indicate standard deviations.

Figure S5

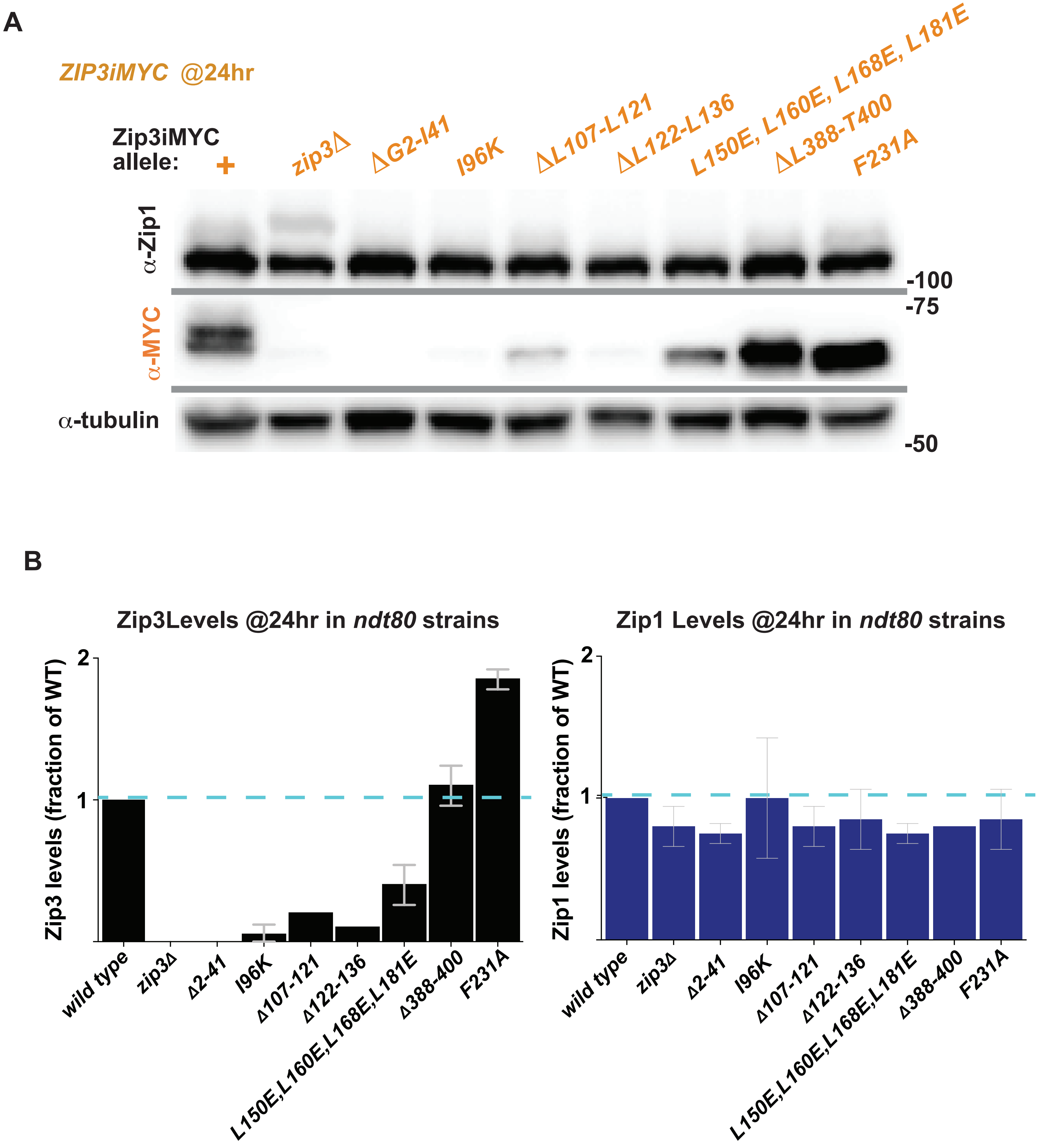
A
ZIP3iMYC @24hr
Zip3iMYC allele:
+
zip3Δ
ΔG2-I41
I96K
ΔL107-L121
ΔL122-L136
L150E, L160E, L168E, L181E
ΔL388-T400
F231A
α-Zip1
α-MYC
α-tubulin
-100
-75
-50
B
Zip3Levels @24hr in ndt80 strains
Zip3 levels (fraction of WT)
2
1
wild type
zip3Δ
Δ2-41
I96K
Δ107-121
Δ122-136
L150E,L160E,L168E,L181E
Δ388-400
F231A
Zip1 Levels @24hr in ndt80 strains
Zip1 levels (fraction of WT)
2
1
wild type
zip3Δ
Δ2-41
I96K
Δ107-121
Δ122-136
L150E,L160E,L168E,L181E
Δ388-400
F231A

**Figure S5. Several putative crossover-deficient *zip3* mutants generate unstable Zip3 protein.** (A) Western blot shows protein extracted from *ndt80* cells arrested at mid-meiotic prophase, separated on an 8% polyacrylamide gel; the membrane in (A) was sequentially probed with anti-MYC, anti-tubulin, and anti-Zip1 antibodies. Strains examined are homozygous for *ndt80* and *Zip3iMYC* and homozygous for one of several *zip3* internal deletion or point mutants (listed along the *x* axis above the blot). Graphs in (B) plot the relative levels of Zip3iMYC and Zip1 protein in each strain, normalized using tubulin; the level of Zip3iMYC and Zip1 protein in the control strain is set to one and indicated by a dotted blue line. Bars indicate standard deviations.

Figure S6

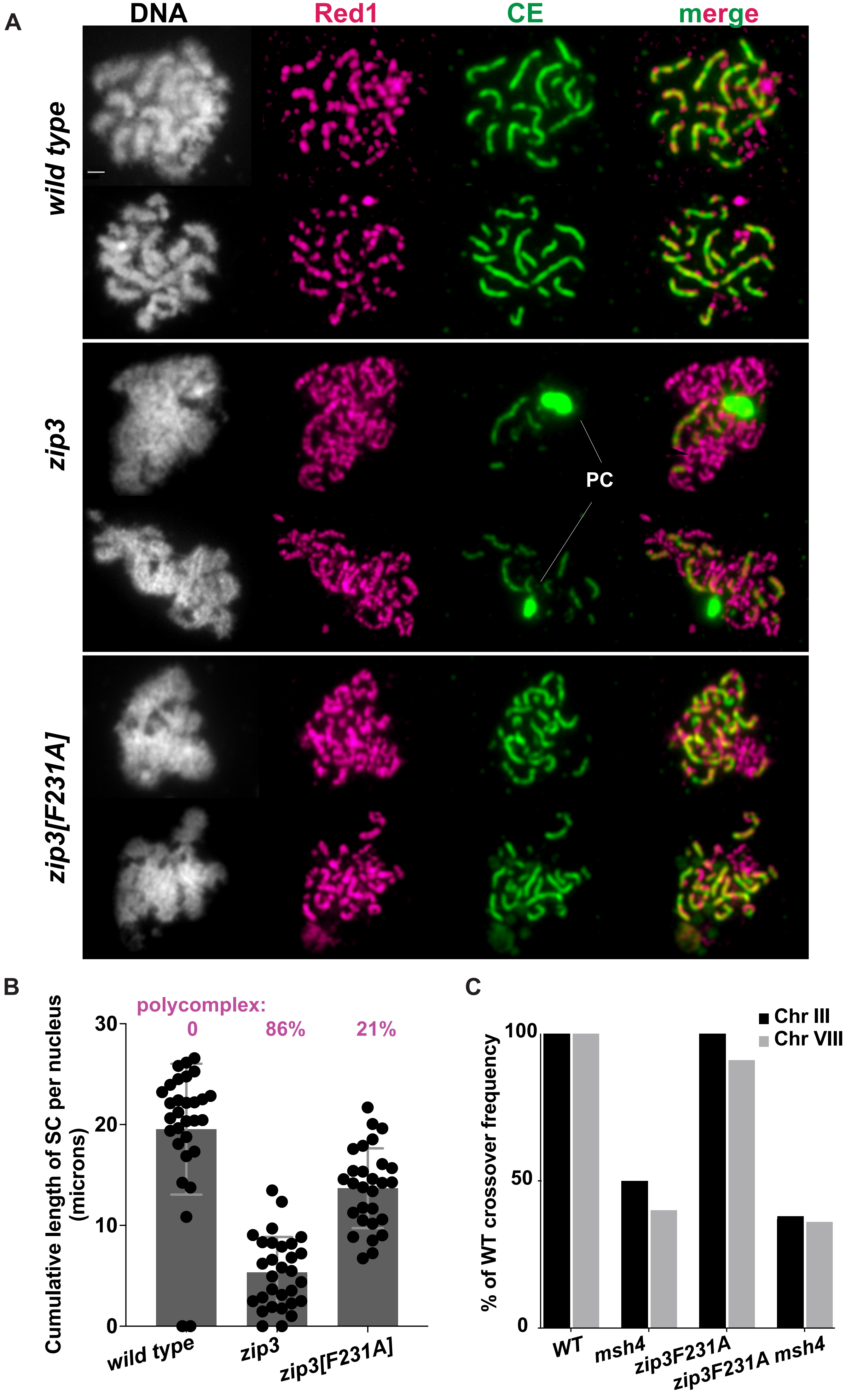
A
DNA
Red1
CE
merge
wild type
zip3
PC
zip3[F231A]
B
polycomplex:
0
86%
21%
Cumulative length of SC per nucleus (microns)
30
20
10
0
wild type
zip3
zip3[F231A]
C
Chr III
Chr VIII
% of WT crossover frequency
100
50
0
WT
msh4
zip3F231A
zip3F231A msh4

**Figure S6.** ***Zip3[F231A]* shows mild defects in MutSγ crossover recombination and synapsis.** Images in (A) show surface-spread meiotic chromosomes from *ndt80* strains sporulated for 24 hours. DAPI labels DNA (white), and immunofluorescence with anti-Red1 (magenta) and an antibody raised against the Ecm11-Gmc2 complex (CE; green) labels chromosome axes and the SC central element, respectively. Polycomplex is indicated (PC). Bar = 1 micron. Scatterplot in (B) gives the cumulative length of SC measured in microns per nucleus, (n=30 nuclei for wild type and *zip3*, 28 nuclei for *zip3[F231A])*. Shaded area for each strain indicates the average total length of SC per nucleus, and bars indicate standard deviation. Graph in (C) plots crossover recombination frequency in the *msh4,* the *zip3[F231A]* and the *zip3[F231A] msh4* strains as a percentage of wild type (100%, dotted blue line). Crossovers were measured in four intervals spanning most of the length of chromosome III (black bar) and in three intervals spanning over half of chromosome VIII (grey bar). Data are calculated from more than: 1000 tetrads for wild-type, 400 tetrads for *msh4*, 500 tetrads for *zip3[F231A]* and from ~100 tetrads for *zip3[F231A] msh4* (see Table S2). Wild-type and *msh4* data was previously published in (VOELKEL-MEIMAN *et al*. 2016; VOELKEL-MEIMAN 2021).

Figure S7

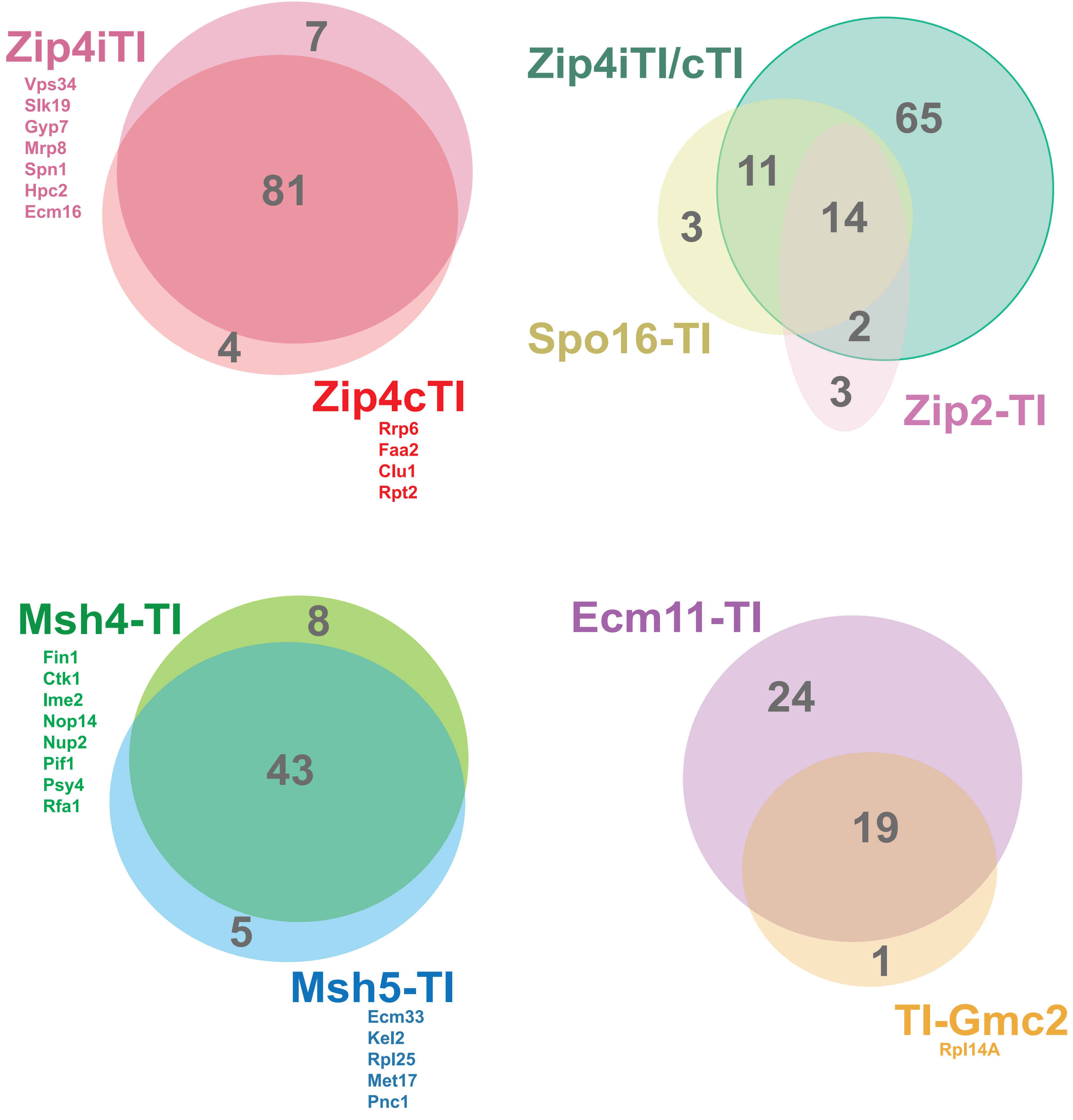

Zip4iTI
Vps34
Slk19
Gyp7
Mrp8
Spn1
Hpc2
Ecm16
7
81
4
Zip4cTI
Rrp6
Faa2
Clu1
Rpt2
Zip4iTI/cTI
65
11
14
3
2
3
Spo16-TI
Zip2-TI
Msh4-TI
Fin1
Ctk1
Ime2
Nop14
Nup2
Pif1
Psy4
Rfa1
8
43
5
Msh5-TI
Ecm33
Kel2
Rpl25
Met17
Pnc1
Ecm11-TI
24
19
1
TI-Gmc2
Rpl14A

**Figure S7. Proximity labeling targets of MutSγ pathway proteins substantially overlap.** Venn diagrams plot the shared targets, identified by mass spectrometry, of those TurboID fusion proteins that are either distinct fusions involving the same meiotic protein (Zip4iTI versus Zip4cTI) or are known to be part of a stable subcomplex with one another (members of the ZZS complex, the Msh4-Msh5 (MutSγ) heterodimer, or the Ecm11-Gmc2 heterocomplex). "TI" indicates TurboID. Number of targets that are either overlapping or specific to one component are indicated, respectively, in the overlapping or non-overlapping portion of the ovals. Targets specific to a given component (not considered overlapping) are absent in both biological replicates of the TurboID experiments corresponding to the other members of the complex. For comparisons between two proteins, non-overlapping targets are individually listed. See Table S3 for full list of proteins identified by mass spectrometry.

Figure S8

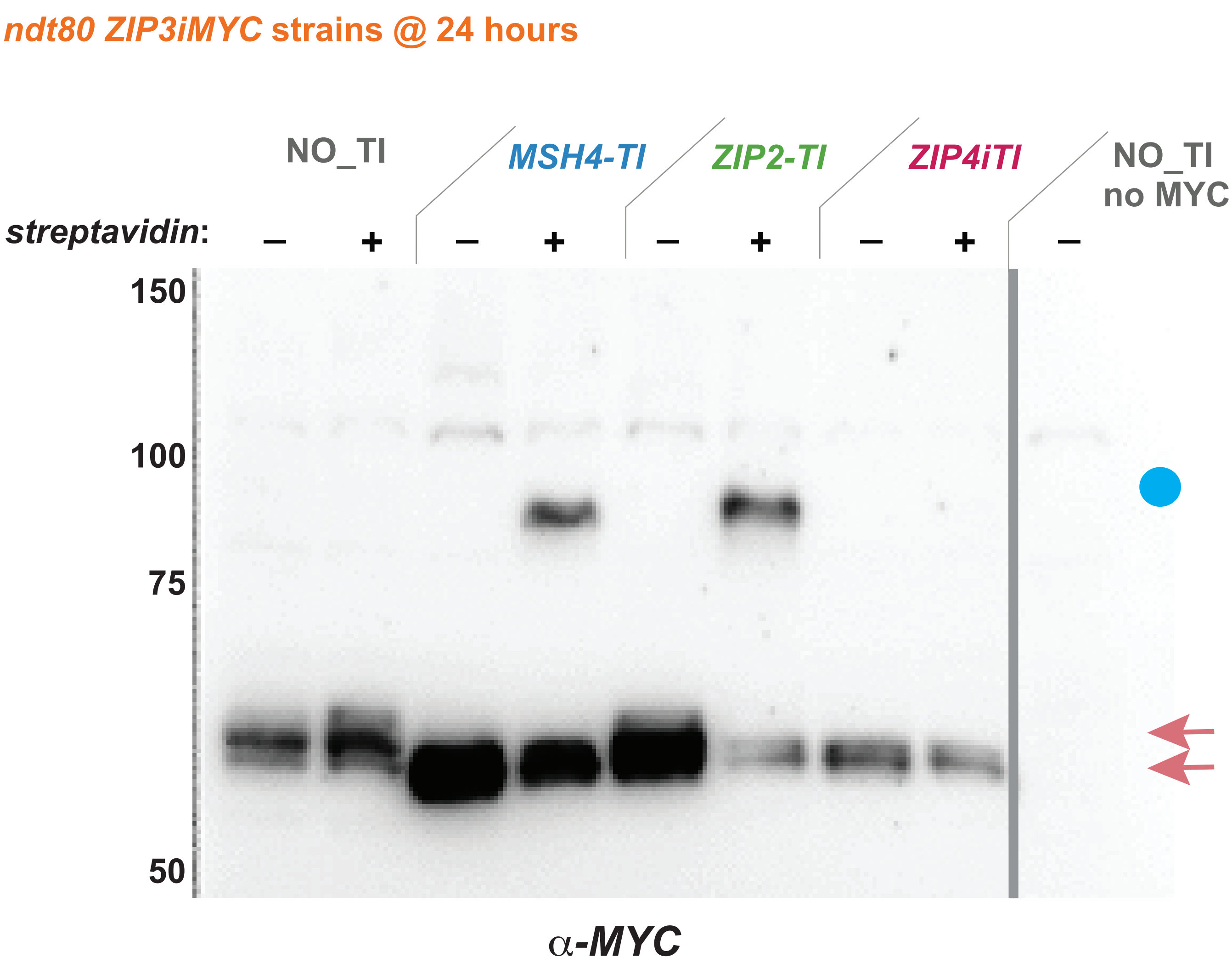

ndt80 ZIP3iMYC strains @ 24 hours
NO_TI
MSH4-TI
ZIP2-TI
ZIP4iTI
NO_TI
no MYC
streptavidin:
–
+
–
+
–
+
–
+
–
150
100
75
50
α-MYC

**Figure S8. Biotinylation of a specific protein can be observed with streptavidin pre-incubation followed by a traditional western blot.** Blot displays Zip3iMYC protein extracted from various *ndt80 ZIP3iMYC* strains homozygous for *MSH4-TurboID*, *ZIP2-TurboID*, or *ZIP4iTurboID*. The first lanes carry protein from a strain devoid of any *TurboID* fusion, and the last lane corresponds to a *ZIP3+ ndt80* (no MYC) control. Protein extracts were generated from cells arrested at mid-meiotic prophase, and proteins separated on an 8% polyacrylamide gel. The grey line indicates where the blot was cut to remove extraneous data. A "+" above the lane indicates that the protein sample was pre-incubated with streptavidin as described in (XIANG AND KOSHLAND 2021). Pink arrows correspond to the positions of Zip3iMYC protein unbound to streptavidin, while the blue dot corresponds to the position of streptavidin-bound, biotinylated Zip3iMYC.

**Table S1**
**Strains used in this study.** *Strains are of the BR1919-8B background (ROCKMILL AND ROEDER 1998).*

| | **GENOTYPE** |
|---|---|
| YAM1252 | *lys2ΔNhe his4-260,519 leu2-3,112 MAT**α** trp1-289 ura3-1 thr1-4 ade2-1*<br>*lys2ΔNhe his4-260,519 leu2-3,112 MAT**a** trp1-289 ura3-1 thr1-4 ade2-1* |
| JS21 | YAM1252 homozygous *ndt80::LEU2* |
| JS15 | YAM1252 homozygous *ZIP2-TurboID-3xMYC-kanMX6 ndt80::LEU2* |
| JS51 | JS15 homozygous *zip1::URA3* |
| AM4736 | JS15 homozygous *zip3::hphMX4* |
| AM4750 | JS15 homozygous *ZIP3-i3xMYC* (3x*MYC* inserted between amino acids 245/246) |
| AM5601 | JS15 homozygous *zip4::kanMX4* |
| AM4937 | JS15 homozygous *ecm11::hphMX4* |
| AM5024 | JS15 homozygous *ecm11-K5R,K101R* |
| AM5046 | JS15 homozygous *gmc2::hphMX4* |
| AM5332b | JS15 homozygous *rad51::hphMX4 dmc1::kanMX4* |
| AM4806 | JS15 homozygous *spo11::ADE2* |
| AM5304 | JS15 homozygous *spo16::hphMX4* |
| AM4935 | JS15 homozygous *msh4::ADE2* |
| AM5350 | JS15 homozygous *msh4::ADE2 ZIP3-i3xMYC* |
| AM5000 | JS15 homozygous *msh5::kanMX4* |
| KAV5-1 | JS15 homozygous *mer3::hphMX4* |
| AM6155 | JS15 homozygous *spo11::ADE2 ecm11::kanMX4* |
| AM5093 | JS15 homozygous *red1::hphMX4* |
| AM4747 | JS15 homozygous *zip1-F4A,F5A* |
| AM4746 | JS15 homozygous *zip1-N3A,R6A,D7A* |
| JS53 | JS15 homozygous *zip1[ΔM10-P14]* |
| AM4953 | JS15 homozygous *zip1[ΔR15-A20]* |
| AM5160 | JS15 homozygous *zip1[ΔK21-A163]* |
| AM4961 | JS15 homozygous *zip1[ΔR258-L278]* |
| AM4967 | JS15 homozygous *zip1[ΔN279-L296]* |
| AM5015 | JS15 homozygous *zip1[ΔM297-L317]* |
| AM5033 | JS15 homozygous *zip1[ΔS318-L327]* |
| AM4971 | JS15 homozygous *zip1[ΔI328-L354]* |
| AM5014 | JS15 homozygous *zip3[ΔG2-I41]* |
| AM4982 | JS15 homozygous *zip3-I96K* |
| AM4991 | JS15 homozygous *zip3[Δl107-L121]* |

| | |
|---|---|
| AM4987 | JS15 homozygous *zip3[ΔL122-L136]* |
| AM5034 | JS15 homozygous *zip3[ΔS137-L150]* |
| AM4772 | JS15 homozygous *zip3-L150E,L160E,L168E,L181E* |
| AM4980 | JS15 homozygous *zip3[ΔL388-T400]* |
| AM5261 | JS15 homozygous *zip3[ΔS203-R482]* |
| AM5347 | JS15 homozygous *zip3-F231A* |
| JS27 | YAM1252 homozygous *MSH4-TurboID-3xMYC-kanMX6 ndt80::LEU2* |
| JS83 | JS27 homozygous *zip1::URA3* |
| AM4928 | JS27 homozygous *zip2::hphMX4* |
| AM4742 | JS27 homozygous *zip3::hphMX4* |
| AM4752 | JS27 homozygous *ZIP3-i3xMYC* |
| AM5594a | JS27 homozygous *zip4::kanMX4* |
| AM4939 | JS27 homozygous *ecm11::hphMX4* |
| AM5040 | JS27 homozygous *ecm11-K5R,K101R* |
| AM4941 | JS27 homozygous *gmc2::hphMX4* |
| AM5334 | JS27 homozygous *rad51::hphMX4 dmc1::kanMX4* |
| AM4805 | JS27 homozygous *spo11::ADE2* |
| AM6075a | JS27 homozygous *spo11::kanMX4 ZIP3-i3xMYC* |
| AM4930 | JS27 homozygous *spo16::hphMX4* |
| AM4933 | JS27 homozygous *msh5::kanMX4* |
| AM5001 | JS27 homozygous *mer3::hphMX4* |
| AM6072 | JS27 homozygous *mer3::hphMX4 ZIP3-i3xMYC* |
| AM5002 | JS27 homozygous *red1::hphMX4* |
| AM4749 | JS27 homozygous *zip1-F4A,F5A* |
| AM4748 | JS27 homozygous *zip1-N3A,R6A,D7A* |
| JS85 | JS27 homozygous *zip1[ΔM10-P14]* |
| AM5043 | JS27 homozygous *zip1[ΔR15-A20]* |
| AM4735 | JS27 homozygous *zip1[ΔK21-A163]* |
| AM4962 | JS27 homozygous *zip1[ΔR258-L278]* |
| AM4968 | JS27 homozygous *zip1[ΔN279-L296]* |
| AM5039 | JS27 homozygous *zip1[ΔM297-L317]* |
| AM4970 | JS27 homozygous *zip1[ΔS318-L327]* |
| AM4972 | JS27 homozygous *zip1[ΔI328-L354]* |
| AM4986 | JS27 homozygous *zip3[ΔG2-I41]* |

| | |
|---|---|
| AM5027 | JS27 homozygous *zip3-I96K* |
| AM5075 | JS27 homozygous *zip3[Δl107-L121]* |
| AM5032 | JS27 homozygous *zip3[ΔS137-L150]* |
| AM4818 | JS27 homozygous *zip3-L150E,L160E,L168E,L181E* |
| AM4891 | JS27 homozygous *zip3[ΔL388-T400]* |
| AM5307c | JS27 homozygous *zip3[ΔS203-R482]* |
| AM4774 | JS27 homozygous *zip3-F231A* |
| K2303 | JS27 homozygous *zip4-N919Q* |
| AM5390 | YAM1252 homozygous *SPO16-TurboID-3xMYC-kanMX6 ndt80::LEU2* |
| K2175 | AM5390 homozygous *spo11::ADE2* |
| K2174 | AM5390 homozygous *zip1::URA3* |
| K2180 | AM5390 homozygous *zip2::hphMX4* |
| K2181 | AM5390 homozygous *zip3::URA3* |
| K2182 | AM5390 homozygous *zip4::kanMX4* |
| K2183 | AM5390 homozygous *mer3::hphMX4* |
| K2313 | AM5390 homozygous *ecm11::hphMX4* |
| K2312 | AM5390 homozygous *gmc2::hphMX4* |
| K2314 | AM5390 homozygous *msh4::ADE2* |
| K2304 | AM5390 homozygous *zip4-N919Q* |
| AM5649 | YAM1252 homozygous *ZIP4-iTurboID ndt80::LEU2* (*TurboID* inserted between amino acids 90/91) |
| K2237 | AM5649 homozygous *spo11::ADE2* |
| K2248 | AM5649 homozygous *ECM11-3xFLAG-kanMX4* |
| K2236 | AM5649 homozygous *zip1::URA3* |
| K2242 | AM5649 homozygous *zip2::hphMX4* |
| K2243 | AM5649 homozygous *zip3::URA3* |
| K2244 | AM5649 homozygous *spo16::hphMX4* |
| K2245 | AM5649 homozygous *mer3::hphMX4* |
| K2246 | AM5649 homozygous *ecm11::hphMX4* |
| K2247 | AM5649 homozygous *gmc2::hphMX4* |
| K2271 | AM5649 homozygous *ecm11-K5R,K101R* |
| K2249 | AM5649 homozygous *red1::hphMX4* |
| K2311 | YAM1252 homozygous *ZIP4-iTurboID-N919Q ndt80::LEU2* (*TurboID* inserted between amino acids 90/91) |
| JS32 | YAM1252 homozygous *MSH5-TurboID-3xMYC-kanMX6 ndt80::LEU2* |
| JS39 | JS32 homozygous *zip1::URA3* |

| | |
|---|---|
| AM5712 | YAM1252 homozygous *ECM11-TurboID-3xMYC-kanMX6 ndt80::LEU2* |
| K2198 | AM5712 homozygous *spo11::ADE2* |
| K2197 | AM5712 homozygous *zip1::URA3* |
| K2199 | AM5712 homozygous *zip2::hphMX4* |
| K2200 | AM5712 homozygous *zip3::URA3* |
| K2201 | AM5712 homozygous *zip4::kanMX4* |
| K2202 | AM5712 homozygous *spo16::hphMX4* |
| K2203 | AM5712 homozygous *mer3::hphMX4* |
| K2204 | AM5712 homozygous *gmc2::hphMX4* |
| K2425 | AM5712 homozygous *msh4::ADE2* |
| K2300 | AM5712 homozygous *zip4-N919Q* |
| AM5713 | AM5712 heterozygous *ECM11-TurboID-3xMYC-kanMX6* / *ECM11* |
| K2207 | AM5713 homozygous *spo11::ADE2* |
| K2206 | AM5713 homozygous *zip1::URA3* |
| K2208 | AM5713 homozygous *zip2::hphMX4* |
| K2209 | AM5713 homozygous *zip3::URA3* |
| K2210 | AM5713 homozygous *zip4::kanMX4* |
| K2211 | AM5713 homozygous *spo16::hphMX4* |
| K2212 | AM5713 homozygous *mer3::hphMX4* |
| K2213 | AM5713 homozygous *gmc2::hphMX4* |
| K2205 | AM5713 homozygous *ecm11-K5R,K101R* |
| K2426 | AM5713 homozygous *msh4::ADE2* |
| K2301 | AM5713 homozygous *zip4-N919Q* |
| K2269 | YAM1252 homozygous *ZIP3-iTurboID ndt80::LEU2* (*TurboID* inserted between amino acids 245/246) |
| K2160 | K2269 homozygous *zip1::URA3* |
| K2161 | K2269 homozygous *spo11::ADE2* |
| K2169 | K2269 homozygous *zip2::hphMX4* |
| K2170 | K2269 homozygous *zip4::kanMX4* |
| K2166 | K2269 homozygous *mer3::hphMX4* |
| K2316 | K2269 homozygous *ecm11::hphMX4* |
| K2315 | K2269 homozygous *gmc2::hphMX4* |
| K2423 | K2269 homozygous *spo16::hphMX4* |
| K2167 | K2269 homozygous *msh4::ADE2* |
| K2168 | K2269 homozygous *red1::hphMX4* |

| | |
|---|---|
| K2305 | K2269 homozygous *zip4-N919Q* |
| K2270 | K2269 heterozygous *ZIP3-iTurboID/ ZIP3* |
| K2185 | K2270 homozygous *zip1::URA3* |
| K2186 | K2270 homozygous *spo11::ADE2* |
| K2194 | K2270 homozygous *zip2::hphMX4* |
| K2195 | K2270 homozygous *zip4::kanMX4* |
| K2191 | K2270 homozygous *mer3::hphMX4* |
| K2356 | K2270 homozygous *ecm11::hphMX4* |
| K2355 | K2270 homozygous *gmc2::hphMX4* |
| K2424 | K2270 homozygous *spo16::hphMX4* |
| K2192 | K2270 homozygous *msh4::ADE2* |
| K2193 | K2270 homozygous *red1::hphMX4* |
| AM5392a | YAM1252 homozygous *ZIP4-TurboID-3xMYC-kanMX6 ndt80::LEU2* |
| AM5397a | YAM1252 homozygous *MER3-TurboID-3xMYC-kanMX6 ndt80::LEU2* |
| AM5396a | YAM1252 homozygous *MLH3-TurboID-3xMYC-kanMX6 ndt80::LEU2* |
| AM5828 | YAM1252 homozygous *TurboID-GMC2 ndt80::LEU2* |
| AM5829 | YAM1252 *TurboID-GMC2/GMC2 ndt80::LEU2* |
| K842 | *lys2ΔNhe HIS4 leu2-3,112 hphMX4@CEN3 MAT**α** ADE2@RAD18 natMX4@HMR*<br>*lys2ΔNhe his4-260,519 leu2-3,112 CEN3 MAT**a** RAD18 HMR*<br>*trp1-289 ura3-1 TRP1MX4@SPO11 spo13::URA3 THR1 210kb ade2-1*<br>*trp1-289 ura3-1 SPO11 SPO13 thr1-4 LYS2@210kb ade2-1* |
| K852 | K842 homozygous *msh4::kanMX4* |
| K1309 | K842 homozygous *zip1-F4A,F5A* |
| K1281 | K842 homozygous *zip1-N3A,R6A,D7A* |
| SYC107 | K842 homozygous *zip1[ΔM10-P14] thr1-4 LEU2@152kbXI 193kb XI*<br>*152kb XI THR1@193kb* |
| AF8 | K842 homozygous *zip1[ΔR15-A20]* |
| AF6 | K842 homozygous *zip1[ΔK21-A163]* |
| LY674 | K842 homozygous *zip1[ΔR258-L278]* |
| LY582 | K842 homozygous *zip1[ΔN279-L296]* |
| LY579 | K842 homozygous *zip1[ΔM297-L317]* |
| LY583 | K842 homozygous *zip1[ΔS318-L327]* |
| LY584 | K842 homozygous *zip1[ΔI328-L354]* |
| AP204 | K842 homozygous *zip3-F231A* |
| AP207 | K842 homozygous *zip3-F231A msh4::kanMX4* |
| K1268 | YAM1252 homozygous *MSH4-13xMYC-kanMX4 ndt80::LEU2* |

| | |
|---|---|
| AM4263 | K1268 homozygous *zip1::URA3* |
| K1840 | K1268 homozygous *zip1-F4A,F5A* |
| K1841 | K1268 homozygous *zip1-N3A,R6A,D7A* |
| AM4269 | K1268 homozygous *zip1[ΔM10-P14]* |
| AM4264 | K1268 homozygous *zip1[ΔR15-A20]* |
| K1838 | K1268 homozygous *zip1[ΔK21-A163]* |
| K1849 | K1268 homozygous *zip1[ΔN279-L296]* |
| K1850 | K1268 homozygous *zip1[ΔM297-L317]* |
| K1851 | K1268 homozygous *zip1[ΔI328-L354]* |
| AP184 | K1268 homozygous *zip3::hphMX4* |
| AM4689 | K1268 homozygous *zip3[ΔG2-I41]* |
| AP189 | K1268 homozygous *zip3-I96K* |
| AM4690 | K1268 homozygous *zip3[Δl107-L121]* |
| AM4691 | K1268 homozygous *zip3[ΔL122-L136]* |
| AM4692 | K1268 homozygous *zip3[ΔS137-L150]* |
| AM4696 | K1268 homozygous *zip3-L150E,L160E,L168E,L181E* |
| AM4692 | K1268 homozygous *zip3[ΔL388-T400]* |
| AM4695 | K1268 homozygous *zip3-F231A* |
| AM6201 | K1268 homozygous *zip3-F231A-i3xMYC* (3x*MYC* inserted between amino acids 245/246) |
| AM6202 | K1268 homozygous *ZIP3-i3xMYC* |
| AM5329 | K1268 homozygous *spo11::ADE2* |
| AM4263 | K1268 homozygous *zip1::URA3* |
| K2003 | K1268 homozygous *zip2::hphMX4* |
| K2006 | K1268 homozygous *zip4::hphMX4* |
| K2000 | K1268 homozygous *spo16::hphMX4* |
| AM4510 | K1268 homozygous *msh5::kanMX4* |
| AM4506 | K1268 homozygous *mer3::hphMX4* |
| AM4524 | K1268 homozygous *ecm11::kanMX4* |
| K1988 | K1268 homozygous *gmc2::hphMX4* |
| K1770 | YAM1252 homozygous *ZIP3-i3xMYC ndt80::LEU2* |
| K1884 | K1770 homozygous *spo11::ADE2* |
| K1771 | K1770 homozygous *zip1::URA3* |
| K1976 | K1770 homozygous *zip2::hphMX4* |
| K1979 | K1770 homozygous *zip4::hphMX4* |

| | |
|---|---|
| K1973 | K1770 homozygous *spo16::hphMX4* |
| AM5351 | K1770 homozygous *msh4::ADE2* |
| K1958 | K1770 homozygous *msh5::hphMX4* |
| K1961 | K1770 homozygous *mer3::hphMX4* |
| K1970 | K1770 homozygous *red1::hphMX4* |
| K1952 | K1770 homozygous *ecm11::hphMX4* |
| K1955 | K1770 homozygous *gmc2::hphMX4* |
| K2461 | K1770 homozygous *zip4-N919Q* |
| K1772 | K1770 homozygous *zip1::URA3/ ZIP1* |
| K1777 | K1770 homozygous *zip1-F4A,F5A* |
| K1778 | K1770 homozygous *zip1-N3A,R6A,D7A* |
| K1774 | K1770 homozygous *zip1[ΔS2-S9]* |
| K1775 | K1770 homozygous *zip1[ΔM10-P14]* |
| K1776 | K1770 homozygous *zip1[ΔR15-A20]* |
| K1773 | K1770 homozygous *zip1[ΔK21-A163]* |
| K1791 | K1770 homozygous *zip1[ΔN279-L296]* |
| K1792 | K1770 homozygous *zip1[ΔM297-L317]* |
| K1793 | K1770 homozygous *zip1[ΔS318-L327]* |
| K1794 | K1770 homozygous *zip1[ΔI328-L354]* |
| AM6122 | K1770 homozygous *spo11::ADE2 zip1::URA3* |
| AM6148 | K1770 homozygous *spo11::ADE2 zip2::hphMX4* |
| AM6149 | K1770 homozygous *spo11::ADE2 msh4::kanMX4* |
| AM5820 | K1770 homozygous *zip1::URA3 zip2::hphMX4* |
| AM5819 | K1770 homozygous *zip1::URA3 msh4::ADE2* |
| K2013 | K1770 homozygous *zip3-F231A-i3xMYC* (3x*MYC* inserted between amino acids 245/246) |
| AM6163 | K2013 homozygous *zip1::URA3* |
| AM6161 | K2013 homozygous *zip1::URA3 spo11::ADE2* |
| AP148 | K2013 homozygous *spo11::ADE2* |
| K2009 | K1770 homozygous *zip3[ΔG2-I41]-i3xMYC* (3x*MYC* inserted between amino acids 245/246) |
| K2014 | K1770 homozygous *zip3-I96K-i3xMYC* (3x*MYC* inserted between amino acids 245/246) |
| K2010 | K1770 homozygous *zip3[Δl107-L121]-i3xMYC* (3x*MYC* inserted between amino acids 245/246) |
| K2008 | K1770 homozygous *zip3[ΔL122-L136]-i3xMYC* (3x*MYC* inserted between amino acids 245/246) |
| K2011 | K1770 homozygous *zip3-L150E,L160E,L168E,L181E-i3xMYC* (3x*MYC* inserted between amino acids 245/246) |

| | |
|---|---|
| K2015 | K1770 homozygous *zip3-i3xMYC-[ΔL388-T400]* (3x*MYC* inserted between amino acids 245/246) |
| K1795 | YAM1252 homozygous *ZIP4-i3xHA ndt80::LEU2* (*3xHA* inserted between amino acids 90/91) |
| K1796 | K1795 *zip1::URA3* |
| AM6257 | YAM1252 *ZIP4-i3xHA*/*ZIP4* homozygous *ZIP3-i3xMYC ndt80::LEU2* |
| AM6259 | AM6257 homozygous *mer3::hphMX4* |
| AM6071 | YAM1252 *ZIP3-i3xMYC*/*ZIP3* homozygous *MSH4-3xHA-kanMX4 ndt80::LEU2* |
| AM6059 | AM6071 homozygous *mer3::hphMX4* |
| AP313 | YAM1252 homozygous *ndt80::LEU2* (same as JS21) |
| AP314 | AP313 homozygous *zip3::URA3* |
| AP320 | YAM1252 homozygous *zip3-F231A* |

**Table S2. Crossover frequency of select *zip1* and *zip3* alleles**

| GENOTYPE (STRAIN) | INTERVAL (CHROMOSOME) | PD | TT | NPD | TOTAL | cM (± SE) | %WT | cM by chrm | %WT by chrm | NPDobs/NPDexp (± SE) | viability |
|---|---|---|---|---|---|---|---|---|---|---|---|
| *WT**** | *HIS4-CEN3* (III) | 584 | 528 | 10 | 1122 | **26.2 (1.1)** | **100** | 103.7 (III) | **100** | 0.20 (0.07) | |
| (K842) | *CEN3-MAT* (III) | 708 | 416 | 4 | 1128 | **19.5 (0.9)** | **100** | | | 0.15 (0.08) | **97%** |
| | *MAT-RAD18* (III) | 412 | 676 | 18 | 1106 | **35.4 (1.3)** | **100** | | | 0.16 (0.04) | |
| | *RAD18-HMR* (III) | 651 | 454 | 8 | 1113 | **22.6 (1.0)** | **100** | | | 0.24 (0.09) | |
| | *SPO11-SPO13* (VIII) | 453 | 630 | 33 | 1116 | **37.1 (1.6)** | **100** | 75.2 (VIII) | **100** | 0.40 (0.07) | |
| | *SPO13-THR1* (VIII) | 913 | 180 | 2 | 1095 | **8.8 (0.7)** | **100** | | | 0.48 (0.34) | |
| | *THR1-LYS2* (VIII) | 490 | 590 | 8 | 1088 | **29.3 (1.0)** | **100** | | | 0.11 (0.04) | |
| *msh4Δ** | *HIS4-CEN3* (III) | 375 | 96 | 1 | 472 | **10.8 (1.1)** | **41** | 53.4 (III) | **51** | 0.35 (0.35) | |
| (K852) | *CEN3-MAT* (III) | 425 | 51 | 1 | 477 | **6.0 (0.9)** | **31** | | | 1.36 (1.36) | **71%** |
| | *MAT-RAD18* (III) | 276 | 184 | *7* | 467 | **24.2 (1.9)** | **68** | | | 0.55 (0.21) | |
| | *RAD18-HMR* (III) | 352 | 116 | 0 | 468 | **12.4 (1.0)** | **55** | | | ND | |
| | *SPO11-SPO13* (VIII) | 365 | 89 | 3 | 457 | **11.7 (1.4)** | **32** | 30.4 (VIII) | **40** | 1.20 (0.70) | |
| | *SPO13-THR1* (VIII) | 423 | 27 | 0 | 450 | **3.0 (0.6)** | **34** | | | ND | |
| | *THR1-LYS2* (VIII) | 319 | 129 | 2 | 450 | **15.7 (1.4)** | **54** | | | 0.34 (0.24) | |
| *zip1-F4A,F5A*** | *HIS4-CEN3* (III) | 297 | 114 | 3 | 414 | **15.9 (1.6)** | **61** | 72.4 (III) | **70** | 0.61 (0.36) | |
| (K1309) | *CEN3-MAT* (III) | 300 | 114 | 2 | 416 | **15.1 (1.5)** | **77** | | | 0.41 (0.29) | **87%** |
| | *MAT-RAD18* (III) | 260 | 148 | 6 | 414 | **22.2 (2.0)** | **63** | | | 0.67 (0.28) | |
| | *RAD18-HMR* (III) | 272 | 142 | 3 | 417 | **19.2 (1.7)** | **85** | 29.2 (VIII) | **39** | 0.38 (0.38) | |
| | *SPO13-THR1* (VIII) | 373 | 34 | 0 | 407 | **4.2 (0.7)** | **48** | | | ND | |
| | *THR1-LYS2* (VIII) | 304 | 101 | 2 | 407 | **13.9 (1.5)** | **47** | | | 0.52 (0.38) | |
| *zip1-N3A,R6A,D7A*** | *HIS4-CEN3* (III) | 304 | 219 | 3 | 526 | **22.5 (1.4)** | **86** | 83.1 (III) | **80** | 0.18 (0.11) | |
| (K1281) | *CEN3-MAT* (III) | 367 | 156 | 1 | 524 | **15.5 (1.1)** | **79** | | | 0.14 (0.14) | **93%** |
| | *MAT-RAD18* (III) | 264 | 255 | 5 | 524 | **27.2 (1.6)** | **77** | | | 0.20 (0.09) | |
| | *RAD18-HMR* (III) | 354 | 171 | 3 | 528 | **17.9 (1.4)** | **79** | | | 0.33 (0.19) | |
| | *SPO11-SPO13* (VIII) | 289 | 223 | 5 | 517 | **24.5 (1.6)** | **66** | 51.8 (VIII) | **69** | 0.28 (0.13) | |
| | *SPO13-THR1* (VIII) | 462 | 52 | 0 | 514 | **5.1 (0.7)** | **58** | | | ND | |
| | *THR1-LYS2* (VIII) | 292 | 224 | 1 | 517 | **22.2 (1.2)** | **76** | | | 0.06 (0.06) | |
| *zip1[ΔM10-P14]*** | HIS4-CEN3 (III) | 411 | 139 | 5 | 555 | **15.2 (1.5)** | **54** | | | 0.94 (0.43) | **92%** |
| (SYC107) | CEN3-MAT (III) | 443 | 119 | 5 | 567 | **13.1 (1.4)** | **75** | 68.5 (III) | **64** | 1.37 (0.62) | |
| | MAT-RAD18 (III) | 336 | 219 | 5 | 560 | **22.2 (1.5)** | **57** | | | 0.33 (0.15) | |
| | RAD18-HMR (III) | 365 | 196 | 1 | 562 | **18.0 (1.1)** | **79** | | | n.d. | |
| | SPO11-SPO13 (VIII) | 399 | 149 | 2 | 550 | **14.6 (1.2)** | **48** | (VIII) | **48** | 0.32 (0.23) | |
| | **iTHR1-iLEU2 (XI)** | 515 | 47 | 0 | 562 | **4.2 (0.6)** | **46** | | | n.d. | |
| *zip1[ΔR15-A20]*** | *HIS4-CEN3* (III) | 409 | 139 | 4 | 552 | **14.8 (1.4)** | **56** | 84.2 (III) | **81** | 0.75 (0.38) | |
| (AF8) | *CEN3-MAT* (III) | 388 | 172 | 7 | 567 | **18.9 (1.6)** | **97** | | | 0.84 (0.32) | **87%** |
| | *MAT-RAD18* (III) | 304 | 232 | 14 | 550 | **28.7 (2.1)** | **81** | | | 0.78 (0.22) | |
| | *RAD18-HMR* (III) | 348 | 199 | 7 | 554 | **21.8 (1.7)** | **96** | | | 0.58 (0.22) | |
| | *SPO11-SPO13* (VIII) | 380 | 169 | 6 | 555 | **18.5 (1.6)** | **50** | 55.0 (VIII) | **73** | 0.73 (0.30) | |
| | *SPO13-THR1* (VIII) | 434 | 89 | 0 | 523 | **8.5 (0.8)** | **97** | | | NA | |
| | *THR1-LYS2* (VIII) | 275 | 239 | 9 | 523 | **28.0 (1.9)** | **96** | | | 0.43 (0.15) | |
| *zip1[Δ21-163]*** | *HIS4-CEN3* (III) | 263 | 311 | 10 | 584 | **31.8 (1.8)** | **121** | 139.9 (III) | **135** | 0.28 (0.09) | |
| (AF6) | *CEN3-MAT* (III) | 242 | 331 | 13 | 586 | **34.9 (1.9)** | **179** | | | 0.30 (009) | **89%** |
| | *MAT-RAD18* (III) | 208 | 329 | 17 | 554 | **38.9 (2.2)** | **110** | | | 0.35 (0.09) | |
| | *RAD18-HMR* (III) | 231 | 320 | 11 | 562 | **34.3 (1.9)** | **152** | | | 0.26 (0.08) | |
| | *SPO11-SPO13* (VIII) | 182 | 332 | 44 | 558 | **53.4 (3.2)** | **144** | 124.5 (VIII) | **166** | 0.88 (0.16) | |
| | *SPO13-THR1* (VIII) | 325 | 194 | 3 | 522 | **20.3 (1.4)** | **231** | | | 0.24 (0.14) | |
| | *THR1-LYS2* (VIII) | 162 | 329 | 34 | 525 | **50.8 (3.0)** | **173** | | | 0.59 (0.12) | |
| *zip1[ΔR258-L278]* | *HIS4-CEN3* (III) | 42 | 65 | 1 | 108 | **32.9 (3.4)** | **126** | 111.4 (III) | **107** | 0.10 (0.10) | |
| (LY674) | *CEN3-MAT* (III) | 76 | *35* | 0 | 111 | **15.9 (2.2)** | **82** | | | ND | **97%** |
| | *MAT-RAD18* (III) | 34 | 71 | 2 | 107 | **38.8 (4.1)** | **110** | | | 0.12 (0.10) | |

| | | | | | | | | | | | |
|---|---|---|---|---|---|---|---|---|---|---|---|
| | *RAD18-HMR* (III) | 61 | 45 | 1 | 107 | **23.8 (3.5)** | **105** | | | 0.29 (0.29) | |
| | *SPO11-SPO13* (VIII) | 47 | 59 | 3 | 109 | **35.3 (4.9)** | **95** | 77.1 (VIII) | **103** | 0.42 (0.26) | |
| | *SPO13-THR1* (VIII) | 87 | 19 | 1 | 107 | **11.7 (3.3)** | **133** | | | 2.08 (2.11) | |
| | *THR1-LYS2* (VIII) | 43 | 65 | 0 | 108 | **30.1 (2.4)** | **103** | | | ND | |
| *zip1[ΔN279-L296]* (LY582) | *HIS4-CEN3* (III) | 192 | 300 | 17 | 509 | **39.5 (2.4)** | **151** | 158.6 (III) | **153** | 0.39 (0.10) | |
| | *CEN3-MAT* (III) | 141 | 374 | 11 | 526 | **41.8 (1.9)** | **214** | | | 0.34 (0.04) | **89%** |
| | *MAT-RAD18* (III) | 171 | 313 | 25 | 509 | **45.5 (2.8)** | **129** | | | 0.48 (0.11) | |
| | *RAD18-HMR* (III) | 230 | 270 | 9 | 509 | **31.8 (1.9)** | **141** | | | 0.29 (0.10) | |
| | *SPO11-SPO13* (VIII) | 130 | 329 | 40 | 499 | **57.0 (3.4)** | **154** | 117.3 (VIII) | **156** | 0.55 (0.13) | |
| | *SPO13-THR1* (VIII) | 327 | 152 | 2 | 481 | **17.1 (1.4)** | **194** | | | 0.26 (0.18) | |
| | *THR1-LYS2* (VIII) | 135 | 332 | 14 | 481 | **43.2 (2.3)** | **147** | | | 0.29 (0.03) | |
| *zip1[ΔM297-L317]* (LY579) | *HIS4-CEN3* (III) | 399 | 118 | 2 | 519 | **12.5 (1.2)** | **48** | 71.5 (III) | **69** | 0.50 (0.36) | |
| | *CEN3-MAT* (III) | 388 | 134 | 3 | 525 | **14.5 (1.3)** | **74** | | | 0.57 (0.33) | **84%** |
| | *MAT-RAD18* (III) | 308 | 184 | 7 | 499 | **22.7 (1.8)** | **64** | | | 0.60 (0.23) | |
| | *RAD18-HMR* (III) | 332 | 167 | 9 | 508 | **21.8 (2.0)** | **96** | | | 1.00 (0.34) | |
| | *SPO11-SPO13* (VIII) | 323 | 168 | 15 | 506 | **25.5 (2.4)** | **69** | 66.1 (VIII) | **88** | 1.63 (0.44) | |
| | *SPO13-THR1* (VIII) | 373 | 106 | 1 | 480 | **11.7 (1.1)** | **133** | | | 0.29 (0.29) | |
| | *THR1-LYS2* (VIII) | 263 | 206 | 12 | 481 | **28.9 (2.3)** | **99** | | | 0.73 (0.22) | |
| *zip1[ΔS318-L327]* (LY583) | *HIS4-CEN3* (III) | 377 | 100 | 2 | 479 | **11.7 (1.3)** | **45** | 71.5 (VIII) | **69** | 0.65 (0.47) | |
| | *CEN3-MAT* (III) | 363 | 118 | 3 | 484 | **14.1 (1.4)** | **72** | | | 0.69 (0.40) | **81%** |
| | *MAT-RAD18* (III) | 271 | 194 | 7 | 472 | **25.0 (1.9)** | **71** | | | 0.49 (0.19) | |
| | *RAD18-HMR* (III) | 306 | 159 | 6 | 471 | **20.7 (1.8)** | **92** | | | 0.67 (0.28) | |
| | *SPO11-SPO13* (VIII) | 354 | 113 | 3 | 470 | **13.9 (1.4)** | **37** | 45.3 (VIII) | **60** | 0.73 (0.43) | |
| | *SPO13-THR1* (VIII) | 399 | 54 | 0 | 453 | **6.0 (0.8)** | **68** | | | ND | |
| | *THR1-LYS2* (VIII) | 249 | 201 | 5 | 455 | **25.4 (1.8)** | **87** | | | 0.30 (0.14) | |
| *zip1[ΔI328-L354]* (LY584) | *HIS4-CEN3* (III) | 363 | 137 | 4 | 504 | **16.0 (1.5)** | **61** | 81.7 (III) | **79** | 0.69 (0.35) | |
| | *CEN3-MAT* (III) | 344 | 168 | 1 | 513 | **17.0 (1.2)** | **87** | | | 0.11 (0.11) | **90%** |
| | *MAT-RAD18* (III) | 266 | 231 | 12 | 509 | **29.8 (2.2)** | **84** | | | 0.60 (0.18) | |
| | *RAD18-HMR* (III) | 341 | 162 | 5 | 508 | **18.9 (1.6)** | **84** | | | 0.60 (0.27) | |
| | *SPO11-SPO13* (VIII) | 343 | 146 | 4 | 493 | **17.2 (1.5)** | **46** | 46.3 (VIII) | **62** | 0.58 (0.30) | |
| | *SPO13-THR1* (VIII) | 417 | 70 | 0 | 487 | **7.2 (0.8)** | **82** | | | ND | |
| | *THR1-LYS2* (VIII) | 289 | 195 | 3 | 487 | **21.9 (1.5)** | **75** | | | 0.22 (0.13) | |
| *zip3-F231A* (AP204) | *HIS4-CEN3* (III) | 298 | 222 | 4 | 524 | **23.5 (1.5)** | **90** | 104.2 (III) | **100** | 0.23 (0.12) | |
| | *CEN3-MAT* (III) | 306 | 216 | 1 | 523 | **21.2 (1.2)** | **109** | | | 0.06 (0.06) | **95%** |
| | *MAT-RAD18* (III) | 208 | 290 | 14 | 512 | **36.5 (2.2)** | **103** | | | 0.36 (0.11) | |
| | *RAD18-HMR* (III) | 298 | 224 | 3 | 525 | **23.0 (1.4)** | **102** | | | 0.17 (0.10) | |
| | *SPO11-SPO13* (VIII) | 277 | 239 | 7 | 523 | **26.9 (1.8)** | **73** | 68.3 (VIII) | **91** | 0.33 (0.14) | |
| | *SPO13-THR1* (VIII) | 431 | 89 | 0 | 520 | **8.6 (0.8)** | **98** | | | ND | |
| | *THR1-LYS2* (VIII) | 239 | 268 | 12 | 519 | **32.8 (2.1)** | **112** | | | 0.41 (0.13) | |
| *zip3-F231A msh4Δ* (AP207) | *HIS4-CEN3* (III) | 94 | 14 | 0 | 108 | **6.5 (1.6)** | **25** | 38.9 (III) | **38** | ND | |
| | *CEN3-MAT* (III) | 94 | 14 | 0 | 108 | **6.5 (1.6)** | **33** | | | ND | **65%** |
| | *MAT-RAD18* (III) | 66 | 40 | 0 | 106 | **18.9 (2.4)** | **56** | | | ND | |
| | *RAD18-HMR* (III) | 93 | 13 | 0 | 106 | **6.1 (1.6)** | **27** | | | ND | |
| | *SPO11-SPO13* (VIII) | 85 | 15 | 0 | 100 | **7.5 (1.8)** | **20** | 27.3 (VIII) | **36** | ND | |
| | *SPO13-THR1* (VIII) | 98 | 8 | 0 | 106 | **3.8 (1.3)** | **43** | | | ND | |
| | *THR1-LYS2* (VIII) | 82 | 22 | 2 | 106 | **16.0 (4.3)** | **55** | | | 2.99 (2.51) | |

Data previously published:

* VOELKEL-MEIMAN et al. 2016

** VOELKEL-MEIMAN et al. 2019

*** VOELKEL-MEIMAN et al. 2021

**Table S2. Crossover frequency of select *zip1* and *zip3* alleles.** Map distances and genetic interference values were calculated using tetrad analysis or random spore analysis and coefficient of coincidence measurements as described previously (VOELKEL-MEIMAN *et al.* 2013; VOELKEL-MEIMAN *et al.* 2015; VOELKEL-MEIMAN *et al.* 2019). Table gives map distances and their corresponding percentages of the wild-type values for individual intervals, and for the entire chromosome (by summing the intervals on III or VIII). For intervals marked (ND), interference measurements are not obtainable using the coefficient of coincidence method due to an absence of NPD tetrads. Crossover frequencies for strains marked with an *, **, *** were previously published (VOELKEL-MEIMAN *et al.* 2016; VOELKEL-MEIMAN *et al.* 2019; VOELKEL-MEIMAN 2021, respectively). Data is plotted on the graph in Figures 5 and S6.

| Table S3. Streptavidin-purified proteins identified using UPLC-M | | | no TurboID | | Ecm11_TI | | Mer3_TI | | Mlh3_TI | | Msh4_TI | | Msh5_TI | | Spo16_TI | | TI_Gmc2 | | Zip2_TI | | Zip3iTI | | Zip4iTI | | Zip4_cTI | |
|---|---|---|---|---|---|---|---|---|---|---|---|---|---|---|---|---|---|---|---|---|---|---|---|---|---|---|
| Bio View:334 Proteins in 304 Clusters | Alternate ID | olecular Weig | 230113_AM_ | 230113_AM_ | 230113_AM_ | 230113_AM_ | 230113_AM_ | 230113_AM_ | 30113_AM_ | 30113_AM_ | 230113_AM_ | 230113_AM_ | 230113_AM_ | 230113_AM_ | 230113_AM_ | 230113_AM_ | 230113_AM_ | 230113_AM_ | 30113_AM_ | 30113_AM_ | 230113_AM_ | 230113_AM_ | 230113_AM_ | 230113_AM_ | 230113_AM_ | 230113_AM_ |
| Chromosome stability protein 9 OS=S | **Zip3/Cst9** | 54 kDa | 0 | 0 | 8.74E+06 | 1.61E+07 | 1.09E+07 | 8.13E+06 | 0 | 0 | 4.03E+07 | 2.85E+07 | 3.00E+07 | 2.35E+07 | 4.40E+07 | 3.91E+07 | 0 | 0 | 2.98E+07 | 2.69E+07 | 4.15E+07 | 3.52E+07 | 2.73E+07 | 1.73E+07 | 2.51E+07 | 1.74E+07 |
| ATP-dependent DNA helicase MER3 C | **HFM1** | 135 kDa | 0 | 0 | 2.33E+06 | 3.59E+06 | 2.22E+07 | 2.07E+07 | 0 | 0 | 1.04E+07 | 1.46E+07 | 5.64E+06 | 2.56E+06 | 4.18E+06 | 3.36E+06 | 0 | 0 | 7.63E+06 | 7.15E+06 | 1.96E+06 | 0 | 1.28E+07 | 1.10E+07 | 7.14E+06 | 6.98E+06 |
| Protein ECM11 OS=Saccharomyces ce | **ECM11** | 34 kDa | 0 | 0 | 2.79E+07 | 3.09E+07 | 9.16E+06 | 8.71E+06 | 0 | 0 | 1.39E+07 | 1.01E+07 | 6.43E+06 | 4.74E+06 | 7.82E+06 | 7.71E+06 | 4.11E+06 | 4.24E+06 | 9.90E+06 | 1.06E+07 | 1.02E+07 | 9.82E+06 | 4.62E+07 | 3.04E+07 | 3.22E+07 | 2.32E+07 |
| Nucleolar protein NET1 OS=Saccharo | **NET1** | 129 kDa | 0 | 0 | 3.08E+06 | 5.93E+06 | 8.00E+06 | 6.94E+06 | 0 | 1.76E+06 | 8.94E+06 | 1.04E+07 | 7.47E+06 | 4.42E+06 | 3.52E+06 | 2.50E+06 | 0 | 1 | 2.85E+06 | 5.26E+06 | 5.63E+06 | 4.47E+06 | 5.18E+06 | 5.55E+06 | 5.22E+06 | 4.57E+06 |
| Synaptonemal complex protein ZIP1 C | **ZIP1** | 100 kDa | 0 | 0 | 6.84E+06 | 1.01E+07 | 4.54E+06 | 3.37E+06 | 0 | 0 | 4.92E+06 | 4.90E+06 | 2.35E+06 | 1.89E+06 | 3.42E+06 | 2.69E+06 | 0 | 0 | 2.68E+06 | 2.22E+06 | 3.00E+06 | 2.76E+06 | 9.40E+06 | 6.83E+06 | 6.44E+06 | 6.73E+06 |
| Protein ZIP2 OS=Saccharomyces cere | **ZIP2** | 83 kDa | 0 | 0 | 0 | 0 | 0 | 0 | 0 | 0 | 8.35E+06 | 9.78E+06 | 4.26E+06 | 2.30E+06 | 3.45E+06 | 2.34E+06 | 0 | 0 | 1.82E+07 | 2.46E+07 | 2.12E+06 | 0 | 5.85E+06 | 4.94E+06 | 7.96E+06 | 8.28E+06 |
| Protein RED1 OS=Saccharomyces cere | **RED1** | 96 kDa | 0 | 0 | 1.72E+06 | 2.04E+06 | 2.95E+06 | 3.66E+06 | 0 | 0 | 3.70E+06 | 3.37E+06 | 1.42E+06 | 1.04E+06 | 3.32E+06 | 3.61E+06 | 0 | 0 | 6.75E+06 | 8.25E+06 | 0 | 0 | 8.67E+06 | 6.89E+06 | 7.54E+06 | 6.13E+06 |
| Heat shock transcription factor OS=Sa | **HSF1** | 93 kDa | 0 | 0 | 6.18E+06 | 1.28E+07 | 3.51E+06 | 3.20E+06 | 0 | 0 | 3.56E+06 | 3.86E+06 | 3.09E+06 | 2.82E+06 | 2.44E+06 | 2.11E+06 | 1.64E+06 | 0 | 2.51E+06 | 3.47E+06 | 3.24E+06 | 3.53E+06 | 7.33E+06 | 5.58E+06 | 9.52E+06 | 6.07E+06 |
| MutS protein homolog 5 OS=Saccharo | **MSH5** | 102 kDa | 0 | 0 | 0 | 0 | 0 | 0 | 0 | 0 | 8.10E+06 | 7.01E+06 | 4.62E+07 | 3.53E+07 | 1.82E+06 | 1.46E+06 | 0 | 0 | 1.79E+06 | 1.45E+06 | 2.54E+06 | 2.13E+06 | 1.96E+06 | 1.12E+06 | 1.26E+06 | 0 |
| DNA repair protein RAD2 OS=Sacchar | **RAD2** | 118 kDa | 0 | 0 | 1.30E+06 | 3.23E+06 | 2.26E+06 | 2.09E+06 | 0 | 0 | 3.36E+06 | 2.89E+06 | 2.54E+06 | 1.75E+06 | 0 | 0 | 0 | 0 | 0 | 0 | 1.16E+06 | 1.71E+06 | 6.62E+06 | 4.88E+06 | 8.50E+06 | 7.09E+06 |
| ATP-dependent DNA helicase PIF1 OS | **PIF1** | 98 kDa | 0 | 0 | 0 | 0 | 9.58E+06 | 8.79E+06 | 0 | 0 | 2.15E+06 | 3.23E+06 | 0 | 0 | 1.84E+06 | 9.10E+05 | 0 | 0 | 1.57E+06 | 0 | 0 | 0 | 3.95E+06 | 2.90E+06 | 2.36E+06 | 1.60E+06 |
| Centromere-binding protein 1 OS=Sac | **CBF1** | 39 kDa | 0 | 0 | 6.10E+06 | 1.08E+07 | 2.93E+06 | 1.98E+06 | 0 | 0 | 3.90E+06 | 3.50E+06 | 2.60E+06 | 1.93E+06 | 1.00E+06 | 1.57E+06 | 8.67E+05 | 0 | 9.25E+05 | 1.27E+06 | 1.19E+06 | 1.21E+06 | 4.78E+06 | 3.23E+06 | 5.23E+06 | 2.29E+06 |
| Meiotic activator RIM4 OS=Saccharon | **RIM4** | 80 kDa | 0 | 0 | 2.85E+06 | 3.74E+06 | 7.90E+06 | 3.82E+06 | 3.46E+06 | 3.70E+06 | 3.40E+06 | 3.73E+06 | 3.90E+06 | 4.72E+06 | 3.89E+06 | 2.45E+06 | 2.39E+06 | 2.63E+06 | 2.04E+06 | 3.00E+06 | 3.45E+06 | 1.91E+06 | 1.30E+07 | 8.40E+06 | 1.24E+07 | 1.15E+07 |
| Thioredoxin-2 OS=Saccharomyces cer | **TRX2** | 11 kDa | 0 | 0 | 4.57E+06 | 7.53E+06 | 8.86E+06 | 7.36E+06 | 2.76E+06 | 0 | 1.22E+07 | 1.23E+07 | 1.03E+07 | 8.84E+06 | 3.13E+06 | 3.41E+06 | 1.24E+06 | 9.08E+05 | 2.40E+06 | 3.10E+06 | 3.95E+06 | 4.23E+06 | 1.88E+07 | 1.10E+07 | 1.16E+07 | 6.52E+06 |
| Meiosis-specific protein SPO13 OS=Sa | **SPO13** | 33 kDa | 0 | 0 | 3.00E+06 | 3.16E+06 | 6.55E+06 | 3.03E+06 | 0 | 0 | 5.34E+06 | 3.62E+06 | 3.54E+06 | 3.32E+06 | 4.32E+06 | 2.68E+06 | 0 | 0 | 2.45E+06 | 4.56E+06 | 3.76E+06 | 4.75E+06 | 6.03E+06 | 4.51E+06 | 7.15E+06 | 5.49E+06 |
| Securin OS=Saccharomyces cerevisiae | **PDS1** | 42 kDa | 0 | 0 | 4.09E+06 | 5.50E+06 | 3.74E+06 | 5.44E+06 | 0 | 0 | 7.03E+06 | 5.71E+06 | 6.25E+06 | 2.94E+06 | 5.70E+06 | 4.03E+06 | 0 | 0 | 3.21E+06 | 7.54E+06 | 4.11E+06 | 2.18E+06 | 1.03E+07 | 5.57E+06 | 9.63E+06 | 1.09E+07 |
| Protein KRI1 OS=Saccharomyces cere | **KRI1** | 69 kDa | 0 | 0 | 1.50E+06 | 2.41E+06 | 4.81E+06 | 4.16E+06 | 0 | 0 | 5.86E+06 | 4.62E+06 | 4.67E+06 | 4.24E+06 | 0 | 0 | 0 | 0 | 1.87E+07 | 1 | 2.65E+06 | 3.47E+06 | 3.55E+06 | 3.06E+06 | 3.23E+06 | 2.50E+06 |
| DNA mismatch repair protein MLH3 C | **MLH3** | 82 kDa | 0 | 0 | 0 | 0 | 0 | 0 | 6.07E+06 | 6.75E+06 | 0 | 0 | 0 | 0 | 0 | 0 | 0 | 0 | 0 | 0 | 0 | 0 | 0 | 0 | 0 | 0 |
| Cell division control protein 48 OS=Sa | **CDC48** | 92 kDa | 0 | 0 | 0 | 2.81E+06 | 2.12E+06 | 1.69E+06 | 2.03E+06 | 1.74E+06 | 2.29E+06 | 2.40E+06 | 3.15E+06 | 1.52E+06 | 0 | 0 | 2.43E+06 | 1.72E+06 | 0 | 0 | 2.62E+06 | 1.39E+06 | 2.51E+06 | 2.27E+06 | 1.78E+06 | 2.30E+06 |
| Glycolytic genes transcriptional activa | **GCR1** | 88 kDa | 0 | 0 | 3.68E+06 | 6.11E+06 | 0 | 0 | 0 | 0 | 0 | 0 | 0 | 0 | 0 | 0 | 0 | 0 | 0 | 0 | 2.92E+06 | 3.28E+06 | 5.17E+06 | 3.96E+06 | 4.24E+06 | 2.13E+06 |
| Phosphatidylinositol 3-phosphate-bind | **PIB2** | 71 kDa | 0 | 0 | 0 | 0 | 0 | 0 | 0 | 0 | 0 | 0 | 0 | 0 | 0 | 0 | 0 | 0 | 0 | 7.92E+05 | 0 | 0 | 6.51E+06 | 6.83E+06 | 7.06E+06 | 4.18E+06 |
| DNA mismatch repair protein MLH1 C | **MLH1** | 87 kDa | 0 | 0 | 0 | 0 | 2.85E+06 | 3.03E+06 | 4.85E+06 | 5.30E+06 | 0 | 0 | 0 | 0 | 0 | 0 | 0 | 0 | 0 | 0 | 0 | 0 | 0 | 0 | 0 | 0 |
| Sister chromatid cohesion protein 2 O | **SCC2** | 171 kDa | 0 | 0 | 0 | 1.82E+06 | 1.71E+06 | 1.71E+06 | 0 | 0 | 2.88E+06 | 2.05E+06 | 1.09E+06 | 0 | 0 | 0 | 0 | 0 | 0 | 0 | 0 | 0 | 2.20E+06 | 1.63E+06 | 3.04E+06 | 1.95E+06 |
| Cluster of Cell wall mannoprotein HSF | **HSP150** | 41 kDa | 0 | 0 | 9.96E+05 | 0 | 7.95E+06 | 0 | 3.88E+06 | 1.05E+07 | 1.58E+07 | 3.22E+06 | 7.87E+06 | 1.10E+07 | 2.79E+06 | 0 | 0 | 0 | 0 | 0 | 0 | 2.30E+07 | 4.53E+06 | 3.64E+07 | 3.89E+07 | 1.96E+07 |
| Kelch repeat-containing protein 2 OS= | **KEL2** | 100 kDa | 0 | 0 | 0 | 1.47E+06 | 0 | 0 | 0 | 0 | 0 | 0 | 1.97E+06 | 1.16E+06 | 0 | 0 | 0 | 0 | 0 | 0 | 0 | 0 | 2.78E+06 | 1.84E+06 | 2.57E+06 | 2.46E+06 |
| rRNA biogenesis protein RRP5 OS=Sa | **RRP5** | 193 kDa | 0 | 0 | 0 | 0 | 3.89E+06 | 2.99E+06 | 0 | 0 | 0 | 1 | 0 | 0 | 0 | 0 | 0 | 0 | 0 | 0 | 0 | 0 | 0 | 0 | 0 | 0 |
| Cell wall protein CWP1 OS=Saccharon | **CWP1** | 24 kDa | 0 | 0 | 0 | 0 | 0 | 0 | 0 | 0 | 1 | 0 | 4.96E+06 | 3.96E+06 | 0 | 0 | 0 | 0 | 0 | 0 | 0 | 4.43E+06 | 2.65E+06 | 1.76E+07 | 1.44E+07 | 9.49E+06 |
| Zinc finger transcription factor YRR1 ( | **YRR1** | 92 kDa | 0 | 0 | 1.69E+06 | 1.51E+06 | 4.87E+06 | 4.38E+06 | 0 | 0 | 0 | 3.05E+06 | 0 | 0 | 0 | 0 | 0 | 0 | 0 | 0 | 0 | 0 | 1.90E+06 | 1.19E+06 | 4.68E+06 | 4.14E+06 |
| Nucleolar complex protein 14 OS=Sac | **NOP14** | 94 kDa | 0 | 0 | 0 | 2.77E+06 | 2.59E+06 | 2.53E+06 | 0 | 0 | 1.65E+06 | 2.74E+06 | 0 | 0 | 0 | 0 | 0 | 0 | 0 | 0 | 0 | 0 | 2.04E+06 | 1.80E+06 | 2.68E+06 | 2.48E+06 |
| Ubiquitin-like protein SMT3 OS=Sacch | **SMT3** | 12 kDa | 0 | 0 | 1.83E+07 | 2.60E+07 | 6.73E+06 | 4.76E+06 | 0 | 0 | 9.76E+06 | 5.24E+06 | 3.55E+06 | 4.79E+06 | 7.22E+06 | 2.89E+06 | 0 | 0 | 0 | 0 | 6.71E+06 | 6.11E+06 | 5.76E+06 | 6.11E+06 | 5.72E+06 | 4.32E+06 |
| Cop9 signalosome-interactor 1 OS=Sa | **CSI1** | 35 kDa | 0 | 0 | 0 | 0 | 0 | 0 | 0 | 0 | 3.91E+06 | 4.94E+06 | 4.56E+06 | 3.65E+06 | 4.76E+06 | 2.18E+06 | 0 | 0 | 0 | 0 | 0 | 1 | 4.18E+06 | 2.04E+06 | 8.50E+06 | 3.81E+06 |
| Uncharacterized protein MRP8 OS=Sa | **MRP8** | 25 kDa | 0 | 0 | 4.19E+06 | 7.11E+06 | 0 | 0 | 0 | 0 | 1.08E+07 | 6.96E+06 | 5.21E+06 | 3.21E+06 | 1.52E+06 | 2.45E+06 | 0 | 0 | 0 | 0 | 0 | 0 | 2.62E+06 | 1.09E+06 | 0 | 0 |
| Sphingolipid long chain base-responsi | **PIL1** | 38 kDa | 0 | 0 | 1.22E+06 | 1.51E+06 | 0 | 0 | 2.19E+06 | 1 | 0 | 9.43E+05 | 1.07E+06 | 8.27E+05 | 1.01E+06 | 7.95E+05 | 0 | 1.13E+06 | 0 | 0 | 8.75E+05 | 0 | 1.95E+06 | 1.36E+06 | 1.75E+06 | 1.43E+06 |
| AP-2 complex subunit beta OS=Saccha | **APL1** | 80 kDa | 0 | 0 | 0 | 0 | 1.57E+06 | 1.70E+06 | 0 | 0 | 2.84E+06 | 2.55E+06 | 9.02E+05 | 0 | 0 | 0 | 0 | 0 | 0 | 0 | 0 | 0 | 3.59E+06 | 1.82E+06 | 1.20E+06 | 1.60E+06 |
| Glucose-6-phosphate 1-epimerase OS | **YMR099C** | 34 kDa | 0 | 0 | 1.30E+06 | 2.07E+06 | 2.71E+06 | 2.33E+06 | 1.40E+06 | 0 | 3.67E+06 | 3.95E+06 | 2.79E+06 | 1.56E+06 | 1.51E+06 | 7.75E+05 | 7.34E+05 | 0 | 9.36E+05 | 0 | 1.18E+06 | 0 | 4.34E+06 | 2.87E+06 | 2.55E+06 | 1.78E+06 |
| Phosphatidylinositol 3-kinase VPS34 C | **VPS34** | 101 kDa | 0 | 0 | 0 | 0 | 0 | 0 | 0 | 0 | 0 | 0 | 0 | 0 | 0 | 0 | 0 | 0 | 0 | 0 | 0 | 0 | 6.49E+06 | 4.82E+06 | 0 | 0 |
| Thioredoxin-1 OS=Saccharomyces cer | **TRX1** | 11 kDa | 0 | 0 | 5.68E+06 | 9.00E+06 | 1.20E+07 | 8.52E+06 | 4.21E+06 | 0 | 7.48E+06 | 6.43E+06 | 6.34E+06 | 1.06E+07 | 4.30E+06 | 4.72E+06 | 0 | 0 | 0 | 0 | 5.79E+06 | 5.31E+06 | 8.37E+06 | 7.41E+06 | 6.01E+06 | 1.05E+07 |
| Ribonucleoside-diphosphate reductas | **RNR2** | 46 kDa | 0 | 0 | 1.88E+06 | 2.77E+06 | 2.56E+06 | 2.22E+06 | 2.89E+06 | 0 | 3.76E+06 | 2.16E+06 | 2.82E+06 | 1.87E+06 | 1.06E+06 | 8.64E+05 | 1.34E+06 | 0 | 0 | 0 | 2.26E+06 | 1.51E+06 | 4.86E+06 | 3.41E+06 | 1.92E+06 | 2.02E+06 |
| Mitochondrial protein import protein | **YDJ1** | 45 kDa | 0 | 0 | 0 | 3.56E+06 | 2.79E+06 | 0 | 0 | 0 | 0 | 4.14E+06 | 3.81E+06 | 2.27E+06 | 0 | 0 | 0 | 0 | 0 | 0 | 2.72E+06 | 0 | 3.57E+06 | 2.89E+06 | 4.04E+06 | 3.05E+06 |
| Topoisomerase 1-associated factor 2 | **TOF2** | 86 kDa | 0 | 0 | 0 | 1 | 2.70E+06 | 3.01E+06 | 0 | 0 | 2.27E+06 | 2.93E+06 | 2.16E+06 | 1 | 0 | 0 | 0 | 0 | 0 | 0 | 0 | 0 | 1.56E+06 | 0 | 1.93E+06 | 0 |
| Replication factor A protein 1 OS=Sac | **RFA1** | 70 kDa | 0 | 0 | 0 | 0 | 3.59E+06 | 2.62E+06 | 0 | 0 | 4.30E+06 | 2.08E+06 | 0 | 0 | 0 | 0 | 0 | 0 | 0 | 0 | 0 | 0 | 3.54E+06 | 2.50E+06 | 0 | 1 |
| Homocysteine/cysteine synthase OS= | **MET17** | 49 kDa | 0 | 0 | 2.74E+06 | 3.93E+06 | 0 | 2.43E+06 | 3.68E+06 | 0 | 0 | 0 | 2.98E+06 | 1.57E+06 | 0 | 1.50E+06 | 3.58E+06 | 1.87E+06 | 0 | 0 | 0 | 1 | 1.28E+06 | 1.65E+06 | 1 | 8.12E+05 |
| UBA domain-containing protein RUP1 | **RUP1** | 75 kDa | 0 | 0 | 0 | 0 | 0 | 3.13E+06 | 0 | 0 | 4.98E+06 | 3.07E+06 | 4.88E+06 | 3.13E+06 | 0 | 0 | 0 | 0 | 0 | 0 | 0 | 0 | 3.92E+06 | 3.18E+06 | 6.39E+06 | 2.74E+06 |
| Reticulon-like protein 1 OS=Saccharor | **RTN1** | 33 kDa | 0 | 0 | 8.40E+05 | 3.05E+06 | 2.59E+06 | 2.56E+06 | 3.24E+06 | 9.69E+05 | 1.78E+06 | 3.15E+06 | 3.37E+06 | 1.46E+06 | 2.01E+06 | 1.79E+06 | 3.64E+06 | 1.15E+06 | 0 | 0 | 0 | 2.29E+06 | 2.83E+06 | 0 | 6.51E+05 | 5.54E+05 |
| Antiviral helicase SKI2 OS=Saccharom | **SKI2** | 146 kDa | 0 | 0 | 0 | 8.53E+06 | 8.17E+06 | 0 | 0 | 7.50E+06 | 0 | 4.67E+06 | 6.33E+06 | 9.29E+05 | 0 | 0 | 0 | 2.27E+06 | 0 | 0 | 0 | 0 | 2.60E+06 | 1.74E+06 | 4.22E+06 | 1.44E+06 |
| Meiosis-specific protein HOP1 OS=Sa | **HOP1** | 69 kDa | 0 | 0 | 1.70E+06 | 1.76E+06 | 2.35E+06 | 1.26E+06 | 0 | 0 | 0 | 1.65E+06 | 6.42E+05 | 0 | 0 | 0 | 0 | 0 | 0 | 0 | 0 | 0 | 2.27E+06 | 1.11E+06 | 2.39E+06 | 1.26E+06 |
| Filament protein FIN1 OS=Saccharom | **FIN1** | 33 kDa | 0 | 0 | 0 | 0 | 0 | 2.37E+06 | 0 | 0 | 3.54E+06 | 3.15E+06 | 0 | 0 | 0 | 0 | 0 | 0 | 0 | 0 | 0 | 0 | 3.89E+06 | 2.92E+06 | 4.59E+06 | 3.14E+06 |
| Oligoribonuclease, mitochondrial OS= | **REX2** | 31 kDa | 0 | 0 | 9.96E+05 | 0 | 1 | 1 | 0 | 0 | 1.66E+06 | 2.15E+06 | 2.27E+06 | 1.16E+06 | 0 | 0 | 0 | 0 | 0 | 0 | 0 | 0 | 4.49E+06 | 1.41E+06 | 3.12E+06 | 2.44E+06 |
| Stationary phase protein 4 OS=Saccha | **SPG4** | 13 kDa | 0 | 0 | 0 | 0 | 0 | 3.33E+06 | 0 | 0 | 4.85E+06 | 2.89E+06 | 2.72E+06 | 2.24E+06 | 1.30E+06 | 0 | 0 | 0 | 0 | 0 | 0 | 0 | 4.73E+06 | 4.12E+06 | 4.14E+06 | 2.62E+06 |
| 12 kDa heat shock protein OS=Saccha | **HSP12** | 12 kDa | 0 | 0 | 1.82E+06 | 0 | 8.09E+05 | 2.24E+06 | 1.01E+06 | 0 | 2.38E+06 | 1.48E+06 | 2.67E+06 | 0 | 0 | 0 | 0 | 0 | 0 | 0 | 0 | 1.60E+06 | 3.90E+06 | 2.28E+06 | 1.95E+06 | 0 |
| Nuclear GTP-binding protein NUG1 O | **NUG1** | 58 kDa | 0 | 0 | 0 | 0 | 3.00E+06 | 2.96E+06 | 0 | 0 | 0 | 1 | 0 | 0 | 0 | 0 | 0 | 0 | 0 | 0 | 0 | 0 | 2.45E+06 | 1.26E+06 | 3.28E+06 | 1.48E+06 |
| Protoplast secreted protein 2 OS=Sac | **PST2** | 21 kDa | 0 | 0 | 3.49E+06 | 6.11E+06 | 1 | 3.68E+06 | 5.20E+06 | 0 | 5.97E+06 | 5.02E+06 | 4.16E+06 | 2.56E+06 | 1 | 3.64E+06 | 3.75E+06 | 0 | 0 | 0 | 0 | 1 | 6.62E+06 | 4.92E+06 | 5.60E+06 | 5.05E+06 |
| Multiprotein-bridging factor 1 OS=Sac | **MBF1** | 16 kDa | 0 | 0 | 1.24E+06 | 0 | 2.69E+06 | 1.89E+06 | 3.71E+06 | 0 | 4.30E+06 | 4.88E+06 | 2.42E+06 | 1.93E+06 | 1.59E+06 | 1.79E+06 | 0 | 0 | 0 | 0 | 0 | 1.83E+06 | 4.12E+06 | 2.60E+06 | 3.96E+06 | 3.31E+06 |
| 40S ribosomal protein S1-A OS=Sacch | **RPS1A** | 29 kDa | 0 | 0 | 0 | 2.25E+06 | 1.69E+06 | 0 | 4.45E+06 | 1.88E+06 | 0 | 1 | 0 | 1.23E+06 | 0 | 1.96E+06 | 3.16E+06 | 1.93E+06 | 0 | 0 | 2.26E+06 | 1.93E+06 | 1 | 1.23E+06 | 1.34E+06 | 1.07E+06 |
| 27S pre-rRNA (guanosine(2922)-2'-O) | **SPB1** | 96 kDa | 0 | 0 | 0 | 0 | 4.23E+06 | 4.52E+06 | 0 | 0 | 0 | 0 | 0 | 0 | 0 | 0 | 0 | 0 | 0 | 0 | 2.89E+06 | 3.43E+06 | 5.90E+05 | 0 | 0 | 0 |
| Exosome complex exonuclease RRP6 | **RRP6** | 84 kDa | 0 | 0 | 1.46E+06 | 2.96E+06 | 0 | 0 | 0 | 0 | 0 | 0 | 0 | 0 | 0 | 0 | 0 | 0 | 0 | 0 | 0 | 0 | 0 | 0 | 2.27E+06 | 2.26E+06 |
| CTD kinase subunit alpha OS=Sacchar | **CTK1** | 61 kDa | 0 | 0 | 0 | 3.07E+06 | 4.08E+06 | 3.07E+06 | 0 | 0 | 2.53E+06 | 2.22E+06 | 0 | 0 | 0 | 0 | 0 | 0 | 1.64E+06 | 0 | 0 | 0 | 2.85E+06 | 1.80E+06 | 3.06E+06 | 1.49E+06 |
| ATP-dependent helicase SGS1 OS=Sa | **SGS1** | 164 kDa | 0 | 0 | 0 | 0 | 2.91E+06 | 2.01E+06 | 0 | 0 | 0 | 0 | 0 | 0 | 0 | 0 | 0 | 0 | 1 | 0 | 0 | 0 | 1.64E+06 | 1.15E+06 | 1.66E+06 | 0 |
| 1,4-alpha-glucan-branching enzyme C | **GLC3** | 81 kDa | 0 | 0 | 1.47E+06 | 1.58E+06 | 3.37E+06 | 1.14E+06 | 1.38E+06 | 0 | 0 | 1.89E+06 | 2.56E+06 | 1 | 1 | 0 | 1.18E+06 | 1.21E+06 | 0 | 0 | 1.48E+06 | 0 | 1.49E+06 | 1.11E+06 | 0 | 1.16E+06 |
| Protein PET10 OS=Saccharomyces cer | **PET10** | 31 kDa | 0 | 0 | 0 | 0 | 0 | 0 | 1.47E+06 | 0 | 0 | 1.90E+06 | 1.07E+06 | 9.84E+05 | 1.14E+06 | 0 | 0 | 0 | 0 | 0 | 0 | 0 | 1.71E+06 | 1.09E+06 | 1.62E+06 | 1.97E+06 |
| Grand meiotic recombination cluster | **GMC2** | 22 kDa | 0 | 0 | 4.37E+06 | 6.38E+06 | 0 | 0 | 0 | 0 | 3.09E+06 | 2.68E+06 | 2.44E+06 | 1.86E+06 | 0 | 0 | 0 | 0 | 0 | 0 | 0 | 0 | 0 | 0 | 0 | 0 |
| Glycylpeptide N-tetradecanoyltransfe | **NMT1** | 53 kDa | 0 | 0 | 0 | 0 | 0 | 0 | 0 | 0 | 0 | 0 | 0 | 0 | 0 | 0 | 0 | 0 | 0 | 0 | 0 | 0 | 2.83E+06 | 1.35E+06 | 3.46E+06 | 2.77E+06 |
| RNA polymerase II transcription facto | **TFB3** | 38 kDa | 0 | 0 | 0 | 0 | 0 | 0 | 0 | 0 | 2.62E+06 | 3.36E+06 | 4.67E+06 | 3.10E+06 | 0 | 0 | 0 | 0 | 0 | 0 | 0 | 0 | 0 | 0 | 0 | 0 |
| Inner centromere protein-related prot | **SLI15** | 79 kDa | 0 | 0 | 2.28E+06 | 1.24E+06 | 2.43E+06 | 2.49E+06 | 0 | 0 | 3.19E+06 | 2.13E+06 | 2.62E+06 | 0 | 0 | 0 | 0 | 0 | 4.23E+06 | 2.53E+06 | 0 | 0 | 2.49E+06 | 3.42E+06 | 1.86E+06 | 0 |
| Uncharacterized protein YMR147W O | **YMR147W** | 26 kDa | 0 | 0 | 1.42E+06 | 0 | 2.71E+06 | 1 | 0 | 2.07E+06 | 0 | 2.67E+06 | 2.19E+06 | 0 | 0 | 0 | 1.35E+06 | 1.67E+06 | 0 | 0 | 0 | 0 | 0 | 2.38E+06 | 3.63E+06 | 3.61E+06 |
| 60S ribosomal protein L25 OS=Saccha | **RPL25** | 16 kDa | 0 | 0 | 0 | 5.02E+06 | 4.44E+06 | 0 | 4.61E+06 | 1 | 0 | 0 | 1.00E+07 | 4.44E+06 | 1 | 0 | 3.67E+06 | 0 | 0 | 0 | 4.94E+06 | 7.02E+06 | 0 | 0 | 0 | 0 |
| Covalently-linked cell wall protein 14 | **CCW14** | 23 kDa | 0 | 0 | 0 | 0 | 0 | 0 | 0 | 0 | 8.16E+06 | 0 | 0 | 4.18E+06 | 0 | 0 | 0 | 0 | 0 | 0 | 0 | 9.71E+06 | 7.21E+06 | 1.34E+07 | 2.40E+07 | 1.67E+07 |
| SWI/SNF chromatin-remodeling comp | **SNF5** | 103 kDa | 0 | 0 | 2.60E+06 | 1.52E+06 | 0 | 0 | 0 | 0 | 0 | 0 | 0 | 0 | 0 | 0 | 0 | 0 | 0 | 0 | 0 | 0 | 2.92E+06 | 0 | 2.49E+06 | 1.31E+06 |
| Helicase SEN1 OS=Saccharomyces cer | **SEN1** | 253 kDa | 0 | 0 | 0 | 0 | 2.20E+06 | 1.65E+06 | 0 | 0 | 0 | 0 | 4.67E+06 | 0 | 0 | 0 | 0 | 0 | 0 | 0 | 0 | 0 | 0 | 0 | 0 | 0 |
| Transcriptional modulator WTM1 OS= | **WTM1** | 48 kDa | 0 | 0 | 0 | 2.36E+06 | 1.88E+06 | 0 | 1.88E+06 | 0 | 0 | 2.65E+06 | 0 | 0 | 0 | 0 | 0 | 0 | 0 | 0 | 0 | 0 | 4.45E+06 | 2.46E+06 | 1.88E+06 | 8.50E+06 |
| Asparagine-rich zinc finger protein AZ | **AZF1** | 101 kDa | 0 | 0 | 0 | 0 | 2.48E+06 | 2.40E+06 | 0 | 0 | 0 | 3.21E+06 | 0 | 0 | 0 | 0 | 0 | 0 | 0 | 0 | 0 | 0 | 2.71E+06 | 2.52E+06 | 2.30E+06 | 1.39E+06 |
| 1,3-beta-glucanosyltransferase GAS1 | **GAS1** | 60 kDa | 0 | 0 | 0 | 0 | 0 | 0 | 0 | 0 | 0 | 0 | 0 | 1.17E+06 | 0 | 0 | 0 | 0 | 0 | 0 | 0 | 1.50E+06 | 0 | 5.16E+06 | 3.04E+06 | 4.13E+06 |
| ATP synthase subunit alpha, mitochor | **ATP1** | 59 kDa | 0 | 0 | 0 | 0 | 0 | 0 | 2.63E+06 | 0 | 0 | 0 | 0 | 0 | 0 | 0 | 1.54E+06 | 0 | 0 | 0 | 5.27E+05 | 0 | 0 | 0 | 0 | 0 |
| Ubiquitin thioesterase OTU1 OS=Sacc | **OTU1** | 34 kDa | 0 | 0 | 1.43E+06 | 0 | 2.83E+06 | 1.32E+06 | 0 | 0 | 0 | 1.58E+06 | 1.68E+06 | 0 | 0 | 0 | 0 | 0 | 0 | 0 | 5.08E+05 | 9.47E+05 | 2.93E+06 | 2.13E+06 | 1.88E+06 | 0 |
| Nicotinamidase OS=Saccharomyces c | **PNC1** | 25 kDa | 0 | 0 | 0 | 2.14E+06 | 0 | 0 | 5.86E+06 | 0 | 0 | 0 | 3.35E+06 | 1.15E+06 | 2.51E+06 | 0 | 0 | 0 | 0 | 0 | 0 | 0 | 3.62E+06 | 2.17E+06 | 3.11E+06 | 2.86E+06 |
| 5-methyltetrahydropteroyltriglutamat | **MET6** | 86 kDa | 0 | 0 | 0 | 1.25E+06 | 9.04E+05 | 1.03E+06 | 1.33E+06 | 8.96E+05 | 0 | 0 | 0 | 0 | 8.94E+05 | 0 | 0 | 0 | 0 | 0 | 2.08E+06 | 1.23E+06 | 0 | 0 | 1 | 0 |
| 60S ribosomal protein L14-A OS=Sacc | **RPL14A** | 15 kDa | 0 | 0 | 0 | 0 | 4.53E+06 | 0 | 2.98E+06 | 2.44E+06 | 0 | 1 | 0 | 2.42E+06 | 2.28E+06 | 1.96E+06 | 1 | 2.00E+06 | 0 | 0 | 0 | 0 | 0 | 0 | 0 | 0 |
| Transcription elongation factor SPT6 ( | **SPT6** | 168 kDa | 0 | 0 | 0 | 0 | 1.86E+06 | 2.08E+06 | 0 | 0 | 0 | 0 | 0 | 0 | 0 | 0 | 0 | 0 | 0 | 0 | 0 | 0 | 1.68E+06 | 0 | 0 | 0 |
| Glutamine--fructose-6-phosphate am | **GFA1** | 80 kDa | 0 | 0 | 0 | 2.30E+06 | 1.87E+06 | 0 | 3.74E+06 | 3.09E+06 | 0 | 1.69E+06 | 0 | 0 | 1.51E+06 | 1.11E+06 | 1.82E+06 | 1.96E+06 | 0 | 0 | 2.58E+06 | 0 | 0 | 0 | 0 | 0 |
| Hsp90 co-chaperone AHA1 OS=Saccha | **AHA1** | 39 kDa | 0 | 0 | 0 | 0 | 2.54E+06 | 0 | 0 | 1.44E+06 | 9.25E+05 | 3.97E+06 | 2.25E+06 | 0 | 0 | 0 | 1.60E+06 | 0 | 0 | 0 | 3.72E+06 | 0 | 0 | 0 | 0 | 0 |
| Eisosome protein 1 OS=Saccharomyce | **EIS1** | 93 kDa | 0 | 0 | 0 | 0 | 0 | 0 | 0 | 1 | 1 | 0 | 0 | 0 | 0 | 0 | 0 | 1 | 0 | 0 | 0 | 0 | 1.93E+06 | 1.08E+06 | 1.46E+06 | 1.33E+06 |

| | | | | | | | | | | | | | | | | | | | | | | | | | | |
|---|---|---|---|---|---|---|---|---|---|---|---|---|---|---|---|---|---|---|---|---|---|---|---|---|---|---|
| ATP-dependent RNA helicase eIF4A O | **TIF1** | 45 kDa | 0 | 0 | 3.04E+06 | 0 | 1.86E+06 | 1.37E+06 | 3.29E+06 | 1.58E+06 | 0 | 0 | 0 | 0 | 0 | 0 | 2.82E+06 | 0 | 0 | 0 | 0 | 0 | 0 | 0 | 0 | 0 |
| Histone promoter control protein 2 OS | **HPC2** | 68 kDa | 0 | 0 | 0 | 0 | 2.69E+06 | 2.34E+06 | 0 | 0 | 0 | 0 | 0 | 0 | 0 | 0 | 0 | 0 | 0 | 0 | 0 | 0 | 1.76E+06 | 1.31E+06 | 0 | 0 |
| Long-chain-fatty-acid--CoA ligase 2 O | **FAA2** | 83 kDa | 0 | 0 | 0 | 0 | 0 | 0 | 0 | 0 | 0 | 0 | 0 | 0 | 0 | 0 | 0 | 0 | 0 | 0 | 0 | 0 | 0 | 0 | 2.54E+06 | 1.08E+06 |
| Nucleoporin NUP2 OS=Saccharomyce | **NUP2** | 78 kDa | 0 | 0 | 0 | 0 | 0 | 0 | 0 | 0 | 2.72E+06 | 3.74E+06 | 0 | 0 | 0 | 0 | 0 | 0 | 0 | 0 | 0 | 1.23E+06 | 2.65E+06 | 2.62E+06 | 1.72E+06 | 0 |
| NADP-specific glutamate dehydrogen | **GDH3** | 50 kDa | 0 | 0 | 0 | 0 | 0 | 0 | 0 | 0 | 0 | 0 | 0 | 9.40E+05 | 0 | 0 | 0 | 0 | 0 | 0 | 0 | 0 | 2.65E+06 | 2.14E+06 | 1.77E+06 | 1.78E+06 |
| Meiosis induction protein kinase IME2 | **IME2** | 74 kDa | 0 | 0 | 0 | 0 | 0 | 0 | 0 | 0 | 2.77E+06 | 2.95E+06 | 0 | 0 | 0 | 0 | 0 | 0 | 0 | 0 | 3.31E+06 | 2.26E+06 | 0 | 0 | 0 | 0 |
| Arginase OS=Saccharomyces cerevisia | **CAR1** | 36 kDa | 0 | 0 | 1.13E+06 | 0 | 2.29E+06 | 0 | 1.85E+06 | 0 | 0 | 0 | 1.68E+06 | 0 | 0 | 0 | 1.01E+06 | 0 | 0 | 0 | 0 | 0 | 2.04E+06 | 1.18E+06 | 1.90E+06 | 4.32E+06 |
| Elongation factor 1-gamma 1 OS=Sac | **CAM1** | 47 kDa | 0 | 0 | 4.11E+06 | 3.25E+06 | 0 | 0 | 4.43E+06 | 6.72E+06 | 0 | 0 | 0 | 0 | 0 | 0 | 5.82E+06 | 0 | 3.42E+06 | 1 | 2.46E+06 | 0 | 0 | 0 | 0 | 0 |
| Serine/threonine-protein phosphatase | **PSY4** | 51 kDa | 0 | 0 | 1.20E+06 | 1.62E+06 | 0 | 0 | 0 | 0 | 1.63E+06 | 1.82E+06 | 0 | 0 | 0 | 0 | 0 | 0 | 0 | 0 | 0 | 0 | 3.26E+06 | 2.51E+06 | 3.44E+06 | 2.39E+06 |
| Inositol-3-phosphate synthase OS=Sa | **INO1** | 60 kDa | 0 | 0 | 0 | 0 | 0 | 0 | 3.07E+06 | 0 | 0 | 0 | 0 | 0 | 0 | 0 | 0 | 0 | 0 | 0 | 0 | 0 | 3.61E+06 | 2.21E+06 | 2.10E+06 | 3.21E+06 |
| Glucan 1,3-beta-glucosidase OS=Sacc | **BGL2** | 34 kDa | 0 | 0 | 0 | 0 | 0 | 0 | 3.20E+06 | 0 | 0 | 0 | 0 | 0 | 0 | 0 | 0 | 0 | 0 | 0 | 0 | 0 | 2.20E+06 | 2.41E+06 | 1.94E+06 | 1.62E+06 |
| Transcription factor SPN1 OS=Sacchar | **SPN1** | 46 kDa | 0 | 0 | 0 | 1.84E+06 | 0 | 1.31E+06 | 0 | 0 | 0 | 0 | 0 | 0 | 0 | 0 | 0 | 0 | 0 | 0 | 0 | 0 | 1.92E+06 | 2.34E+06 | 0 | 0 |
| Nucleoporin NSP1 OS=Saccharomyces | **NSP1** | 87 kDa | 0 | 0 | 0 | 0 | 0 | 0 | 0 | 0 | 0 | 0 | 0 | 0 | 0 | 0 | 0 | 0 | 0 | 0 | 0 | 8.60E+05 | 1 | 0 | 2.63E+06 | 1.43E+06 |
| Probable glycosidase CRH1 OS=Sacch | **CRH1** | 53 kDa | 0 | 0 | 0 | 0 | 0 | 0 | 0 | 0 | 0 | 0 | 0 | 0 | 0 | 0 | 0 | 0 | 0 | 0 | 0 | 0 | 0 | 4.81E+06 | 8.58E+06 | 2.20E+06 |
| 60S ribosomal protein L5 OS=Sacchar | **RPL5** | 34 kDa | 0 | 0 | 0 | 0 | 0 | 0 | 2.61E+06 | 8.03E+05 | 0 | 0 | 1 | 0 | 2.04E+06 | 0 | 0 | 0 | 0 | 0 | 1 | 0 | 0 | 1 | 1 | 0 |
| Cell wall protein ECM33 OS=Saccharo | **ECM33** | 44 kDa | 0 | 0 | 0 | 0 | 0 | 0 | 0 | 0 | 0 | 0 | 2.25E+06 | 3.09E+06 | 0 | 0 | 0 | 0 | 0 | 0 | 0 | 2.29E+06 | 0 | 6.09E+06 | 6.34E+06 | 3.25E+06 |
| Pre-mRNA-splicing factor SLU7 OS=S | **SLU7** | 45 kDa | 0 | 0 | 0 | 0 | 0 | 0 | 0 | 0 | 2.43E+06 | 0 | 0 | 0 | 0 | 0 | 0 | 0 | 0 | 0 | 0 | 0 | 2.41E+06 | 1.32E+06 | 1.19E+06 | 3.91E+05 |
| Probable ATP-dependent RNA helicas | **ECM16** | 145 kDa | 0 | 0 | 0 | 0 | 1.95E+06 | 1.56E+06 | 0 | 0 | 0 | 0 | 0 | 0 | 0 | 0 | 0 | 0 | 1 | 0 | 0 | 0 | 1.44E+06 | 8.46E+05 | 0 | 0 |
| Sphingolipid long chain base-responsi | **LSP1** | 38 kDa | 0 | 0 | 1.18E+06 | 1.62E+06 | 1.02E+06 | 0 | 2.28E+06 | 0 | 0 | 0 | 0 | 0 | 0 | 0 | 0 | 0 | 0 | 0 | 0 | 0 | 1.70E+06 | 0 | 0 | 0 |
| cAMP-dependent protein kinase regul | **BCY1** | 47 kDa | 0 | 0 | 0 | 0 | 0 | 0 | 0 | 0 | 0 | 0 | 0 | 0 | 0 | 0 | 0 | 0 | 0 | 0 | 0 | 0 | 2.38E+06 | 2.08E+06 | 3.08E+06 | 2.24E+06 |
| Putative mitochondrial carnitine O-ac | **YAT1** | 78 kDa | 0 | 0 | 2.21E+06 | 0 | 0 | 0 | 3.38E+06 | 3.39E+06 | 0 | 0 | 0 | 0 | 0 | 0 | 2.52E+06 | 2.25E+06 | 0 | 0 | 0 | 0 | 0 | 1.89E+06 | 0 | 0 |
| Succinate dehydrogenase [ubiquinone | **SDH1** | 70 kDa | 0 | 0 | 1.55E+07 | 1.89E+06 | 0 | 0 | 1 | 8.68E+05 | 0 | 0 | 0 | 0 | 3.12E+07 | 0 | 1.95E+06 | 0 | 0 | 0 | 0 | 0 | 0 | 0 | 0 | 0 |
| Elongation of fatty acids protein 3 OS | **ELO3** | 39 kDa | 0 | 0 | 0 | 2.88E+06 | 2.11E+06 | 0 | 0 | 1 | 0 | 0 | 0 | 0 | 0 | 0 | 0 | 1.77E+06 | 9.94E+05 | 1 | 0 | 0 | 0 | 0 | 0 | 0 |
| 40S ribosomal protein S18-A OS=Sacc | **RPS18A** | 17 kDa | 0 | 0 | 0 | 2.18E+06 | 1.26E+06 | 0 | 2.90E+06 | 1.66E+06 | 0 | 0 | 0 | 0 | 0 | 0 | 0 | 0 | 0 | 0 | 0 | 1.12E+06 | 0 | 0 | 0 | 0 |
| Chromatin-remodeling complex subur | **IES6** | 19 kDa | 0 | 0 | 0 | 0 | 0 | 0 | 0 | 0 | 3.47E+06 | 0 | 0 | 0 | 0 | 0 | 0 | 0 | 0 | 0 | 0 | 0 | 7.49E+06 | 4.68E+06 | 1.63E+06 | 5.27E+06 |
| Heat shock protein 42 OS=Saccharom | **HSP42** | 43 kDa | 0 | 0 | 0 | 0 | 0 | 0 | 0 | 0 | 0 | 0 | 0 | 0 | 0 | 0 | 0 | 0 | 0 | 0 | 4.49E+06 | 4.93E+06 | 3.08E+06 | 0 | 3.87E+06 | 0 |
| Clustered mitochondria protein 1 OS= | **CLU1** | 145 kDa | 0 | 0 | 0 | 0 | 0 | 0 | 0 | 0 | 0 | 0 | 1 | 0 | 0 | 0 | 0 | 0 | 0 | 0 | 0 | 0 | 0 | 0 | 1.63E+06 | 1.26E+06 |
| Protein BOI2 OS=Saccharomyces cere | **BOI2** | 116 kDa | 0 | 0 | 1.08E+07 | 0 | 0 | 1.86E+07 | 0 | 0 | 0 | 0 | 0 | 0 | 0 | 0 | 0 | 0 | 0 | 0 | 0 | 0 | 1.92E+06 | 1 | 1.87E+06 | 1.78E+06 |
| 60S ribosomal protein L28 OS=Saccha | **RPL28** | 17 kDa | 0 | 0 | 0 | 0 | 0 | 0 | 5.04E+06 | 0 | 0 | 3.44E+06 | 0 | 0 | 0 | 0 | 2.66E+06 | 0 | 0 | 3.55E+06 | 0 | 2.40E+06 | 0 | 0 | 0 | 0 |
| Fatty acid synthase subunit alpha OS= | **FAS2** | 207 kDa | 0 | 0 | 0 | 9.11E+05 | 0 | 0 | 2.24E+06 | 0 | 0 | 0 | 0 | 0 | 0 | 0 | 1.38E+06 | 0 | 0 | 0 | 1.44E+06 | 0 | 0 | 0 | 0 | 0 |
| Meiosis-inducing protein 1 OS=Saccha | **IME1** | 42 kDa | 0 | 0 | 0 | 0 | 0 | 0 | 0 | 0 | 0 | 0 | 0 | 0 | 0 | 0 | 0 | 0 | 0 | 0 | 3.91E+06 | 1.24E+06 | 0 | 0 | 3.24E+06 | 1 |
| Carnitine O-acetyltransferase, mitoch | **CAT2** | 77 kDa | 0 | 0 | 0 | 0 | 0 | 0 | 0 | 0 | 0 | 0 | 0 | 0 | 0 | 0 | 0 | 0 | 0 | 0 | 0 | 0 | 1.75E+06 | 0 | 2.15E+06 | 2.14E+06 |
| Kinetochore protein SLK19 OS=Saccha | **SLK19** | 95 kDa | 0 | 0 | 0 | 0 | 0 | 0 | 0 | 0 | 9.22E+05 | 0 | 0 | 0 | 0 | 0 | 0 | 0 | 0 | 0 | 0 | 0 | 4.97E+06 | 1.25E+06 | 0 | 0 |
| Malate dehydrogenase, peroxisomal ( | **MDH3** | 37 kDa | 0 | 0 | 0 | 0 | 0 | 0 | 0 | 0 | 0 | 0 | 0 | 0 | 0 | 0 | 0 | 0 | 0 | 0 | 0 | 0 | 1.79E+06 | 1.73E+06 | 1.90E+06 | 0 |
| Sporulation-specific protein 16 OS=Sa | **SPO16** | 23 kDa | 0 | 0 | 0 | 0 | 0 | 0 | 0 | 0 | 0 | 0 | 0 | 0 | 9.37E+06 | 1.49E+07 | 0 | 0 | 0 | 0 | 0 | 0 | 0 | 0 | 0 | 0 |
| Triacylglycerol lipase 4 OS=Saccharon | **TGL4** | 103 kDa | 0 | 0 | 0 | 0 | 0 | 0 | 0 | 0 | 0 | 0 | 0 | 0 | 0 | 0 | 0 | 0 | 0 | 0 | 2.60E+06 | 1.18E+06 | 0 | 0 | 0 | 0 |
| DASH complex subunit DUO1 OS=Sac | **DUO1** | 27 kDa | 0 | 0 | 2.82E+06 | 4.87E+06 | 0 | 0 | 0 | 0 | 0 | 0 | 0 | 0 | 0 | 0 | 0 | 0 | 0 | 0 | 0 | 0 | 0 | 0 | 0 | 0 |
| D-3-phosphoglycerate dehydrogenase | **SER3** | 51 kDa | 0 | 0 | 0 | 0 | 3.55E+06 | 0 | 2.69E+06 | 0 | 2.19E+07 | 0 | 0 | 0 | 6.77E+06 | 0 | 0 | 0 | 0 | 0 | 0 | 0 | 0 | 0 | 6.86E+06 | 0 |
| 40S ribosomal protein S20 OS=Saccha | **RPS20** | 14 kDa | 0 | 0 | 1 | 7.12E+06 | 1 | 0 | 3.80E+06 | 0 | 0 | 0 | 0 | 0 | 0 | 0 | 0 | 0 | 0 | 0 | 2.19E+06 | 0 | 0 | 0 | 0 | 0 |
| Tyrosine-protein phosphatase CDC14 ( | **CDC14** | 62 kDa | 0 | 0 | 0 | 0 | 1.94E+06 | 9.45E+05 | 0 | 0 | 0 | 3.25E+06 | 0 | 0 | 0 | 0 | 0 | 0 | 0 | 0 | 0 | 0 | 3.58E+05 | 0 | 0 | 0 |
| Probable 1,3-beta-glucanosyltransfera | **GAS3** | 57 kDa | 0 | 0 | 0 | 0 | 0 | 0 | 0 | 0 | 0 | 0 | 0 | 0 | 0 | 0 | 0 | 0 | 0 | 0 | 0 | 0 | 0 | 1.82E+06 | 1.94E+06 | 0 |
| RNA polymerase II-associated protein | **PAF1** | 52 kDa | 0 | 0 | 0 | 0 | 2.82E+06 | 2.27E+06 | 0 | 0 | 0 | 0 | 0 | 0 | 0 | 0 | 0 | 0 | 0 | 0 | 0 | 0 | 0 | 0 | 0 | 0 |
| Peptidyl-prolyl cis-trans isomerase OS | **CPR1** | 17 kDa | 0 | 0 | 0 | 0 | 0 | 0 | 2.27E+06 | 0 | 0 | 0 | 0 | 0 | 0 | 0 | 0 | 0 | 0 | 0 | 0 | 0 | 0 | 1 | 0 | 0 |
| Uncharacterized protein YNL195C OS= | **YNL195C** | 28 kDa | 0 | 0 | 0 | 0 | 0 | 3.77E+05 | 0 | 0 | 0 | 0 | 0 | 0 | 0 | 0 | 0 | 0 | 0 | 0 | 0 | 0 | 8.86E+05 | 0 | 9.19E+05 | 5.14E+05 |
| GTPase-activating protein GYP7 OS=S | **GYP7** | 87 kDa | 0 | 0 | 0 | 0 | 0 | 8.76E+05 | 0 | 0 | 2.65E+06 | 0 | 0 | 0 | 0 | 0 | 0 | 0 | 0 | 0 | 0 | 0 | 4.03E+06 | 1.60E+06 | 0 | 0 |
| Aconitate hydratase, mitochondrial O | **ACO1** | 85 kDa | 0 | 0 | 0 | 0 | 0 | 0 | 1.47E+06 | 0 | 0 | 0 | 0 | 0 | 0 | 0 | 1.14E+06 | 0 | 0 | 0 | 0 | 1.45E+06 | 0 | 0 | 0 | 6.06E+05 |
| Potassium-activated aldehyde dehydr | **ALD4** | 57 kDa | 0 | 0 | 1.11E+06 | 0 | 0 | 0 | 1.62E+06 | 0 | 0 | 0 | 0 | 0 | 0 | 0 | 0 | 0 | 0 | 0 | 1.29E+06 | 0 | 0 | 8.47E+05 | 0 | 0 |
| Flavoprotein-like protein YCP4 OS=Sa | **YCP4** | 26 kDa | 0 | 0 | 1 | 0 | 0 | 0 | 2.61E+06 | 0 | 0 | 0 | 0 | 0 | 0 | 0 | 1.40E+06 | 0 | 0 | 0 | 0 | 0 | 0 | 0 | 0 | 0 |
| MAU2 chromatid cohesion factor hon | **SCC4** | 72 kDa | 0 | 0 | 0 | 0 | 0 | 0 | 0 | 0 | 0 | 0 | 0 | 0 | 0 | 0 | 3.66E+06 | 0 | 8.39E+06 | 0 | 0 | 0 | 0 | 0 | 4.93E+07 | 0 |
| Probable family 17 glucosidase SCW4 | **SCW4** | 40 kDa | 0 | 0 | 0 | 0 | 0 | 0 | 0 | 0 | 0 | 0 | 0 | 0 | 0 | 0 | 0 | 0 | 0 | 0 | 0 | 0 | 0 | 3.84E+06 | 2.72E+06 | 2.17E+06 |
| 26S proteasome regulatory subunit 4 | **RPT2** | 49 kDa | 0 | 0 | 0 | 0 | 0 | 0 | 0 | 0 | 0 | 0 | 0 | 0 | 0 | 0 | 0 | 0 | 0 | 0 | 0 | 0 | 0 | 0 | 1.61E+06 | 1.32E+06 |
| ATP synthase subunit beta, mitochond | **ATP2** | 55 kDa | 0 | 0 | 0 | 0 | 0 | 0 | 1.67E+06 | 0 | 0 | 0 | 0 | 0 | 0 | 0 | 0 | 0 | 0 | 0 | 0 | 0 | 0 | 0 | 0 | 0 |
| Structure-specific endonuclease subu | **SLX4** | 84 kDa | 0 | 0 | 0 | 0 | 0 | 0 | 0 | 0 | 0 | 0 | 0 | 0 | 0 | 0 | 0 | 0 | 0 | 0 | 0 | 0 | 1 | 0 | 2.68E+06 | 1 |
| ATP-dependent RNA helicase DBP10 ( | **DBP10** | 113 kDa | 0 | 0 | 0 | 0 | 2.26E+06 | 2.53E+06 | 0 | 0 | 0 | 0 | 0 | 0 | 0 | 0 | 0 | 0 | 0 | 0 | 0 | 0 | 0 | 0 | 0 | 0 |
| DDK kinase regulatory subunit DBF4 C | **DBF4** | 81 kDa | 0 | 0 | 0 | 0 | 1.23E+06 | 2.17E+06 | 0 | 0 | 0 | 0 | 0 | 0 | 0 | 0 | 0 | 0 | 0 | 0 | 0 | 0 | 0 | 0 | 0 | 0 |
| Protein ZEO1 OS=Saccharomyces cere | **ZEO1** | 13 kDa | 0 | 0 | 0 | 0 | 0 | 0 | 1.80E+06 | 0 | 0 | 0 | 0 | 0 | 0 | 0 | 0 | 0 | 0 | 0 | 0 | 0 | 8.95E+05 | 0 | 0 | 0 |
| DASH complex subunit DAM1 OS=Sac | **DAM1** | 38 kDa | 0 | 0 | 8.09E+06 | 1.11E+07 | 0 | 0 | 0 | 0 | 0 | 0 | 0 | 0 | 0 | 0 | 0 | 0 | 0 | 0 | 0 | 0 | 0 | 0 | 0 | 0 |
| 1-phosphatidylinositol 3-phosphate 5- | **FAB1** | 257 kDa | 0 | 0 | 0 | 0 | 0 | 0 | 0 | 0 | 0 | 0 | 0 | 0 | 0 | 0 | 0 | 0 | 2.46E+06 | 2.97E+06 | 0 | 0 | 0 | 0 | 0 | 0 |
| SWI/SNF complex subunit SWI3 OS=S | **SWI3** | 93 kDa | 0 | 0 | 0 | 2.83E+06 | 0 | 0 | 0 | 0 | 0 | 0 | 0 | 0 | 0 | 0 | 0 | 0 | 0 | 0 | 0 | 0 | 0 | 0 | 2.68E+06 | 0 |
| Exosome complex protein LRP1 OS=S | **LRP1** | 21 kDa | 0 | 0 | 2.37E+06 | 2.30E+06 | 0 | 0 | 0 | 0 | 0 | 0 | 0 | 0 | 0 | 0 | 0 | 0 | 0 | 0 | 0 | 0 | 0 | 0 | 0 | 0 |
| 60S ribosomal protein L13-B OS=Sacc | **RPL13B** | 23 kDa | 0 | 0 | 0 | 0 | 0 | 0 | 3.29E+06 | 1.55E+06 | 0 | 0 | 0 | 0 | 0 | 0 | 0 | 0 | 0 | 0 | 0 | 0 | 0 | 0 | 0 | 0 |
| 60S acidic ribosomal protein P0 OS=S | **RPP0** | 34 kDa | 0 | 0 | 0 | 0 | 0 | 0 | 2.84E+06 | 0 | 0 | 0 | 0 | 0 | 0 | 0 | 0 | 0 | 0 | 0 | 0 | 0 | 0 | 0 | 0 | 0 |
| 40S ribosomal protein S13 OS=Saccha | **RPS13** | 17 kDa | 0 | 0 | 0 | 0 | 0 | 0 | 2.74E+06 | 0 | 0 | 0 | 0 | 0 | 0 | 0 | 0 | 0 | 0 | 0 | 0 | 0 | 0 | 0 | 0 | 0 |
| Malate dehydrogenase, mitochondrial | **MDH1** | 36 kDa | 0 | 0 | 0 | 0 | 0 | 0 | 2.48E+06 | 0 | 0 | 0 | 0 | 0 | 0 | 0 | 0 | 0 | 0 | 0 | 0 | 0 | 0 | 0 | 0 | 0 |
| Isocitrate dehydrogenase [NAD] subu | **IDH2** | 40 kDa | 0 | 0 | 0 | 0 | 0 | 0 | 1.76E+06 | 0 | 0 | 0 | 0 | 0 | 0 | 0 | 0 | 0 | 0 | 0 | 0 | 0 | 0 | 0 | 0 | 0 |
| Protein GLC8 OS=Saccharomyces cere | **GLC8** | 27 kDa | 0 | 0 | 0 | 0 | 0 | 0 | 0 | 0 | 0 | 0 | 0 | 0 | 0 | 0 | 0 | 0 | 0 | 0 | 0 | 0 | 0 | 0 | 1.77E+06 | 0 |

**Table S3. Streptavidin-purified proteins from 11 *TurboID* fusion strains identified using UPLC-MS/MS.** File displays the average precursor intensity data corresponding to *S. cerevisiae* proteins detected in streptavidin pull down samples from either replicate of any *TurboID* fusion, and that were also not detected in either of the “no TurboID” replicates.
